# Epigenomic profiling of primate LCLs reveals the evolutionary patterns of epigenetic activities in gene regulatory architectures

**DOI:** 10.1101/2019.12.18.872531

**Authors:** Raquel García-Pérez, Paula Esteller-Cucala, Glòria Mas, Irene Lobón, Valerio Di Carlo, Meritxell Riera, Martin Kuhlwilm, Arcadi Navarro, Antoine Blancher, Luciano Di Croce, José Luis Gómez-Skarmeta, David Juan, Tomàs Marquès-Bonet

**Affiliations:** Institute of Evolutionary Biology (UPF-CSIC), PRBB, Barcelona, Spain; Centre for Genomic Regulation (CRG), The Barcelona Institute of Science and Technology, Spain; Universitat Pompeu Fabra (UPF), Barcelona, Spain; National Institute for Bioinformatics (INB), PRBB, Barcelona, Spain; Institució Catalana de Recerca i Estudis Avançats (ICREA), Barcelona, Spain; Laboratoire d’immunologie, CHU de Toulouse, Institut Fédératif de Biologie, hôpital Purpan, Toulouse, France; Centre de Physiopathologie Toulouse-Purpan (CPTP), Université de Toulouse, Centre National de la Recherche Scientifique (CNRS), Institut National de la Santé et de la Recherche Médicale (Inserm), Université Paul Sabatier (UPS), Toulouse, France; Centro Andaluz de Biología del Desarrollo (CABD), Consejo Superior de Investigaciones Científicas-Universidad Pablo de Olavide-Junta de Andalucía, Seville, Spain; CNAG-CRG, Centre for Genomic Regulation (CRG), Barcelona Institute of Science and Technology (BIST), Barcelona, Spain; Institut Català de Paleontologia Miquel Crusafont, Universitat Autònoma de Barcelona, Cerdanyola del Vallès, Barcelona, Spain

**Author notes:** Contributed equally to this work (Raquel García-Pérez, Paula Esteller-Cucala, David Juan, Tomàs Marquès-Bonet). Corresponding author. (T.M.-B); (D.J.); (R.G.-P.).

**Keywords:** Epigenomics, gene regulation, evolution, positive selection

## Abstract

To gain insight into the evolution of the epigenetic regulation of gene expression in primates, we extensively profiled a new panel of human, chimpanzee, gorilla, orangutan, and macaque lymphoblastoid cell lines (LCLs), using ChIP-seq for five histone marks, ATAC-seq and RNA-seq, further complemented with WGS and WGBS. We annotated regulatory elements and integrated chromatin contact maps to define gene regulatory architectures, creating the largest catalog of regulatory elements in primates to date. We report that epigenetic conservation and its correlation with sequence conservation in primates depends on the activity state of the regulatory element. Our gene regulatory architectures reveal the coordination of different types of components and highlight the role of promoters and intragenic enhancers in the regulation of gene expression. We observed that most regulatory changes occur in weakly active intragenic enhancers. Remarkably, novel human-specific intragenic enhancers with weak activities are enriched in human-specific mutations. These elements appear in genes with signals of positive selection, tissue-specific expression and particular functional enrichments, suggesting that the regulatory evolution of these genes may have contributed to human adaptation.

## Introduction

Changes in chromatin structure and gene regulation play a crucial role in evolution^1,2^. Gene expression differences have been extensively studied in a variety of species and conditions^3–6^. However, there is still much unknown about how regulatory landscapes evolve, even in closely related species. Previous work has focused on the dynamics of the addition and removal of regulatory elements with signals of strong activity during mammalian evolution ‒mainly defined from ChIP-seq experiments on a few histone marks^7–10^. These analyses suggested that enhancers evolve faster than promoters^8,11^. The number of active enhancers located near a gene ‒its regulatory complexity ‒has also been reported to influence the conservation of gene expression in mammals^9^.

Moreover, in a selected group of primates ‒mostly chimpanzees and macaques– changes in histone mark enrichments are associated with gene expression differences^12^. Several studies have also targeted the appearance of human-specific methylation patterns^13,14^ and active promoters and enhancers in different anatomical structures and cell types^8,10^. All these studies have proven that comparative epigenomics is a powerful tool to investigate the evolution of regulatory elements^15,16^. However, a deeper understanding of the evolution of gene regulation requires the integration of multi-layered epigenome data. Only such integration can provide the necessary resolution of regulatory activities for investigating recent evolutionary time frames, as is the case within the primate lineage. Here, we provide an in-depth comparison of the recent evolution of gene regulatory architectures using a homologous cellular model system in human and non-human primates.

## Results

### Comprehensive profiling of primate lymphoblastoid cell lines (LCLs)

We have extensively characterized a panel of lymphoblastoid cell lines (LCLs) from human, chimpanzee, gorilla, orangutan, and macaque, including two independent biological replicates for each species. This characterization includes chromatin immunoprecipitation data (ChIP-seq) from five key histone modifications (H3K4me1, H3K4me3, H3K36me3, H3K27ac, and H3K27me3) and deep-transcriptome sequencing (RNA-seq) (Fig. 1). We integrate these datasets into gene regulatory architectures (Fig. 2a and Supplementary Figs. 1 and 2) to (1) understand how primate gene expression levels are controlled and how expression changes between species occur and to (2) study patterns of evolutionary conservation of regulatory elements in primates. To complement this resource, we have also processed high coverage whole-genome and whole-genome bisulfite sequencing data, as well as chromatin accessibility data (Supplementary Tables 1-10 and Additional files 1-5). Taken together, this is the most extensive collection of great apes and macaque transcriptomic and epigenomic data to date.

**Figure. 1.**
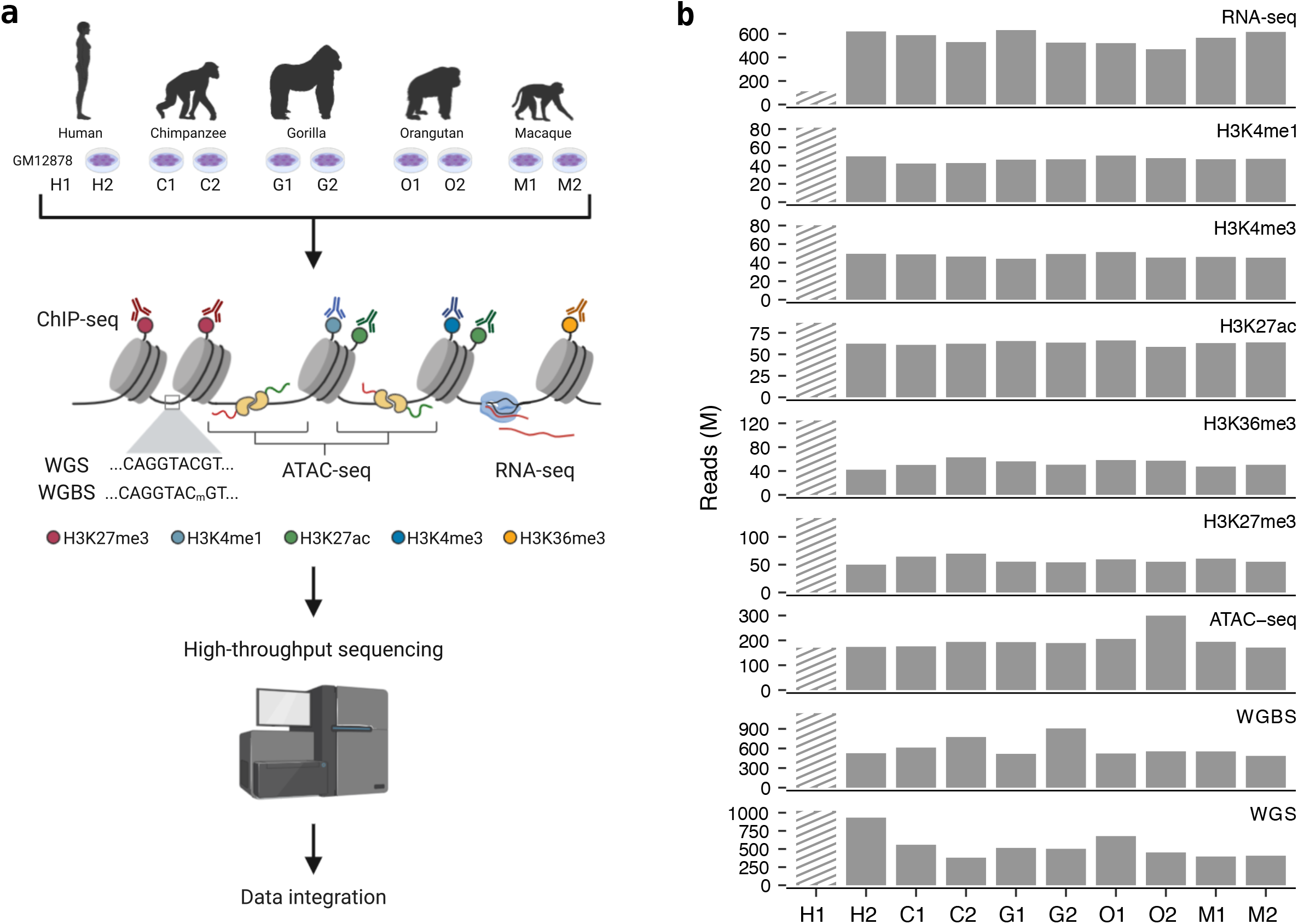
Overview of the study design and data generated. **a,** One human and eight non-human primate lymphoblastoid cell lines (LCLs) were cultured to perform a variety of high-throughput techniques including whole genome sequencing (WGS), whole genome bisulfite sequencing (WGBS), chromatin-accessibility sequencing (ATAC-seq), chromatin immunoprecipitation sequencing (ChIP-seq) targeting five different histone modifications (H3K27me3, H3K4me1, H3K27ac, H3K4me3 and H3K36me3) and transcriptome sequencing (RNA-seq). We integrated previously published datasets from an extensively profiled human LCL (GM12878) to balance the number of human samples (Supplementary Methods). **b,** Number of sequencing reads generated per sample and experiment. Striped lines indicate data retrieved from previously published experiments^81,82^.

**Figure 2.**
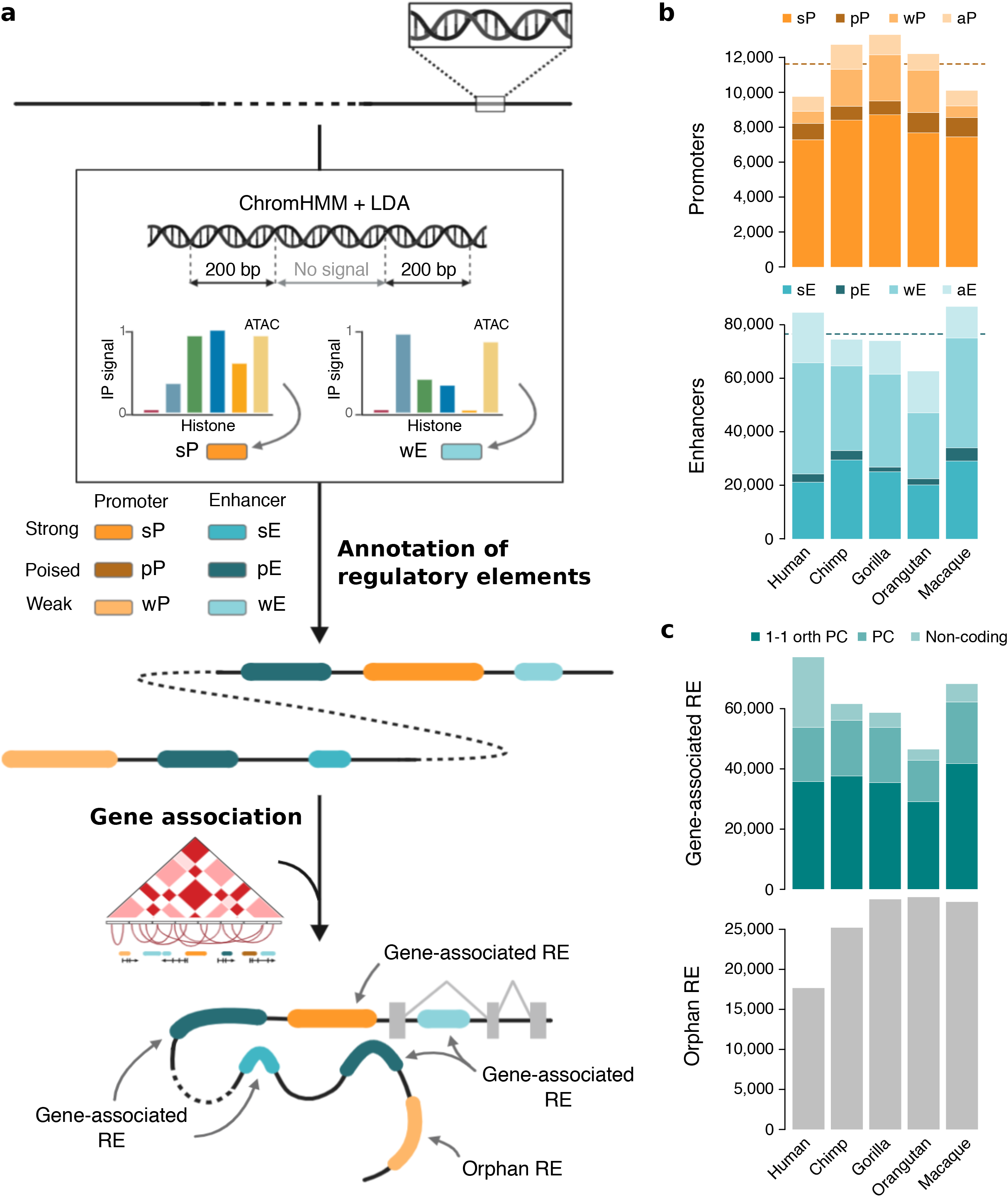
Epigenetic and regulatory characterization of regulatory elements annotated in primates. **a,** Approach followed to annotate and classify regulatory elements (RE). In short, promoter and enhancer states with three activity levels (strong, poised or weak) were annotated for DNA regions based on a combination of chromatin marks and ATAC-seq signals. Regulatory elements (RE) were then linked to genes based on 1D gene proximity and 3D published chromatin maps for LCLs. RE not associated with any gene are referred to as orphan RE. Extended representation in Supplementary Fig. 1. **b,** Number of regulatory elements with promoter and enhancer epigenetic states in each species. **c,** Number of regulatory elements associated with genes and orphan regulatory elements in each species. Genes are divided in 1-to-1 orthologous protein-coding (1-1 orth PC), protein-coding (PC) and non-protein-coding genes. Dashed lines in **b**, and **c**, indicates the average number of RE with promoter and enhancer states annotated across species.

### Annotation of regulatory elements

We used the signal of the ChIP-seq experiments from the five histone marks to identify regulatory regions with characteristic marks of promoters or enhancers (Supplementary Figs. 1 and 2). We defined regulatory regions for each cell line as those containing chromatin states (overrepresented combinations of histone marks detected by ChromHMM^17^) enriched in any regulatory-related histone mark (Fig. 2a and Supplementary Fig. 1). We merged overlapping regulatory regions in the two replicates of every species to define species regulatory elements.

We classified the chromatin states of the regulatory elements based on a hierarchy of functionally interpretable epigenetic states. This hierarchy differentiates chromatin states into promoter (P) and enhancer (E) states, with three different levels of activity each: strong (s), poised (p), or weak (w) (Methods and Supplementary Fig. 1). We improved these assignments by applying a linear discriminative analysis (LDA) with normalized histone and open chromatin enrichments (Methods and Supplementary Figs. 3 and 4). The refined classification results in more similar regulatory landscapes between biological replicates (Wilcoxon signed rank-test: *P* < 0.05 in all species; Supplementary Figs. 5 and 6), with more regulatory elements with the same state in all species (Wilcoxon signed rank-test: *P* = 0.03; Supplementary Figs. 7 and 8).

On average, we found ~11,000 and ~76,000 regulatory elements with promoter and enhancer states per species, respectively (Fig. 2b), of which 69% and 33% are strong, 8% and 4% are poised, and 14% and 45% are weak, respectively (Supplementary Fig. 9 and Supplementary Table 1). Strong and poised activities are more associated with promoter states, whereas weak activities are more frequently associated with enhancer states (Chi-square test: *P* < 2.2 × 10^−16^ in all species; Supplementary Fig. 10). We associated regulatory elements with genes using 1D gene proximity and existing high-resolution 3D chromatin contact data for one of the human LCLs (Fig. 2a and Methods). On average, 70% of the regulatory elements are associated with genes, of which 93% are protein-coding and 61% are 1-to-1 orthologous protein-coding genes in all primate species (Fig. 2c). The set of regulatory elements associated with a gene defines its regulatory architecture.

Altogether, this catalog of regulatory elements provides a comprehensive view of the regulatory landscape of LCLs in humans and non-human primates. In contrast to other commonly used definitions of promoters and enhancers limited to strongly active regions, our multi-layered integration approach allows the additional annotation of weak and poised activities^7,8^. These activities are of particular relevance to improve the definition of elements in regulatory gene architectures. In sum, a detailed primate regulatory catalog enables the study of the evolution of these regulatory activities using LCLs as a proxy of their regulatory potential in other cell types or conditions.

### The evolutionary dynamics of promoters and enhancers in primate LCLs recapitulate previous observations in more distant mammals

Inter-species differences in regulatory regions can be associated with genomic or epigenetic changes. Inconsistencies in the quality of genome assemblies make it difficult to distinguish actual inter-species genomic differences, an issue aggravated in multi-species comparisons. To overcome this problem, we restricted our analyses to unambiguous 1-to-1 orthologs between all species. We detected 28,703 1-to-1 orthologous genomic regions in the five species with a promoter or enhancer state in at least one species (Supplementary Fig. 11). Most of these orthologous regulatory regions (~76%, Binomial test: *P* < 2.2 × 10^−16^) are associated with genes (Methods). In downstream analyses, we focused on these regions integrating the regulatory architectures of protein-coding and non-coding genes.

We quantified the conservation of epigenetic states in regulatory regions as the number of primate species with the epigenetic state in the orthologous regions. In the regulatory architectures of protein-coding genes, promoter states are more conserved than enhancer states (Supplementary Figs. 12-14), with 73% and 60% of regions with a promoter or enhancer state being fully conserved across primates, respectively (Fisher’s exact test: *P* < 2.2 × 10^−16^, *OR* = 1.84; Supplementary Fig. 13). Less than 14% and 8% of orthologous regulatory regions with a promoter or enhancer state are specific to a primate species, respectively (Supplementary Fig. 13). These results for protein-coding genes fall in line with the higher conservation of promoters previously observed in mammals^7^. In contrast, for non-coding genes, promoter states are less conserved than enhancer states (Fisher’s exact test: *P* < 2.2 × 10^−16^, *OR* = 0.39; Supplementary Fig. 14), with 46% and 69% of fully conserved and 26% and 3% of species-specific elements, respectively.

Intrigued by the different epigenetic conservation patterns in protein-coding and non-coding genes, we studied the repurposing and acquisition of regulatory elements. We defined *recently repurposed promoters* ‒or *enhancers*– as regulatory regions with a promoter state in only one species and enhancer states in the remaining species ‒or vice versa. Similarly, *recently gained promoters* or *enhancers* are those regions with a promoter or enhancer state in one species and without regulatory states in any other species.

In agreement with previous studies in more distant species^18^, nearly all (93%) recently evolved promoter states are acquired through repurposing events, whereas the majority (90%) of recently evolved enhancer states are gained (Chi-square test: *P* < 2.2 × 10^−16^; Methods and Supplementary Figs. 15 and 16). The regulatory architectures of protein-coding and non-coding genes ‒the latter evaluated in human due to underrepresentation of non-coding gene annotations in non-human species– show this same pattern (Fisher’s exact test: *P* < 2.2 × 10^−16^, *OR* = Inf, and *P* = 6.2 × 10^−16^, *OR* = 138 respectively; Supplementary Fig. 15).

Our findings confirm those in more distant species^7,18^ and reinforce the generality of these evolutionary dynamics in protein-coding genes. The acquisition of regulatory states in the regulatory architectures of non-coding genes resembles that of protein-coding genes. However, the lower conservation of promoter states associated with non-coding genes suggests that their overall higher conservation is not an intrinsic characteristic of promoter states and that it depends on their specific regulatory relevance in different genes.

### The activity of promoter and enhancers influences their epigenetic and sequence conservation

Taking advantage of our classification of promoters and enhancers into three different activities (strong, poised, and weak), we further explored the patterns of evolutionary conservation of the different regulatory states. Globally, orthologous regulatory regions conserve their regulatory state (Randomization analyses: 1,000 simulations, *P* < 0.05; Supplementary Figs. 17-19 and Supplementary Table 11), but different promoter and enhancer activities show characteristic patterns of conservation (Kruskal-Wallis test: *P* < 2.2 × 10^−16^; Fig. 3a and Supplementary Figs. 20-22).

**Figure 3.**
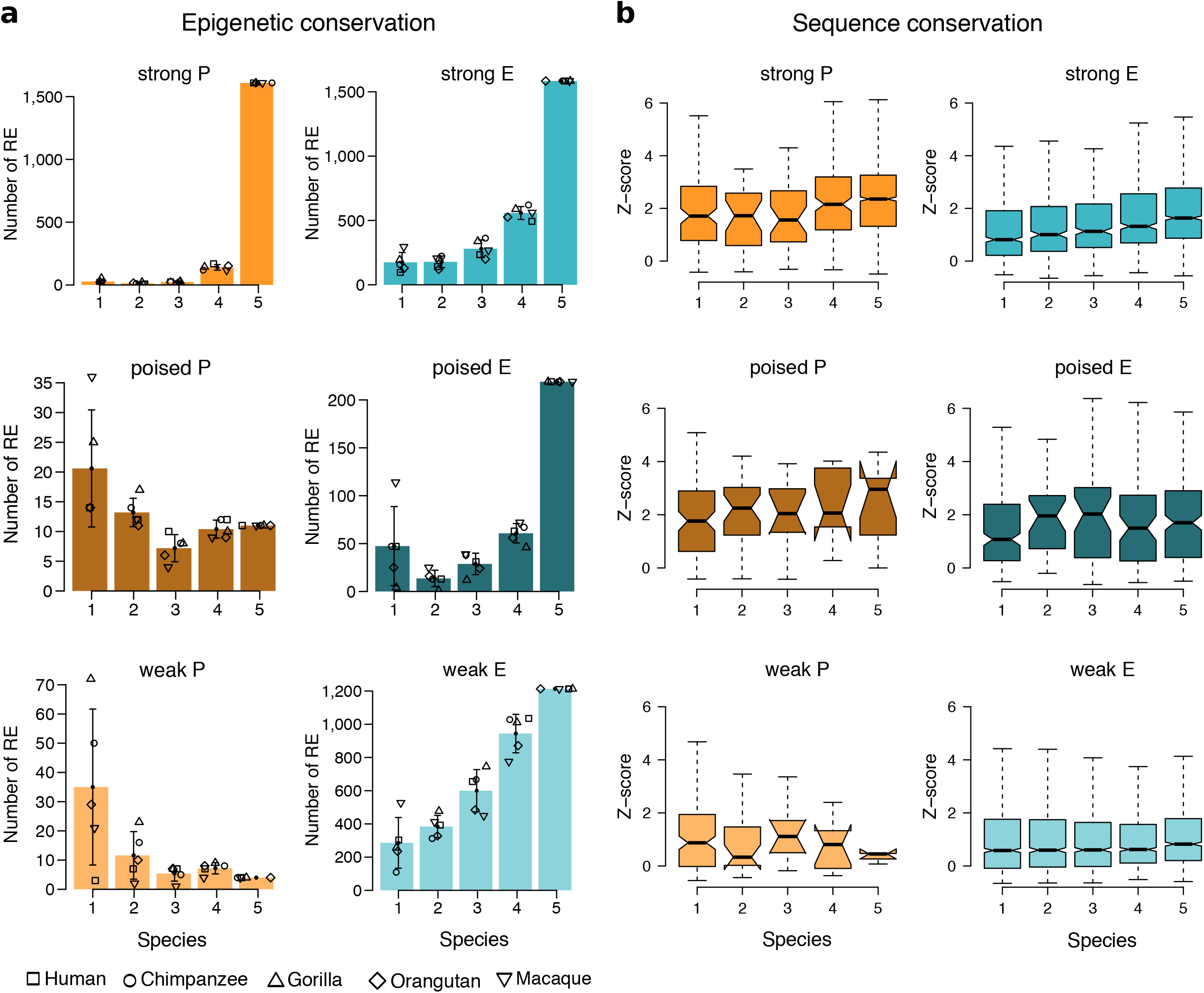
Different regulatory activities have different patterns of epigenetic and sequence conservation. **a,** Barplots show the average number of orthologous regulatory regions across species with the corresponding color-coded epigenetic state conserved in 1, 2, 3, 4 or 5 species. **b,** Distribution of the sequence conservation scores (calculated as z-scores of the distribution of phastCons30way^19^ values for non-coding regions in the same Topologically Associated Domain^58^; Methods) of human orthologous regulatory regions with different epigenetic states conserved in 1, 2, 3, 4 or 5 of our primate species.

Strong promoters are the most conserved activities: 80% of them are fully conserved in primates. On the contrary, poised and weak promoters are poorly conserved (Fig. 3a). All enhancer activities show a similar pattern of evolutionary conservation (Fig. 3a). Enhancer states with strong activities are second in conservation after strong promoters. Nearly 40% of the orthologous regulatory regions with strong enhancer states are fully conserved. Poised enhancers follow closely, with 36% of them conserved in the five species. Lastly, around 21% of the regions with a weak enhancer conserve their activity across primates. The regulatory regions associated with protein-coding and non-coding genes show the same conservation trends (Supplementary Figs. 23 and 24). However, strong activities in promoter states are less common for non-coding than for protein-coding genes, leading to lower conservation of promoter compared to enhancer states. This shows that differences in activity composition can lead to differences in the conservation of the regulatory architectures.

The epigenetic states in a given cell type and their evolutionary conservation reflect the specific function of the regulatory regions in this cell type. These regions are expected to show different states in other cell types and so their evolutionary patterns might also be different. To investigate whether changes in activity are likely to affect the epigenetic conservation of regulatory elements, we assessed the association between epigenetic and sequence conservation ‒which is cell type-independent. First, we observed that epigenetic conservation significantly correlates with the conservation of the underlying sequence ‒quantified as z-scores of background normalized PhastCons values^19^– in all epigenetic states but weak promoter states (Fig. 3b, Methods and Supplementary Fig. 25). These correlations are seen in the architectures of protein-coding but not in non-coding genes (Randomization analyses: 1,000 simulations; Fig. 3b, Supplementary Figs. 26-30). Of note, orthologous regulatory regions with fully conserved epigenetic states show significant differences in sequence conservation (Kruskal-Wallis test: *P* < 2.2 × 10^−16^; Supplementary Fig. 31). In particular, strong and weak promoters are associated with higher and lower sequence conservation scores respectively, whereas all enhancer states range in between these values. (Dwass-Steel-Critchlow-Fligner test, Supplementary Fig. 31). The sequence conservation scores associated with strong and poised enhancers are not significantly different. Note also that conserved poised promoters are associated with very high conservation z-scores, which probably did not reach significance due to their low number (n = 9 pP). Orthologous regions associated with non-coding genes are fewer and less epigenetically conserved (Supplementary Figs. 24 and 27), which could explain the lack of correlation between the conservation of the sequence and the epigenetic state observed in all but strong enhancers (Supplementary Fig. 30).

These results demonstrate that a detailed classification of promoters and enhancers with different activities into regulatory architectures provides a deeper understanding of their evolutionary constraints and dynamics, expanding previous observations in mammals^7^ that could mostly be made for active regulatory activities. The consistent association of epigenetic and sequence conservation also suggests that the epigenetic conservation observed in LCLs is a good proxy for the conservation of the regulatory activity of these elements in our primate species.

### Definition of different types of components in the regulatory architectures

To characterize the evolution of regulatory elements based on their specific role in gene expression, we classified regulatory elements into five different components according to the role they had in the gene regulatory architectures (Fig. 4a, Methods). We first classified regulatory elements based on their proximity to a gene into three types of components: genic promoters (gP), intragenic enhancers (gE), and proximal enhancers (prE). As gene expression is controlled by a combination of short- and long-distance regulatory interactions^20^, we used available 3D chromatin contact maps for human LCLs^21–23^ to link interacting regulatory elements to their target gene/s and define two additional types of components: promoter-interacting enhancers (PiE) and enhancer-interacting enhancers (EiE) (Fig. 4a).

**Figure 4.**
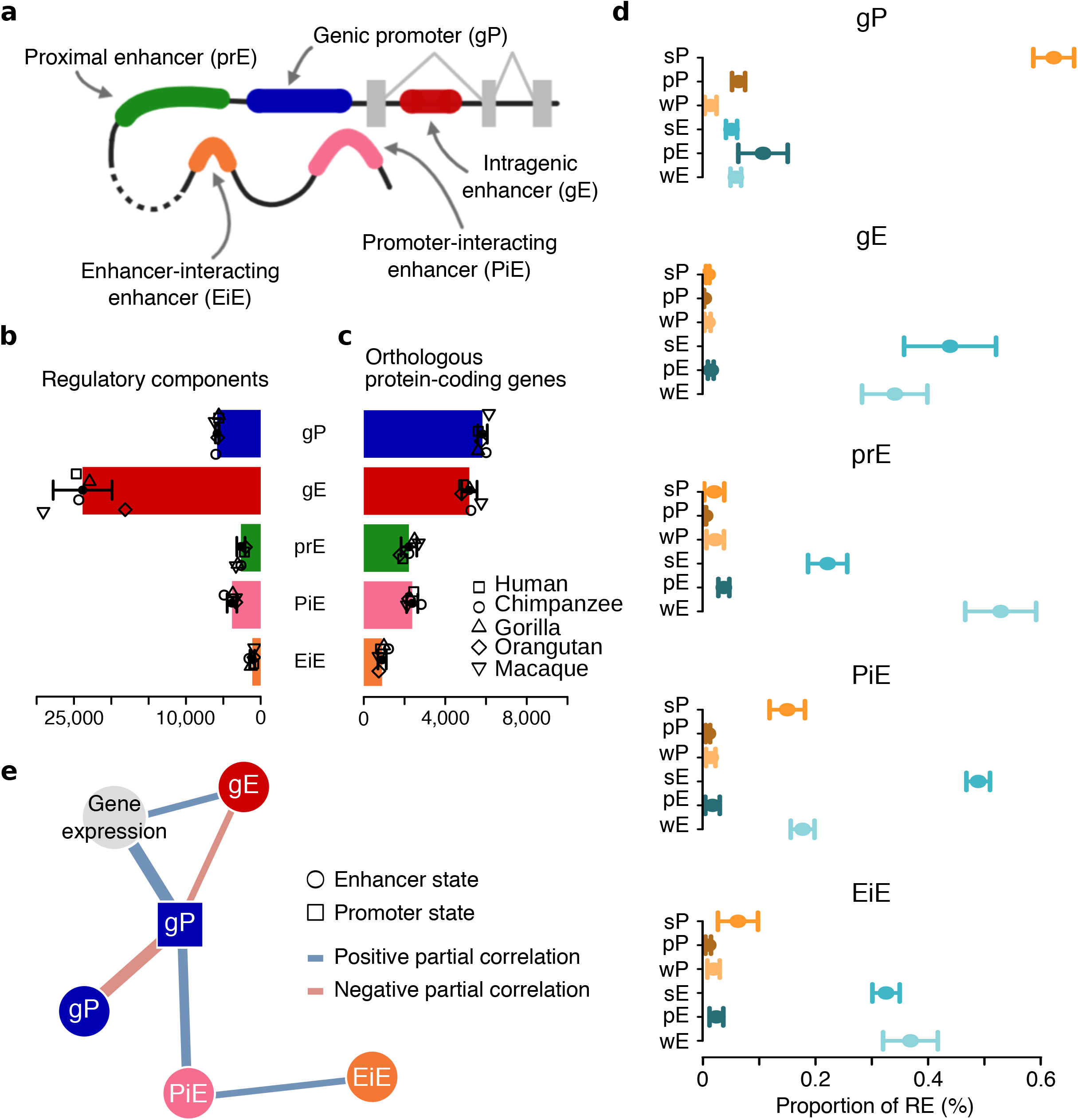
Epigenetic signals in gene regulatory architectures explain gene expression levels. **a,** Classification of regulatory elements according to their regulatory roles in gene architectures. Of note, EiE may interact with any type of enhancer in a regulatory architecture (prE, gE, PiE and EiE). **b,** Average number of orthologous protein-coding genes associated with each type of regulatory element. **c,** Average number of regulatory elements across species associated with 1-to-1 orthologous protein-coding genes classified as gP, gE, prE, PiE and EiE. Error bars show the standard deviation across species and differently shaped dots show the number corresponding to each species. **d,** Proportion of regulatory elements with a given epigenetic state associated with 1-to-1 orthologous protein-coding genes for each type of regulatory component. Dots and error bars show the average proportion and standard deviation across species, respectively. **e,** Sparse Partial Correlation Networks showing the statistical co-dependence of the RNA-seq (Gene expression) and the consensus ChIP-seq signals for the five histone marks in the every component represented by the eigencomponents (minimal partial correlation = −0.41; maximal partial correlation = 0.33; all partial correlations Benjamini-Hochberg’s *P* < 4.1 × 10^−303^). Edge widths are proportional to absolute partial correlation values within each network. The networks are based on the 5,737 1-to-1 orthologous protein-coding genes associated with at least one regulatory element in all species. Only nodes for values with significant and relevant partial correlations are represented.

We were able to link to genes and classify, on average, nearly 3,500 otherwise orphan distal regulatory elements per species (Supplementary Fig. 32). We annotated ~12,500 genic promoters, ~35,000 intragenic enhancers, ~6,700 proximal enhancers, ~6,200 promoter-interacting enhancers, and ~1,800 enhancer-interacting enhancers per species (Fig. 4b and Supplementary Fig. 33), of which 48%, 69%, 40%, 62%, and 61% are associated with 1-to-1 orthologous protein-coding genes in all primate species (Fig. 4c).

To assess the consistency of our classification of regulatory components, we focused on 1-to-1 orthologous protein-coding genes considering all their associated regulatory elements (i.e. 6 epigenetic states × 5 components = 30 regulatory subcategories). We found high concordance between the epigenetic state (based on ChIP-seq and ATAC-seq data, Fig. 2a) and the component (based on the type of association with the gene, Fig. 4a) of the regulatory elements. On average, 75% of genic promoters have a promoter state, and 90% of gene-associated enhancers have an enhancer state (Fisher’s exact test: *P* < 2.2 × 10^−16^ in all species, average *OR* = 64; Supplementary Fig. 34). This concordance is also consistent across species (Chi-square test: *P* < 2.2 × 10^−16^ in all species; Fig. 4d and Supplementary Fig. 35). Genic promoters are enriched in regulatory elements with strong promoter and poised promoter and enhancer states. Strong enhancers are mostly enriched at intragenic and promoter-interacting enhancers, whereas weak enhancers are strongly associated with proximal enhancers (Supplementary Figs. 34 and 35).

Gene expression levels are positively associated with the presence of strong activities in their regulatory architectures and are negatively associated with the presence of poised or weak activities (Kruskal-Wallis test: *P* < 0.05 in all species and regulatory components; Supplementary Fig. 36). These associations are particularly strong in genic promoters and intragenic enhancers (Dwass-Steel-Critchlow-Fligner test; Supplementary Fig. 37). Despite the consistency between the components’ activities and gene expression, our results suggest that different types of components might contribute differently to the regulation of gene expression.

### Distinct regulatory components influence gene expression and its evolution differently

To explore the ability of our classification of components to discriminate different regulatory roles, we disentangled the underlying network of regulatory co-dependencies between the different regulatory components and gene expression in our cell-type. For this, we used Sparse Partial Correlation Analysis (SPCA)^24^ of the normalized RNA-seq and histone mark enrichments (aggregated by promoter and enhancer state in every type of regulatory component) (Methods). This approach establishes a stringent protocol (Benjamini-Hochberg’s correction, *P* < 1.8 × 10^−22^ for all selected partial correlations) that selects informative partial correlations^24^.

To unravel the contribution of each type of component to gene expression, we defined their consensus signal (or eigencomponents) inspired by the notion of eigengenes^25^ (Methods). An SPCA based on the eigencomponents shows a consistent global structure of regulatory interactions, with genic promoters and intragenic enhancers directly regulating gene expression coordinately, promoter-interacting enhancers connected with promoters and enhancer-interacting enhancers connected with promoter-interacting enhancers (Fig. 4e and Supplementary Table 12). This regulatory scaffold is consistently observed for the residuals of the histone marks for these eigencomponents (Supplementary Fig. 38 and Supplementary Table 13) when SPCA was performed for all the histone marks together (Supplementary Fig. 39 and Supplementary Table 14) and for each of them separately (Supplementary Figs. 40-44 and Supplementary Table 15). To account for the possibility of incompleteness in some of our architectures, we replicated all the analyses using only genes with full regulatory architectures (i.e., genes associated with regulatory components of every type) obtaining consistent results (Supplementary Figs. 45-52 and Supplementary Table 15).

In agreement with the structure of regulatory interactions recovered by our SPCAs, a generalized linear model of gene expression based on H3K27ac, H3K27me3, and H3K36me3 signals at genic promoters and intragenic enhancers and their interactions (15 variables) explains ~67% of gene expression variability (Supplementary Table 16). Remarkably, this is only 6% lower than an exhaustive naive model, including the signal from all histone marks at all types of regulatory components with all possible interactions (1,225 variables) (Supplementary Table 17). These results confirm that the epigenetic activities of genic promoters, intragenic enhancers, and their interactions are likely the most direct determinants of gene expression regulation in our regulatory architectures. However, their co-dependency with the other components suggests that they are dependent, in turn, on the coordination of the whole architecture. These networks reflect that regulatory co-dependencies between components depend on the distance of the elements in the network of chromatin contacts (with genic promoters and intragenic enhancers being in the gene locus, promoter-interacting enhancers interacting directly, and enhancer-interacting enhancers interacting indirectly with it). The robustness of these networks of direct co-dependencies, their ability to explain gene expression, and their correspondence with the spatial disposition of the elements show that these components reflect specific regulatory roles.

Previous studies have found that gene expression evolution is associated with changes in the regulatory complexity of a gene (the number of close regulatory elements)^9^. Since we could classify the regulatory elements of a gene into different components (Supplementary Fig. 53), we were able to investigate the association of gene expression changes (Supplementary Figs. 54 and 55) with the evolutionary differences in the complexity of each type of component. We found that the effect of changes in complexity on gene expression levels depends on the epigenetic state gained or lost and the type of regulatory component affected (Supplementary Fig. 55). Evolutionary changes that alter the epigenetic state at genic promoters, specifically the presence of either a strong promoter or poised enhancer, as well as the number of intragenic enhancers with either strong or poised enhancer states, show the most robust associations with gene expression differences (Supplementary Fig. 55). The number of proximal enhancers in any enhancer epigenetic state and strong promoters and strong and poised enhancers in promoter-interacting enhancers also show significant though modest effects (Supplementary Fig. 55). These results highlight that the additive nature of gene regulation depends on regulatory architectures. This dependency can be captured either by the aggregation of histone enrichment signals (as in our SPCAs) or by quantifying the number of regulatory components with specific activities. Moreover, they confirm that our regulatory components represent different regulatory roles with a different contribution to gene expression evolution and which evolutionary relevance should be investigated separately.

### Poised and weak enhancers in genic promoters and intragenic enhancers appear in brain-specific genes with neuronal functions

We next explored to what extent the conservation and species-specificity of the characteristic regulatory states in every component (overrepresented combinations, Supplementary Fig. 56) is important for particular functional processes. For this, we examined their functional annotation and tissue-specificity in their expression (GTEx data^26^, Supplementary Table 18). We found significant enrichment for the genes targeted by conserved strong promoter states in genic promoters, conserved strong and weak enhancer states in intragenic enhancers, and conserved poised enhancer states in genic promoters and proximal enhancers (Fisher’s exact test: Benjamini-Hochberg’s correction, *FDR* < 0.05; Methods, Fig. 5a, Supplementary Fig. 57 and Supplementary Table 19). Remarkably, among the genes associated with species-specific epigenetic states, only those linked to human-specific weak enhancers in intragenic enhancers had significant functional enrichments.

**Figure 5.**
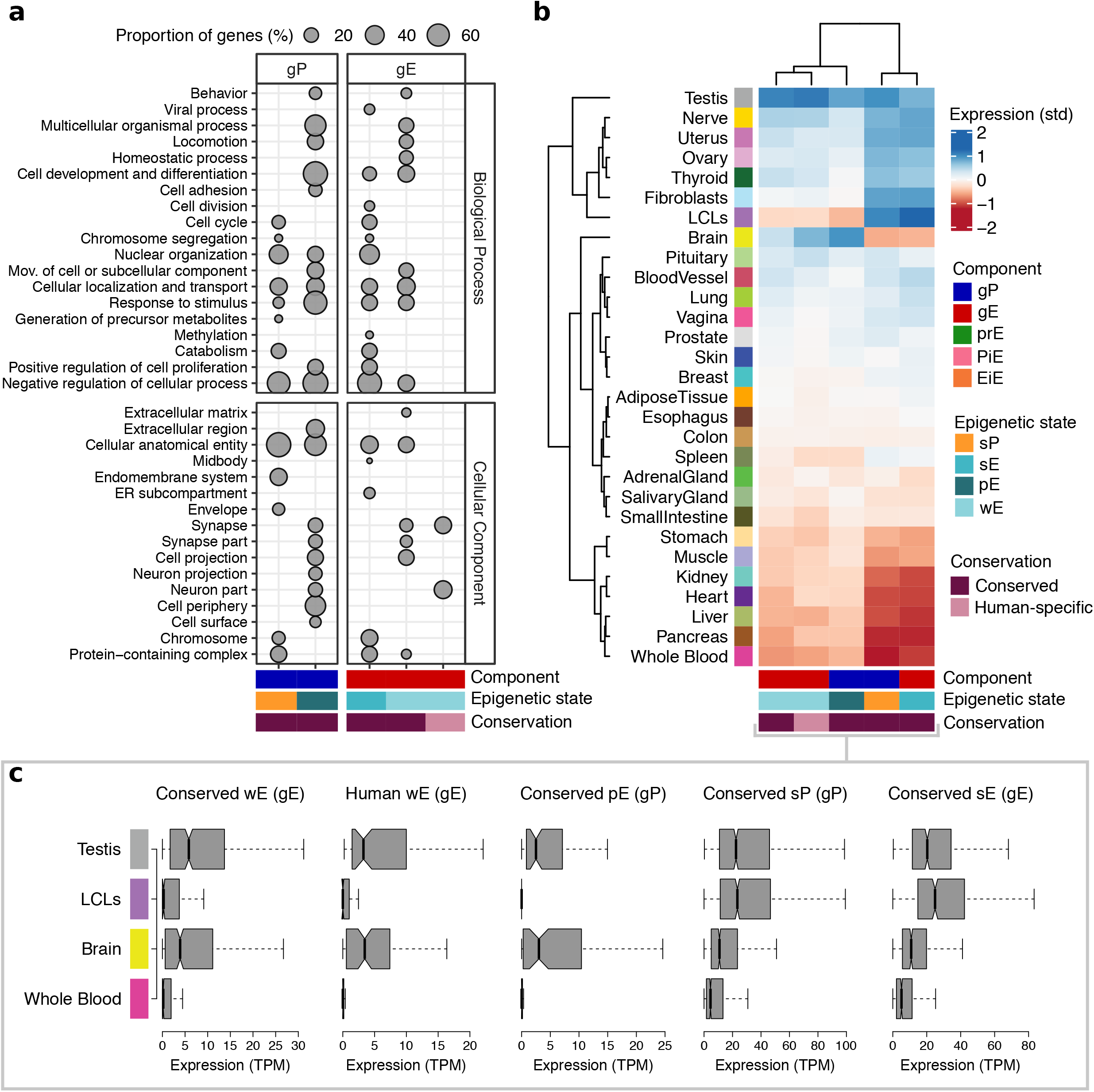
Weak and poised enhancer states echo brain-specific regulation. **a,** Functional enrichment of conserved and human-specific activities in genic promoters and intragenic enhancers. The size of circles indicates the proportion of genes included in each functional category from the total number of genes contained in the corresponding regulatory group. Number of genes in each category and extended functional annotation in Supplementary Fig. 57. **b,** Heatmap of standardised expression across tissues in state/component regulatory groups with functional enrichments. Number of genes included in each category and representation with groups without functional enrichment in Supplementary Fig. 58. **c,** Median expression levels in testes, LCLS, brain and whole blood of genes groups in b.

These enrichments show the expected association of conserved strong epigenetic states (strong promoter states in genic promoters and strong enhancer ones in intragenic enhancers) with genes involved in relevant cellular processes, such as metabolism, chromatin organization, and regulation of the cell cycle (Fig. 5a, Methods, Supplementary Fig. 57 and Supplementary Tables 20 and 21). Functions specific to LCLs like those involving viral processes are specifically enriched in strong enhancers. Moreover and regardless of whether there is an enrichment, all three strong epigenetic states show similar expression profiles across tissues (Fig. 5b and Supplementary Figs. 58-60), with high expression levels in LCLs and most other tissues and wide expression breadth (Fig. 5c and Supplementary Fig. 61).

In contrast, conserved poised enhancer states in genic promoters and proximal enhancers target protein-coding genes enriched in developmental and proliferative functions, echoing their known implication in these processes (Fig. 5a, Supplementary Fig. 57 and Supplementary Tables 22 and 23). Surprisingly, genes with conserved poised enhancer states in their genic promoters are also enriched in neuronal functions and higher expression levels in brain (Fig. 5b-c and Supplementary Figs. 57-60). Genes associated with both types of conserved poised enhancer states show overall minimal expression levels but high tissue-specificity (median tissue specificity index (τ, Tau) > 0.85 in both; Methods and Supplementary Fig. 61).

Protein-coding genes targeted by intragenic enhancers with conserved weak enhancer states are enriched in various functional annotations, including neuronal ones, such as cell projection and synapse (Fig. 5a, Supplementary Fig. 57 and Supplementary Table 24). This gene setshows remarkably low expression in LCLs coupled with higher expression in the brain, which is in agreement with the observed functional annotation (Fig. 5b-c). Also, the tissue-specificity of this group is higher than that of conserved strong regulatory activities both in promoters and enhancers (median τ = 0.72, Dwass-Steel-Critchlow-Fligner test: *P* < 2.2 × 10^−16^ in the three tests; Supplementary Fig. 61). This apparent brain-specificity is not found in genes associated with other weak enhancer states that have overall higher expression levels and not particular specificity, as would be expected from weak epigenetic states (Supplementary Figs. 60 and 61).

Finally, we focused on genes targeted by components with human-specific epigenetic states. These genes are solely enriched in neuron parts and synapse (Fig. 5a and Supplementary Table 25). Similar to genes associated with their analogous conserved group, these genes are typically expressed at low levels with highest expression in tissues unrelated to LCLs, particularly brain, tibial nerve, and testis, while having marginal or no expression in numerous other tissues, including LCLs (Wilcoxon-Nemenyi-McDonald-Thompson test: *P* < 1 × 10^−4^; Rank-biserial correlation effect size between brain and LCLs = 0.633; Fig. 5b-c, Supplementary Figs. 58 and 59 and Supplementary Table 26). Remarkably, these genes have higher tissue-specific expression than those with conserved strong but not weak activities in their components (median τ = 0.84, Dwass-Steel-Critchlow-Fligner test: *P* < 4.5 × 10^−14^ when compared to strong activities and *P* = 0.06 compared to genes with weak enhancer states; Supplementary Fig. 61).

Intrigued by the high tissue-specificity of the genes with novel human weak enhancers, we sought to identify the tissues driving this tissue-specificity taking its analogous conserved group as reference. Testis and brain are the tissues with the highest number of tissue-specific genes (τ_Tissue_ > 0.8), but most interestingly, whereas the fraction of testis-specific genes is comparable between gene sets (Two-tailed Fisher’s exact test: *P* = 0.54, *OR* = 1.20), brain-specific genes are more than 2-fold enriched in genes with human-specific intragenic enhancers (Two-tailed Fisher’s exact test: *P* = 0.02, *OR* = 2.29; Supplementary Fig. 62).

Altogether these results show that while conserved strong epigenetic states are involved in the regulation of important genes highly expressed in LCLs and other tissues, conserved poised enhancer states and conserved and human-specific weak enhancer states in intragenic enhancers are involved in the regulation of genes marginally expressed in LCLs, but with particular functional roles and tissue-specific expression patterns. These unexpected associations are likely to reflect the importance of particular epigenetic states in certain regulatory components to regulate specific processes.

### Genes with novel human-specific intragenic weak enhancers are targeted by positive selection

The unanticipated association of the genes targeted by human-specific weak enhancer states in intragenic enhancers with neuronal functions prompted us to study the relationship these genes might have with positive selection. In fact, among the genes associated with intragenic enhancers with novel human-specific weak activities, we found several genes previously proposed as candidates for positive selection in humans^27–30^. Some of these genes are *FOXP2*, *PALMD*, and *ROBO1*, which have known brain-related functions^31–34^ or *ADAM18*^*35*^, *CFTR*^*36,37*^, and *TBX15*^*38*^.

To assess whether genes with human-specific enhancer states have been targeted by recent human adaptation, we investigated their co-occurrence in genes associated with signals of positive selection^27–30^ (Methods and Supplementary Table 27). We found that more than one third (38%) of the genes with novel weak intragenic enhancers are associated with genes targeted by positive selection (Fisher’s exact test: *P* = 6.52 × 10^−18^, *OR* = 5.69). The results of this analysis (Fig. 6a) indicates that this enrichment is reflected in a significant association of genes targeted by intragenic enhancers (but not in any other component type), genes targeted by intragenic enhancers with weak enhancer states (but not strong or poised enhancer states) and genes targeted by intragenic enhancers with human-specific weak enhancer states (but not in fully conserved or the remaining weak enhancer states). No enrichment in signals of positive selection is observed for genes with genic promoters showing conserved poised enhancer states, even though they are also involved in brain-specific and neuronal functions.

**Figure 6.**
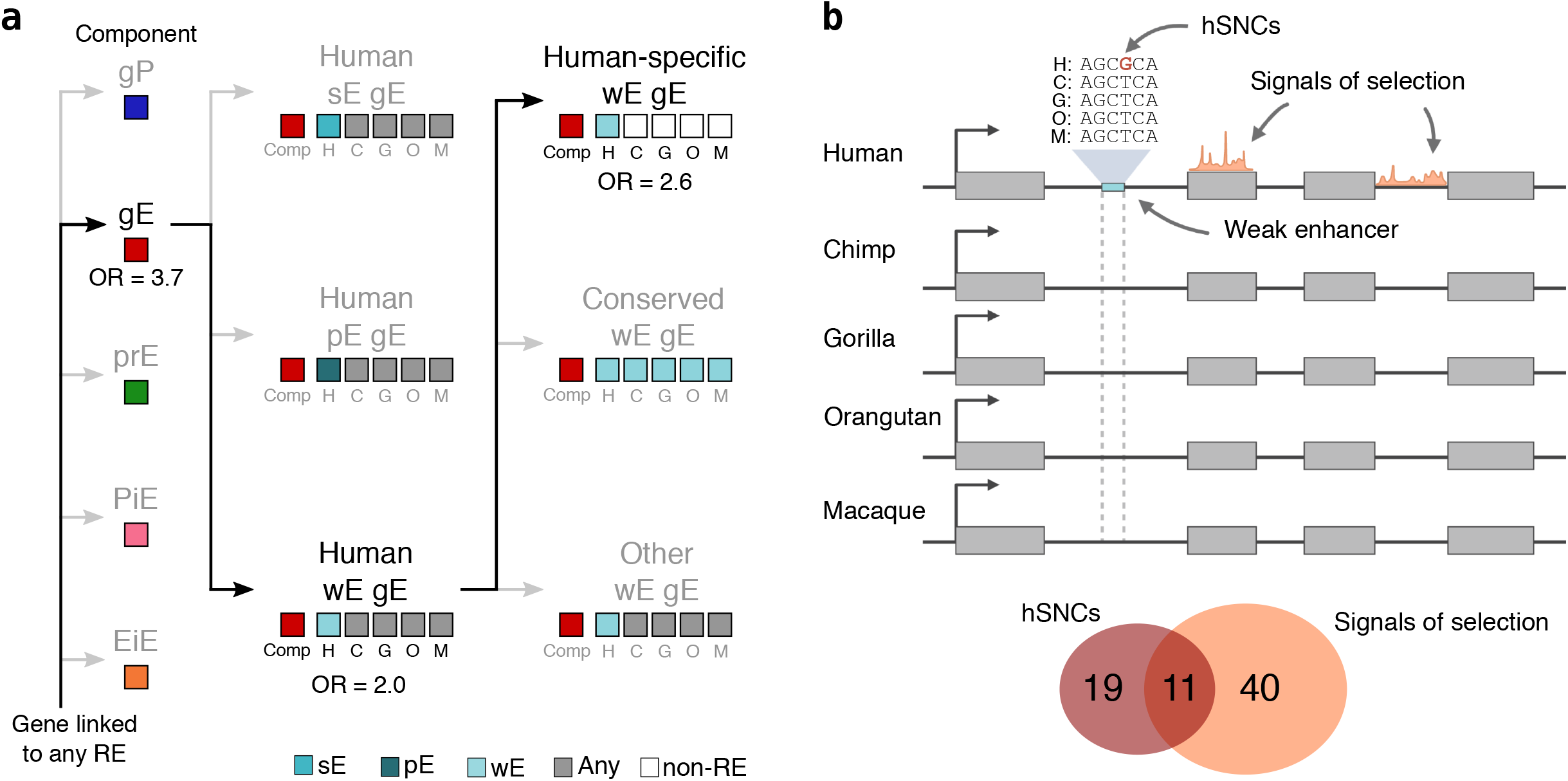
Intragenic enhancers with weak activities co-localize with signals of recent human selection. **a,** Specific enrichment of genes with signals of positive selection in genes that harbor human-specific intragenic enhancers with a weak enhancer state. Epigenetic states for each species are depicted in white, grey or blue boxes. **b,** Top: Schematic representation of a human-specific intragenic weak enhancer with a hSNC (nucleotide change in humans shown in red) contained in a gene with signals of selection. Bottom: Venn diagram illustrating the overlap between the 41 genes containing human-specific weak intragenic enhancers with signals of selection and the 30 genes with these enhancers and with human single nucleotide changes (hSNCs) fixed in humans and distinct from other non-human primates.

Finally, we explored whether these recently evolved human-specific intragenic enhancers are associated with human-specific mutations. For this, we collected a set of over 2.8 million single nucleotide changes fixed in humans (hSNCs) that differ from fixed variants in the genomes of the remaining non-human primates (Methods and Supplementary Table 27). We observed that the hSNCs density is higher in human-specific intragenic enhancers (Mann-Whitney U test: *P* = 0.01; Methods and Supplementary Fig. 62). More than one-third of the genes with novel human-specific intragenic enhancers with weak enhancers states and with hSNCs also have signals of positive selection, a proportion very similar to the expected 38% (see above). This result suggests that although human-specific mutations and positive selection signals are both associated with the presence of intragenic enhancers with human-specific weak activities, they are not mutually conditioned. As such, it implies that none of these signals is necessary (nor sufficient) to explain the appearance of intragenic enhancers with human-specific weak activities.

Among the 11 genes with both signals of positive selection and hSNCs (Fig. 6b), there are several interesting candidates for adaptive evolution of different traits. Many of these genes are associated with neuronal functions (*ROBO1, CLVS1, SEMA5A, KCNH7, SDK1*, and *ADGRL2*), but also with pigmentation (*LRMDA*) or actin organization in cardiomyocytes (*FHOD3*). Other interesting genes that include human-specific weak intragenic enhancers are only associated with signals of human selection (*FOXP2*, *TNIK*, *ASTN2*, *NPAS3*, or *NTM*) or hSNCs in these enhancers (*PALMD*, *VPS13C*, *IGSF21*, or *CADM2*). Interestingly, we found only one antisense RNA gene, *MEF2C-AS1* showing both signals of positive selection and a human-specific enhancer with hSNCs (Supplementary Fig. 63). This gene has been associated with ADHD^39^, and its target gene *MEF2C*, is a very well known target of genetic alterations (many of them also affecting *MEF2C-AS1*) associated with severe intellectual disability^40^, cerebral malformation^40^, or depression^40,41^.

Remarkably, three human-specific intragenic enhancers accumulate more hSNCs than expected (Randomization test: 10,000 simulations, Bonferroni correction, *P* < 0.02 in all cases; Methods and Supplementary Figs. 63 and 64), a number of enhancers which is also significantly higher than expected (Randomization test: 10,000 simulations, *P* = 8 × 10^−4^; Supplementary Fig. 65). Two of these genes are protein-coding genes with known functions in brain cell types and with signals of positive selection. *CLVS1* is a protein-coding gene with brain-specific expression (τ_Brain_ = 0.964) required for the normal morphology of endosomes and lysosomes in neurons^42^. *ROBO1* is a broadly expressed integral membrane protein that participates in axon guidance and neuronal migration (τ = 0.388)^43,44^ that has also been associated with human speech and language acquisition since the split from chimpanzees^32^. The third enhancer is included in *AC005906.2*, a long intergenic non-protein-coding gene specifically expressed in brain (τ_Brain_ = 1). Interestingly, this gene overlaps with *KCNA1*, a voltage-gated potassium channel with the same brain-specific expression pattern (τ_Brain_ = 0.995) and for which mutations have been associated with neurological malfunctions^45^.

Our results show that the most common regulatory innovation detected in human LCLs, the presence of human-specific weak enhancer activities in intragenic enhancers, targets neuron-related brain-specific genes that are significantly associated with signals of positive selection and an excess of hSNCs in these components. These DNA changes were especially concentrated in three of them. Two of the genes in which these elements are harbored, *CLVS1* and *ROBO1*, exemplify the confluence of signals of positive selection and excess of hSNCs in human-specific regulatory regions targeting protein-coding genes important for normal neuronal structure, migration, and axon guidance in the human brains.

## Discussion

The evolution of human and non-human primates is an area of major interest, but ethical, legal, and practical constraints often limit access to direct biological material. In this study, we have generated a unique, comprehensive, and unified dataset of epigenomic landscapes in LCLs for human and four non-human primate species. Despite the artificial nature of our cellular model^46–48^, previous studies have shown the value of LCLs as an experimentally convenient model of somatic cells that accurately resembles the phenotype of its cell type of origin^49^ and which can be robustly used for comparative studies in humans and primates^12,50–52^. Moreover, its clonality ensures a cell type-specific experimental system reducing the confounding factors associated with cell population diversity in bulk tissue samples.

Using this cell model, we reproduced previous observations on the dynamics of the evolution of regulatory elements reported in more distant species using liver samples^7,9,18^ which we show can be extrapolated to closely related species (at least for great apes and macaques). Moreover, we have expanded these observations to explain how these dynamics result from the different evolutionary constraints associated with their epigenetic activities. Therefore, we show that considering weak and poised activities is of major relevance to fully understand the evolution of regulatory regions.

We also observed that different epigenetic activities have characteristic evolutionary patterns with higher conservation for strong promoter and strong and poised enhancer states. The correlations between epigenetic and sequence conservations are also different for each epigenetic state with higher correlations for strong and poised promoter and enhancer states. These differences are likely due to their different influence on gene expression. Therefore, previously reported higher conservation of promoters probably reflects the often bigger influence of these elements in gene expression. This hypothesis is also confirmed by the lower conservation of promoter states observed in the regulatory architectures of the non-coding genes where strong promoter states are scarce.

Here, we have introduced a classification of regulatory elements as components of gene regulatory architectures into genic promoters and intragenic, proximal, promoter-interacting, and enhancer-interacting enhancers. The network of regulatory co-dependencies of these types of components reveals that the epigenetic activities of each type of regulatory component influence gene expression levels differently. In brief, coordinated epigenetic activities in genic promoters and intragenic enhancers form the core of these architectures and explain gene expression levels. Regulatory activities in promoter-interacting enhancers are also coordinated with promoter components, and activities in enhancer-interacting enhancers are associated with promoter-interacting enhancers. These results show that the influence of regulatory components on gene expression reflects the structure of the regulatory architecture.

The importance of this structure in gene expression evolution is reflected in the different association of gene expression changes with the regulatory complexity of each activity in the distinct components. The addition or removal of strong promoter activities in promoter components or strong and poised enhancer activities in intragenic enhancers consistently co-occurs with gene expression changes between primate species. The remaining components show fewer changes linked to expression differences, but they can still be instrumental for gene expression evolution, probably through their influence on promoters and intragenic enhancers. Our conceptual framework provides a starting point for future in-depth investigations on the inter-dependence of different regulatory regions and mechanisms in the evolution of gene regulation. In this sense, we stress the importance of considering promoter and enhancer activity states in the different types of gene components to achieve a more detailed description of the regulatory processes.

Despite the larger influence of strong activities on gene regulation, our results in LCLs suggest that major insights can arise from the analysis of the elements with a repressive or negligible regulatory role in our cell model. Genic promoters and proximal enhancers with poised enhancer activities and intragenic enhancers with weak enhancer activities carry information about the degree of regulatory innovation in unrelated cell types. Conserved poised activities are targeted by genes associated with cell proliferation and differentiation. In the case of poised enhancer states in genic promoters, these genes are specifically expressed in brain. Moreover, we found that recently evolved weakly active intragenic enhancers in the human lineage are the most common regulatory innovation observed in our LCLs. These human-specific weak enhancers occur in genes showing patterns of brain-specific gene expression, neuronal functions, and signals of positive selection, suggesting that these genes may have contributed to human adaptation in several traits. These functional and evolutionary patterns are different from those in genes targeted by any conserved activity in any other component, including conserved poised enhancers in genic promoters that also target genes with brain-related functions.

We have identified a subset of genes in which regulatory innovation in these intragenic enhancers converges with other signals of positive selection. Among these genes, we highlight two protein-coding genes key in neuronal structure, migration, or axon guidance: *CLVS1* and *ROBO1*, also accumulating an excess of human-specific mutations in the corresponding human-specific enhancers. The confluence of epigenetic and sequence innovations in the human lineage for these genes points to their putative relevance in recent human evolution. Our findings suggest that the appearance of novel intragenic enhancers with tissue-specific and functionally relevant implications in certain genes is often bound to the co-appearance of weaker activity signals that can be detected in other cell types. These echoes that we detect as human-specific weak enhancer activities provide an unexpected window to the study of regulatory evolution in the human lineage. Further research will be needed to clarify the specific role of these elements in different tissues and cell types.

Taken together, our results show that the evolution of gene regulation is deeply influenced by the coordination of epigenetic activities in gene regulatory architectures. Our insights call for incorporating better integrative datasets and refined definitions of regulatory architectures in comparative evolutionary studies to fully understand the interplay between epigenetic regulation and gene expression.

## Methods

### Definition of regulatory elements

We used ChromHMM to jointly learn chromatin states across samples and segment the genome of each sample^17^. ChromHMM implements a multivariate Hidden Markov Model aiming to summarize the combinatorial interactions between multiple chromatin datasets. Bam files from the five histone modifications profiled were binarized into 200 bp density maps. Each bin was discretized in two levels, 0 or 1, depending on their enrichment computed by comparing immunoprecipitated (IP) versus background noise (input) signal within each bin and using a Poisson distribution. Binarization was performed using the BinarizeBam function of the ChromHMM software^17^. A common model across species was learned with the LearnModel ChromHMM function for the concatenated genomes of all samples but O1 (orangutan sample 1) due to its anomalous epigenetic profiles (Supplementary Fig. 76). Several models were trained with a number of chromatin states ranging from 8 to 20. To evaluate the different n-state models, for every sample, the overlap and neighborhood enrichments of each state in a series of functional annotations were explored. A 16-state model was selected for further analysis based on the resolution provided by the defined chromatin states, which capture the most significant interactions between histone marks and the state enrichments in function-annotated datasets (Supplementary Fig. 2). The genomic coordinates of regulatory elements (RE) were defined for each sample by merging all consecutive 200 bp bins excluding elongating (E1 and E2), repressed heterochromatin (E16) and low signal (E15) chromatin states. Species regulatory elements were defined as the union of sample regulatory elements. For orangutan we did not include regulatory elements specific to O1.

### Assignment of a regulatory state to regulatory elements

Regulatory elements were assigned a chromatin-state based annotation. Combining the information gathered through the overlap and neighborhood enrichment analyses in functionally defined regions, we established a hierarchy to designate poised (p), strong (s) and weak (w) promoter and enhancer states. Chromatin states E8, E9 and E11 defined promoter states (P); E8 and E9 were strongly enriched at TSSs, CGI, UMR (unmethylated regions) and open chromatin regions, while E11 was mostly located downstream the TSS; the presence of E14 defined poised promoter states (pP); absence of E14 and presence of E9 or E11 defined strong promoter states (sP); remaining P were classified as weak promoter states (wP). Non-promoter regulatory elements were assigned an enhancer state (E). The presence of E14 defined poised enhancer states (pE); absence of E14 and presence of E3, E4, E5, E6 and E12 defined strong enhancer states (sE): E5 and E6 were strongly enriched LMRs (low methylated regions) whereas E3, E4 and E12 were highly abundant at introns; remaining E were classified as weak enhancer states (wE) (Supplementary Figs. 2 and 66).

One of the limitations of chromatin states is that bin assignments are based on the presence or absence of particular epigenetic marks. However, oftentimes, the lines separating different regulatory elements are blurry: e.g. the distinction between promoter and enhancer states generally resides in the H3K4me3/H3K4me1 balance. Hence, some misclassifications are expected due to insufficient precision of the qualitative classification. Considering the quantitative relationship between co-existing histone modifications can help to accurately annotate epigenetic states in regulatory elements. We used linear discriminant analysis (LDA)^53^ to refine chromatin-state based annotations. This method is commonly applied to pattern recognition and category prediction. LDA is a technique developed to transform the features into a lower-dimensional space, which maximizes the ratio of between-class variance to the within-class variance, thereby granting maximum class separation. We performed LDA analysis using the lda function in the R package MASS (version 7.3-47)^54^. The predictor variables were the background-noise normalized IP signals from the five different histone modifications profiled and chromatin accessibility signal at species regulatory elements. The categorical variable to be predicted based on the underlying enrichments was the chromatin-state based annotation. The regulatory state at the species level was determined based on the regulatory state in each of the biological replicates. Thus, the regulatory state of a regulatory element with different epigenetic states in the two replicates (ambiguous), could be aP or aE, when both samples of a given species were annotated as either P or E but differ in their activity; P/E, when a regulatory element was classified as P in one biological replicate and E in the other one; and P/Non-RE or E/Non-RE, when the regulatory elements was so only in one replicate (Supplementary Fig. 7 and Supplementary Table 1). To control for interindividual variability, only regulatory elements with the same activity in the two replicates were considered for downstream analyses.

### Analysis of evolutionary conservation at orthologous regulatory regions

We studied patterns of evolutionary conservation of promoter and enhancer states using a set of 21,753 one-to-one orthologous regions associated with genes in which at least one species showed a promoter or enhancer epigenetic state. We define *recently repurposed promoters* as orthologous regulatory regions in which one species shows a promoter state while the others show an enhancer state or vice versa. *Novel promoter or enhancer states* refer to those orthologous regulatory regions in which a given species showed a promoter or enhancer state while the others showed no evidence of regulatory activity (classified as *non-regulatory*).

To study the patterns of evolutionary conservation of regulatory states, we focused on the subset of 10,641 one-to-one orthologous regions in which at least one species showed a strong, poised or weak regulatory state (we do not include orthologous regions including elements with ambiguities, ie. different activities between biological replicates). To statistically assess the different evolutionary dynamics observed for the different regulatory states we first ran randomization analysis. We randomized (1,000 randomizations) the regulatory states associated with each species in orthologous regulatory regions. We determined the P-value as the number of randomizations with an average conservation equal to or above the observed conservation for each regulatory state. We further explored the different patterns of conservation combining: (1) Kruskal-Wallis test (kruskal.test R function)^55^ to test whether the global distributions of the number of species in which each particular state was conserved were different for the different regulatory states and (2) Dwass-Steel-Critchlow-Fligner test to assess the significance of every pairwise comparison (dscfAllPairsTest function from the R package PMCMRplus version 1.4.4)^56^ and (3) Glass rank biserial correlation coefficient for Mann-Whitney U test to compute the effect sizes associated with all statistically significant pairwise comparisons (wilcoxonRG function from the R package rcompanion version 2.3.25)^57^.

To study the patterns of evolutionary conservation of the sequence underlying orthologous regulatory regions, we assigned each orthologous regulatory region a conservation score. We computed this score based on the phastCons30way sequence conservation track^19^. To control for background sequence conservation levels, we first computed the average and standard deviations phastCons30way in TADs defined in the cell line GM12878^58^ (Supplementary Fig. 25). Then, we used these summary statistics to calculate the z-score for each bp in every orthologous regulatory region, using the average and standard deviations values of the TAD in which each orthologous regulatory region was found. We averaged the z-scores within each orthologous regulatory regions in bins of 200 bp that overlap 50 bp with the next bin and assign each orthologous regulatory region the maximum z-score values associated with its bins. We computed the Spearman rho correlation between the z-scores and the number of species in which each orthologous regulatory region was conserved, separately for each regulatory state. To determine the statistical significance of these correlations we used randomization analysis. For each regulatory state we created 1,000 sets randomizing the z-score associated with each orthologous regulatory region and calculated the Spearman correlation in each randomization. We determined the P-value as the number of randomizations with a Spearman rho correlation value equal to or above the observed correlation (Supplementary Figs. 28-30).

### Classification of regulatory elements in different types of components of gene regulatory architectures

We pre-classified each regulatory element into gene regulatory component based on their genomic location with respect to their corresponding species ENSEMBL release 91^59^ gene annotations. Regulatory elements found up to 5 Kb upstream to the nearest TSS were classified as genic promoters (gP). Additional regulatory elements located up to 10 Kb to the nearest TSS were classified as proximal enhancers (prE). Regulatory elements that overlapped a gene were classified as intragenic enhancers (gE). Other regulatory elements that could not be linked to a gene based on their genomic proximity were initially classified as distal enhancers (dE).

Then, we made use of available interaction data for the cell line GM12878 (HiC^21^, HiChIP-H3K27ac^22^ and ChIA-PET^23^) to map interactions between regulatory elements. Each interacting pair was mapped independently to hg38 coordinates using the liftOver tool from the UCSCTOOLS/331 suite^60^, and only interactions for which both pairs could be mapped were kept. Subsequently, interactions were mapped to the non-human primate reference genome assemblies. For inter-species mappings, coordinates were mapped twice, going forward and backward, and only pairs that could be mapped in both directions were kept. Interacting regulatory elements were defined as those that overlapped with each pair of any given interaction. First-order interactions were annotated between promoters and enhancers, allowing the definition of promoter-interacting enhancers (PiE). Second-order interactions were annotated between enhancer components (gE, prE or PiE), allowing the definition of enhancer-interacting enhancers (EiE) (Fig. 4a and Supplementary Fig. 1).

Considering both classifications of regulatory elements, according to their epigenetic state and regulatory component, regulatory elements were separated into 30 (6×5) different subcategories. We used a Chi-square test to identify the component-epigenetic state combinations enriched in orthologous regulatory regions with fully conserved and species-specific epigenetic states (Supplementary Fig. 56).

### Gene expression levels and regulatory states in gene components

To investigate the influence of the activity state of regulatory elements in each type of component on gene expression levels, we classified 1-to-1 orthologous protein-coding genes, separately for each species, into six mutually excluding categories, one for each regulatory state within each type of component (component-state combinations). Whereas genes can only be associated with one genic promoter and hence, they can only be classified into one category for genic promoters depending on the corresponding epigenetic state of the regulatory element, genes can be associated with more than enhancer component (gE, prE, PiE and EiE). In those cases we classified genes into a given component-state category accordingly to the presence of at least one regulatory element with a given epigenetic state in that component using the following state hierarchy: pE > pP > sE > sP > wE > wP (Supplementary Fig. 36). To statistically assess the influence of each state in each component we used (1) Kruskal-Wallis test (kruskal.test function as implemented in R)^55^ to test whether the distributions of the expression levels of genes associated with each component-state combination were different for the different regulatory states, (2) Dwass-Steel-Critchlow-Fligner test to assess the significance of every pairwise comparison (dscfAllPairsTest function from the R package PCMRplus version 1.4.4)^56^ and (3) Glass rank biserial correlation coefficient effect size for Mann-Whitney U test to compute the effect sizes associated with all statistically significant pairwise comparisons (wilcoxonRG function from the R package rcompanion version 2.3.25)^57^ (Supplementary Fig. 37).

### Partial correlation analysis

To disentangle the network of direct co-dependencies between the different components, regulatory states, histone marks and gene expression, we performed a series of partial correlation analyses^24,61^. To tackle the diversity of architectures detected for the different genes, we added up the calibrated signal of all the regulatory elements with a given regulatory state (promoter or enhancer) in a given type of component for any gene architecture. This decision was based on the observed relationship between the number of strong elements in a gene architecture and the expression level of its target gene. Separation of histone signals in each type of component between those contributing to a promoter or to an enhancer was intended to reflect the potential differences in their role in gene expression regulation. As a result of this design, our system has 51 variables (RNA-seq signal + 5 histone mark signals × 2 regulatory states × 5 components) and 57,370 cases (5,737 genes × 5 species × 2 samples).

All partial correlation analyses were performed using an adaptation of a recently published Sparse Partial Correlation Analysis protocol^24^ based on the continuous values of the accumulated ChIP-seq signals (instead of their ranks) to take advantage of their pseudo-quantitative nature. This protocol combines the recovery of statistically significant partial correlations with a cross-validation process to filter out those relationships leading to overfitted reciprocal linear LASSO models (significant partial correlations unlikely to be biologically meaningful). In our case, in every analysis, we recovered those partial correlations recovered in at least four of the five species without leading to overfitting when determining the reciprocal explanatory power in the remaining species. This protocol is intended to detect biologically relevant co-dependences out of the set of significant partial correlations and as a result, this approach filters out many significant partial correlations with very low explanatory power. In fact, all the partial correlations recovered in any of the analyses performed showed very low P-values (Benjamini-Hochberg’s correction^62^, *P* < 1.8 × 10^−22^). In our case, given the relatively small amount of data, we focused on recovering those partial correlations that are likely to be relevant in any species. For these analyses, we used a modified version of the R code provided by the authors (http://spcn.molgen.mpg.de/code/sparse_pcor.R/) to perform 5-fold cross-validation analyses separati by species instead of the original 10-fold cross-validation protocol suitable for larger datasets. Network visualizations were performed with Cytospace^63^.

In a partial correlation model, direct co-dependencies are established between individual variables. However, we know that coordination of the different histone marks within components is important to define the global epigenetic configuration of a component (also captured in our epigenetic states), which itself could be considered the relevant variable for this analysis. To better address this situation in our analysis, we defined a consensus signal for every component following the same approach established by WGCN^25^ to define eigengenes as representative variables of clusters of co-expressed genes. In brief, we defined eigencomponents as the variables summarizing the common signals of the different histone marks in a component (actually calculated as the first PCA component of these five variables). So that eigencomponents keep the meaning of the activities, they were defined as codirectional with H3K27ac signals in each component (eigenvectors negatively correlated with H3K27ac signals were multiplied by −1). We performed a Sparse Partial Correlation Analysis of these 10 eigencomponents and RNA-seq that recovers very clearly the structure of direct co-dependecies between the epigenetic configuration of the different components and gene expression (Fig. 4e and Supplementary Table 12).

In addition, we defined the remaining unexplained signal of every histone mark by its eigencomponent as the residuals of a linear model of the original variables and the corresponding eigencomponent. A Sparse Partial Correlation Analysis of these residuals (Supplementary Fig. 38 and Supplementary Table 13) shows that even these residuals reflect the same inter-component structure and highlights that our eigencomponents miss some relevant information for the definition of this regulatory coordination (mainly weaker co-dependencies involving promoter states in intragenic and promoter-interacting enhancers and enhancer states in promoters).

To assess to what extent eigencomponents reflect the behavior of the whole network of co-dependencies of the histone marks or of each of the specific histone marks, we also performed SPCAs using the actual ChIP-seq enrichment signals. A global partial correlation analysis considering all 51 variables shows a very clear structure of direct co-dependencies with a strong intra-component contribution for the two states of every single component and a clear but more modest exclusive inter-component contribution (Supplementary Fig. 39 and Supplementary Table 14). Analyses to determine the Sparse Partial Correlation Network of each of the histone marks and RNA-seq without considering the possible influence of the remaining histone marks (Supplementary Figs. 40-44 and Supplementary Table 15) retrieve very similar networks pointing to the common backbone of inter-component co-dependences reflected in our SPCA of the eigencomponents.

Our dataset of regulatory components shows a quite unbalanced contribution of the components to the architectures, with intragenic enhancers being the most abundant type of component and promoter-interacting and enhancer-interacting enhancers being the least abundant (Supplementary Fig. 33). These differences could be at least partially related to our inability to recover some of the chromatin interaction-mediated regulatory associations. More importantly, this imbalance, if not real, could affect the ability of our partial correlation networks to reflect the contribution of those components less represented in our datasets. To explore this point, we recovered the subset of genes (an average of 1068 genes per sample) with full architectures (those with at least one element in every type of component) and repeated all the Sparse Partial Correlation Analyses explained above with this dataset of genes. In all the cases, we obtained very similar results, recovering fewer relevant partial correlations due to the smaller number of genes, but with no signal of any relevant difference in the global structure of the coordinated network of components and gene expression (Supplementary Figs. 45-52 and Supplementary Tables 12-15).

All the components of the connected network can be very influential in gene expression through their direct or indirect connection with gene expression. However, our Sparse Partial Correlation Networks point consistently to the direct co-dependency of RNA-seq with the genic promoter and intragenic enhancer components and the co-dependency between them. To quantify the explanatory power of these dependencies for gene expression, we performed a simple generalized linear model (glm function as implemented in R^55^) for RNA-seq using H3K27ac, H3K27me3 and H3K36me3 signals in genic promoters and intragenic enhancers and the interactions between them. This model was able to explain 67% of the gene expression variance (Supplementary Table 15), a percentage 5% higher than the 62% explained by a naïve model including the signals of all histone marks in all the components but no interaction between them (Supplementary Table 16), supporting that genic promoters and intragenic enhancers contained nearly all the epigenetic information needed to define gene expression levels in our data.

### Differential gene expression analyses

We identified genes with differential expression levels across species using the iDEGES/edgeR pipeline in the R package TCC (version 1.12.1)^64,65^ at an FDR of 0.1 and testing all species pairwise comparisons. Then, we determined the patterns of differential expression, species and direction of the gene expression change, using a two-step approach. For every gene, the Q-values obtained in species pairwise comparisons were ordered from smallest to largest. Different expression labels were then assigned to each species according to the ordered Q-values. Once all species had an assigned label, the average normalized expression values between groups were compared to determine the directionality of the change. We separate differentially expressed genes into two categories: genes with species-specific expression changes and gene with non-species-specific expression changes.

To investigate the relationship between changes in gene expression and changes in the regulatory architecture of a gene, for every type of regulatory component we run a Wilcoxon signed-rank test evaluating whether the number of regulatory elements with a given regulatory state in that particular regulatory component was significantly associated with higher expression levels, for strong and weak activities, or lower expression levels, for poised activities. P-values obtained for each regulatory role were corrected for multiple testing using the Benjamini–Hochberg procedure^62^.

### Over-representation analyses (ORA) of functional annotations

We defined sets of genes associated with fully conserved and species-specific component-epigenetic state combinations and explored their functional enrichments. To ensure the representativeness of the functional enrichments, for the gene lists associated with each type of component, we excluded genes associated with components with different epigenetic states activities (i.e., genes associated with both conserved strong and weak intragenic enhancers) or associated with both conserved and species-specific components levels (i.e, genes associated with both a conserved and a species-specific weak intragenic enhancer) and kept those gene lists with a minimum of 15 genes for enrichment analyses. Of note, orangutan-specific component-epigenetic state combinations were excluded from the analysis because they were defined using only one LCL replicate (see above) and they are likely to be enriched in inter-individual variable activities.

Over-representation of Gene Ontology (GO) terms was performed using the WebGestaltR function from the R package WebGestaltR (version 0.4.3)^66^ with minNum = 25 and remaining default options. This function controls the false discovery rate (FDR) by applying Benjamini-Hochberg procedure (default threshold FDR = 0.05)^62,67^. Previous analyses have shown that recent enhancers tend to occur in the same genes that already have highly conserved enhancers^9^. To control for the particular background of each component, we built different background gene sets including the set of human genes associated with at least one-to-one orthologous regulatory regions of each type of component, hence we have specific and different backgrounds for genic promoters, intragenic enhancers and promoter-interacting enhancers. To represent and compare enriched GO terms between component-state combinations, we performed a clustering of all significantly enriched GO terms using REVIGO^68^. We associated each GO term with the proportion of genes from each component-state combination that overlapped that GO term. In the case of GO terms enriched in more than one gene set, we chose the highest proportion of genes. We used this list as input for REVIGO. Given that REVIGO only reports the clustering of approximately 350 GO terms and our input list was larger than that, we used the R package GofuncR (version 1.8.0)^69^ to retrieve the parent GO terms of the remaining unassigned GO terms and add them to the corresponding group as defined by REVIGO. REVIGO group names were manually assigned, taking into account the most representative parent term (Supplementary Table 19).

### Analysis of tissue-specific gene expression patterns

We defined sets of human genes associated with fully conserved component-state combinations, and human genes associated with human-specific gains/losses of regulatory elements. Note that these gene lists are not mutually exclusive since a gene can we associated with different types of conserved or species-specific component-state combinations (e.g., a gene with both a human-specific intragenic enhancer with weak activity and a fully conserved intragenic enhancer with a strong activity). We obtained expression levels (median TPM values) across a collection of different tissues from the latest GTEx release (v8)^26^. We only included tissues with at least 70 samples and grouped tissue subregions into the same tissue category, as stated in Supplementary Table 18. For each component-state combination we followed a two-step approach to remove consistently low-expressed genes across tissues. For that we first assigned a value of 0 to all genes with a median expression level below 0.1 TPM and then we excluded from the analyses those genes that had an accumulated expression value in all tissues below 0.1xNumber of tissues (n = 29 tissues). For each component-state combination, differences in median expression across tissues were assessed with the Friedman test using the friedman.test function as implemented in R^55^. We used the Wilcoxon-Nemenyi-McDonald-Thompson test implemented in the pWNMT function of the R package NSM3 (version 1.14)^70^ to assess whether expression levels were significantly different for all pairwise tissue combinations. Then, we made use of the rank-biserial correlation to calculate the effect sizes for all statistically significant pairwise tests with the wilcoxonPairedRC function of the R package rcompanion (version 2.3.25)^57^.

We then evaluated the tissue-specificity of the genes associated with the different component-state combinations. For this we calculated the tissue specificity index^71^ (τ, tau) for each gene, which is defined as:

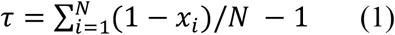

where *N* is the number of tissues and *x*_*i*_ is the expression value normalised by the maximum expression value. This value ranges from 0, for housekeeping genes, to 1, for tissue-specific genes (values above 0.8 are used to identify tissue-specific genes)^72^. Tissue-specificity indices were calculated for all genes included in the latest GTEx release^26^. Gene expression levels (median TMP) of grouped tissue categories (Supplementary Table 18) were normalised within and across tissues before calculating τ as implemented in the R package tispec (version 0.99.0)^73^. The calcTau function from this package provides a tau value for each gene and also a tau expression fraction for each tissue (which also ranges from 0 to 1) that indicates the specificity of a given gene for that tissue.

After calculating τ values, we compared their distributions between gene datasets with the Kruskall-Wallis test and assessed the significance of every pairwise comparison with the Dwass-Steel-Critchlow-Fligner test (dscfAllPairsTest function from the R package PMCMRplus version 1.4.4)^56^. Glass rank biserial correlation coefficient was used to compute the effect sizes associated with all statistically significant pairwise comparisons using the wilcoxonRG function from the R package rcompanion version 2.3.25^57^ (*P* < 0.05).

### Association of genes containing intragenic enhancers with signals of positive selection in humans

We built a database of human genomic regions with previously detected signals of positive selection in humans^27–29^ and selective sweeps in modern compared to archaic humans^30^. BEDtools^74^ was used to assign these regions to both protein-coding and non-coding genes following similar criteria to those used for building the gene regulatory architectures (Methods’ section *Classification of regulatory elements in different types of components of gene regulatory architectures*). We assigned these regions to a protein-coding gene if they were located within the gene or up to 5 Kb upstream of its TSS. Then, we made use of available interaction data for the cell line GM12878 (HiC^21^, HiChIP-H3K27ac^22^ and ChIA-PET^23^) to assign positively selected regions to their interacting protein-coding genes. We defined the 2,004 genes associated with at least one positively selected region as the set of genes with signals of positive selection in the human lineage. We computed the overlaps between this gene list and the lists of genes associated with the different component-state combinations. We used one or two-tailed Fisher’s exact test to assess the enrichment significance.

### Analyses of the density of human-fixed single nucleotide changes (hSNCs) in intragenic enhancers with weak enhancer states

In order to study the distribution of human-fixed changes in a specific type of regulatory element, we first generated a dataset with human-specific changes. We used sequencing data from a diversity panel of 27 orangutans, 42 gorillas, 11 bonobos and 61 chimpanzees^75–77^, as well as 19 modern humans from the 1000 genomes project^78^, all mapped to the human reference assembly hg19. We applied a basic filtering for each site in each individual (sequencing coverage >3 and <100), and kept sites where at least half of the individuals in a given species had sufficient data. Furthermore, at least 90% of the kept individuals at a given site in a given species had to share the same allele, otherwise the site was labeled as polymorphic in the population. Indels and triallelic sites were removed, and only biallelic sites were kept. We used data from a macaque diversity panel^79^, applying the same filters described above. The allele at monomorphic sites was added using bedtools getfasta^74^ from the macaque reference genome rheMac8. Since this panel uses the macaque reference genome, we performed a liftover to hg19 using the R package rtracklayer^80^ and merged the data with the great ape diversity panel.

Lineage-specific changes were retrieved as polymorphisms with sufficient information. Hence, human-specific changes (hSNCs) were defined as positions where each species carry only or mostly one allele within their respective population, the majority of individuals in each population have a genotype call at sufficient coverage, and the human allele differs from the allele in the other populations.

BEDtools^74^ was used to annotate those hSNCs in conserved or human-specific weak intragenic enhancers and the density of changes was calculated as the number of hSNCs present in each enhancer divided by the length of the enhancer.

To determine which human-specific intragenic weak enhancers were enriched in human-specific changes, we compared their density to what would be expected at random. For that, we first established the number of hSNCs that fall in human intragenic enhancers with weak enhancer states associated with 1-to-1 orthologous regulatory regions (our universe of enhancers). In each simulation, this number of mutations was randomly placed in this universe and we computed the density for each of the human-specific weak intragenic enhancers (10,000 simulations). With this approach, we corrected for the differences in the length of the enhancers. The P-value for each enhancer was computed as the number of simulations with a density equal to or above the observed density for that particular enhancer. All P-values were corrected by multiple testing using the Bonferroni method with the number of tests equal to the number of human-specific weak intragenic enhancers.

We then assessed whether the number of enhancers that were statistically enriched in hSNCs (or number of *hits*) was greater than what would be expected at random. In order to do that, for each enhancer we defined its mutation density critical value adjusting by multiple testing and using the simulated values. For example, in a hypothetical case of 100 enhancers and 10,000 simulations, for each enhancer we would order its simulated density of hSNCs from smallest to largest and take the 5th value as the critical one (given that our chosen alpha equals 5%, but it has to be corrected by 100 tests; therefore it becomes 0.05%). Once we established a critical value for each human-specific intragenic weak enhancer, we determined, for each simulation, how many enhancers had a density equal to or above their corresponding critical value. Finally, we computed the P-value comparing the number of artificial *hits* in each simulation with the number of observed hits.

## Acknowledgments

R.G.-P. was supported by a fellowship from MICINN (FPU13/01823). P.E.-C. was supported by a Formació de Personal Investigador fellowship from Generalitat de Catalunya (FI_B00122). M.K. was supported by a Deutsche Forschungsgemeinschaft (DFG) fellowship (KU 3467/1-1) and the Postdoctoral Junior Leader Fellowship Programme from “la Caixa” Banking Foundation (LCF/BQ/PR19/11700002). D.J. was supported by a Juan de la Cierva fellowship (FJCI2016-29558) from MICINN. T.M-B. is supported by funding from the European Research Council (ERC) under the European Union’s Horizon 2020 research and innovation programme (grant agreement No. 864203), BFU2017-86471-P (MINECO/FEDER, UE), “Unidad de Excelencia María de Maeztu”, funded by the AEI (CEX2018-000792-M), Howard Hughes International Early Career, Obra Social "La Caixa" and Secretaria d’Universitats i Recerca and CERCA Programme del Departament d’Economia i Coneixement de la Generalitat de Catalunya (GRC 2017 SGR 880). G.M., V.D.C. and L.D.C. were supported by grants from the Spanish of Economy, Industry and Competitiveness (MEIC) (BFU2016-75008-P) and G.M. was also supported by the “Convocatoria de Ayudas Fundación BBVA a Investigadores, Innovadores y Creadores Culturales”. J.L.G.-S. was supported by the Spanish government (grants BFU2016-74961-P), an institutional grant Unidad de Excelencia María de Maeztu (MDM-2016-0687) and the European Research Council (ERC) under the European Union’s Horizon 2020 research and innovation programme (grant agreement No 740041). A.N. was supported by Fondo Europeo de Desarrollo Regional (FEDER) with project grants BFU2016-77961-P and PGC2018-101927-B-I00 and by the Spanish National Institute of Bioinformatics (PT17/0009/0020). Figs. 1a, 2a, 4a and 6b fully or partially created with BioRender.com.

## Author contributions

T.M.-B. and J.L.G.-S. conceived the study; D.J. designed and supervised the analyses; L.D.C. supervised the work of G.M. and V.D.C.; A.B. procured non-human great ape cell lines; A.N. provided helpful insights; R.G.-P., G.M. and V.D.C. performed the experimental work; R.G.-P., P.E.-C., D.J., I.L., M.R. and M.K analyzed the data; D.J., R.G.-P., and P.E.-C. wrote the manuscript with input from all co-authors.

