## Supplementary_Methods_and_Figures for "Epigenomic profiling of primate LCLs reveals the evolutionary patterns of epigenetic activities in gene regulatory architectures"

This PDF file includes the description and/or content of:

**Supplementary Methods**

**Supplementary Figures 1 to 76**

**Supplementary Tables 1 to 27**

**Additional Files 1 to 5**

### EXPERIMENTAL METHODS

#### Cell line acquisition and cell growth

Lymphoblastoid cell line (LCL) GM19150 (Yoruban male) was purchased from the Coriell Institute. Chimpanzee, gorilla and orangutan LCLs were kindly provided by Dr. Antoine Blancher. Macaque LCLs were kindly provided by Dr. Gaby Dioxiadis.

Cell lines were grown in suspension at a confluency of 200,000 cells/mL to $1\times10^6$ cells/mL, at 37°C and 5% $CO_2$, in RPMI 1640 media (Invitrogen 42401-018) with 15% fetal bovine serum (FBS Invitrogen 10270-106), 1% penicillin-streptomycin (Invitrogen 15140-122) and supplemented with 2 mM L-glutamine (Sigma G7513-100ML).

#### DNA and RNA extractions

For DNA and RNA extractions, ~5 million cells were pelleted and washed with PBS (Sigma D8537).

DNA extractions were performed using a phenol-chloroform protocol. Briefly, cell pellets were resuspended in a lysis buffer (10mM Tris-HCl pH 8, Life Technologies 15568-025; 10mM NaCl, Sigma 71386; 0.2% IGEPAL CA-630, Sigma I8896) and incubated in ice for 30 minutes. Next, cells were washed and incubated at 37°C for 30 minutes in NEB2 buffer (New England Biolabs B7002S) supplemented with 10% SDS (Life Technologies AM9820) and 5µL of RNAse A (QIAGEN 79254), followed by an overnight incubation at 65°C with 10µL Proteinase K (Life Technologies AM2546). Two successive phenol-chloroform extractions were performed the day after. DNA was precipitated with NaAc 3M and washed twice with 70% and 100% EtOH, respectively. DNA integrity was checked in an agarose gel. Extracted DNA was used for WGS and WGBS libraries. Libraries were prepared using the TruSeq DNA PCR Free Library Preparation Kit and TruSeq DNA Methylation Kit, respectively, following Illumina's standard protocol. 150 bp paired-end (150PE) reads were sequenced in a HiSeqX machine.

RNA was extracted using the miRNeasy Mini kit (QIAGEN 217004) following manufacturer's instructions. The TruSeq Stranded Total LT Samples Prep Kit with Ribo-Zero was used for library construction, following Illumina's standard protocol. Library preparation and sequencing were performed at Macrogen (Seoul, Korea). 101 bp paired-end reads (101PE) were sequenced in a HiSeq4000 machine.

### Chromatin Immunoprecipitation

For ChIP-seq experiments, LCL primate cell lines were harvested and washed with PBS (Sigma D8537). Cells were fixed by incubating in 1.5% formaldehyde (Sigma F8775) PBS buffer at room temperature for ten minutes. Formaldehyde was quenched with 125mM glycine (Invitrogen 15527-013) at room temperature for five minutes, after which the samples were placed on ice for 15 minutes. Cells were then washed twice in cold PBS, aliquoted and pellets of ~20 million cells were frozen at -80°C. ChIP experiments were carried out following a standard protocol in the lab[1]. Briefly, crosslinked cell pellets were resuspended in 500 µL ice-cold ChIP buffer (50mM Tris-HCl pH8, 100mM NaCl, 5mM EDTA, 0.33% SDS, 1.67% Triton X-100) with Protease inhibitors (cOmplete™ EDTA-free, SIGMA) and 1mM PMSF. Lysates were sonicated in a Bioruptor (Diagenode) to obtain an average fragment size of 250 bp and centrifuged for 20 minutes at full speed. An aliquot of the soluble chromatin was taken, reverse cross-linked, purified using the Qiagen PCR purification kit (Cat No./ID: 28104) and quantified by Nanodrop. For each ChIP, 30 µg of soluble chromatin, 0.75 µg of mouse E14TG2a (as spike-in control) and 5 µg of the corresponding histone modification antibody were incubated overnight at 4°C. ChIP DNA was recovered by incubating with 42 µL protein A bead slurry (Diagenode) for 2 hours, washed three times with low salt buffer (50mM HEPES pH7.5, 140mM NaCl, 1% Triton X-100) and one time with high salt buffer (50mM HEPES pH7.5, 500mM NaCl, 1% Triton X-100). DNA complexes were decross-linked at 65°C overnight with proteinase K (Invitrogen) and DNA was purified using the PCR purification kit (Qiagen, 28104) and quantified by Nanodrop. We used commercially available antibodies against H3K4me3 (Diagenode, C15310003), H3K4me1 (Abcam, ab8895), H3K36me3 (Abcam, ab9050), H3K27me3 (Millipore, 07-449) and H3K27ac (Millipore 07-360).

A total of 63 ChIP-seq libraries were constructed: 45 libraries for the ChIP-seq experiments (one per sample and histone modification) and 18 libraries for Input samples (Input1 for histones H3K4me3, H3K4me1, H3K36me3 and H3K27me3; Input2 for H3K27ac). Prior to library preparation, ChIP-qPCR validations were performed using primer pairs of known active and repressed genes in human LCLs (active: CEP250, GAPDH, Beta-Actin; repressed: Sox2) and a negative intergenic region as control for the histone modifications studied. 2-10ng of ChIP DNA were used to prepare the sequencing libraries using the NEBNext Ultra DNA Library Prep Kit for Illumina (NEB, E7370L) as per manufacturer's instructions. ChIP-seq libraries were size selected to remove fragments below 100 bp and amplified for 10 PCR cycles. 4 to 5 libraries were pooled and loaded into a total of 13 lanes. 50 bp single-end reads (50SE) were sequenced in an Illumina HiSeq2500 machine, using v4 chemistry.

### Assay of transposable chromatin

ATAC-seq libraries for nine lymphoblast cell lines were generated following the Buenrostro et al. protocol[2]. Briefly, 50,000 cells were harvested, washed in cold PBS and resuspended in 50mL of cold lysis buffer (10 mM Tris-HCl pH 7.4, 10 mM NaCl, 3 mM $MgCl_2$, 0.1% (v/v) Igepal CA-630). Samples were spun down for 10 min at 500g, 4°C. Pellets were resuspended in the transposition reaction mix and incubated at 37°C for 30 minutes. Samples were purified using the Qiagen MinElute PCR purification kit. Transposed DNA was eluted in 10µL of elution buffer and subjected to PCR amplification for 8 cycles using barcoded primers and NEBNext High Fidelity PCR master mix. ATAC-seq libraries were purified using 1.8X volumes of AMPure XP beads to remove fragments below 100 bp. Library quality was assessed using a Bioanalyser High Sensitivity DNA analysis kit (Agilent). 50 bp paired-end reads (50PE) were sequenced on a HiSeq 2500 platform (Illumina).

### Data sources for GM12878 (H1)

We retrieved data for cell line GM12878 from different sources: genome sequencing data was obtained from the Garvan Institute (http://www.garvan.org.au/research/kinghorn-centre-for-clinical-genomics/clinical-genomics/sequencing-services/sample-data); methylation data (GSM2308633) was obtained from ENCODE (https://www.encodeproject.org/experiments/ENCSR890UQO/). RNA-seq

(SRR998197, SRR998198) and ChIP-seq (H3K4me3: SRR998194; H3K4me1: SRR998191; H3K27ac: SRR998178; H3K36me3: SRR998187; H3K27me3: SRR998189; Input: SRR998196) data were obtained from Kasowski et al.[3]. ATAC-seq data (SRR891270) was obtained from Buenrostro et al[2].

### COMPUTATIONAL METHODS

#### Quality control

Sequence quality was assessed using the FASTQC software[4] for all types of generated sequencing data. Quality filtering and adapter removal were performed when deemed necessary using Cutadapt[5] to ensure that raw reads were within standard parameters. We constructed species-specific mappability tracks for the different sequencing read lengths included in this study (151-kmers, 101-kmers and 50-kmers) in order to identify positions in the reference genomes where reads could not be confidently mapped. We used the 'gem-mappability' module from gemtools/1.7[6], which uses a kmer approach to detect genomic regions that are duplicated and, therefore, likely to be problematic. For paired-end data, we considered mappable regions those with up to 2 mismatches (threshold $\geq 0.5$). For single-end data, we considered mappable regions those with less than 2 mismatches (threshold $> 0.5$)

#### ChIP-seq and ATAC-seq short-read alignment and peak calling

ChIP-seq 50 bp SE reads were aligned to the corresponding reference genome along with the genome of the Epstein-Barr virus with Bowtie2 (version 2.2.11)[7] using the default 'sensitive' settings. Low-quality and multiple-mapping reads were removed using Samtools (version 1.2)[8] with options '-q30 –F 1796' ('-q30 -F 1804 -f 2' for sample GM12878 that was sequenced with 75 bp PE reads). PCR duplicates were removed using Picard (version 1.95) (http://broadinstitute.github.io/picard/). Reads mapping to the corresponding species genome were then selected and only autosomal chromosomes were kept for downstream analysis. We identified enriched regions or peaks with MACS2[9] using the following parameters *-nomodel --shift 0 –extsize sample_specific_fragment_size --keep-dup all* for histones H3K4me3, H3K4me1 and H3K27ac, and adding *–broad* for histones H3K36me3 and H3K27me3. We only kept reproducible peaks, which we defined following a partition concordance approach. For each sample, we created two pseudo-replicates (defined by randomly choosing half of each sample reads without replacement) and used them to call peaks using the above-stated command. We defined as reproducible peaks those that overlapped at least 50% with peaks from both sample pseudo-replicates. Samples peaks were filtered based on their corresponding 50-kmer mappability tracks and only peaks with at least 80% mappable bp were retained. Histone peaks were strongly enriched in genomic regions with consistent chromatin states (Supplementary Figs. 2 and 66).

ATAc-seq 50 bp PE reads were mapped to the corresponding reference genome along with the genome of the Epstein-Barr virus using Bowtie2 (version 2.2.11)[7] with default 'sensitive' settings and '--maxins 10000' that allows the mapping of reads with a maximum insert size of 10 kbp. Low-quality and multiple-mapping reads were removed using Samtools (version 1.2)[8] with options '-q30 -F 1804 -f 2'. PCR duplicates were removed using Picard (version 1.95) (http://broadinstitute.github.io/picard/). Only autosomal chromosomes were kept for downstream analyses. Open chromatin regions were called using the ENCODE ATAC-seq pipeline (https://encode-dcc.github.io/wdl-pipelines/), with an IDR threshold of 0.1. Open chromatin regions were further filtered based on 50-kmer species-specific mappability tracks and only accessible regions with at least 80% mappable bp were retained. We applied the same approach as before to retain reproducible open chromatin regions. Open chromatin regions were strongly enriched in genomic regions with consistent chromatin states (Supplementary Figs. 2 and 66).

### Background noise normalization of enrichment values

To obtain background noise corrected histone enrichment signals, we implemented a 3-step approach. First, sample matched immunoprecipitated (IP) and input counts were normalized by sequencing depth. The median number of aligned reads across samples IP and input datasets (for any given histone mark) was used as the total reference count. IP and input counts were then scaled to the total reference count. Second, we removed background noise in a sample- and histone modification-specific fashion. The input-IP relationship in genomic regions with no significant enrichment (non-peaks) was used to estimate the noise contribution to the IP in peaks. We defined a confident set of non-peaks with the same size and same mappability requirements as for the identified samples peaks that did not overlap with any peak detected in any of the five species considered. These strict requirements prevented the inclusion of putative false negatives, enriched regions that were overlooked by the peak calling algorithm. We used a Deming regression to evaluate the linear relationship between the input and IP in non-peaks. Deming regression assumes random measurement error in both the dependent y-variable and the independent x-variable[10], and thus, is more suitable for these analyses than simple linear regression where only the response variable Y is measured with error (Supplementary Fig. 67). The noise contribution to the IP was estimated for each genomic region of interest using the inferred linear input-IP relationship in non-peaks with the corresponding depth-normalized input count (Supplementary Fig. 67). Then, the estimated noise is subtracted from the depth-normalized IP counts in each genomic region, resulting in the background-noise normalized IP enrichment. Finally, the normalized enrichment signal was transformed with the inverse hyperbolic sin ($\boldsymbol{asinh}$):

$$asinh = ln ( x + \sqrt{x^2 + 1})$$

This transformation makes the data homoscedastic, that is, with approximately equal spreads despite pronounced variations in enrichment levels. This eases the handling and interpretation of the data and is fundamental in the regression models that will be implemented later on.

### RNA-seq alignment and gene expression quantification

RNA-seq 101bp PE reads were aligned to the corresponding reference genome along with the genome of the Epstein-Barr virus using hisat2 (version 2.0.4)[11] with default parameters. Low-quality and multiple-mapping reads were removed using Samtools (version 2.1)[8] with options '-q30 -F 1804 -f 2'. Potential PCR duplicates were removed using PICARD v1.91 (http://picard.sourceforge.net). Read counts in genes were computed using htseq-count[12]. We used the R package DESeq2 (version 1.14.1)[13] to evaluate the coherence between technical replicates, exploring sample-to-sample distances and PCA results. Technical replicates were highly correlated and hence collapsed and used thereafter (Supplementary Fig. 68 and Supplementary Table 6). Gene quantification on merged replicates was performed using STRINGTIE (version 1.3.3)[11] based on species-specific Ensembl version 91[14] gene annotations. Estimated gene expression levels in TPM (transcripts per million transcripts) were obtained from the STRINGTIE/1.3.3.

### Establishment of orthologous relationships

To find 1-to-1 orthologous genes we mined the Ensembl[14] database was using biomaRt[15]. We retrieved the following features for the non-human species studied: Ensembl genes ID, gene coordinates, homology type and orthology confidence. Only autosomal one-to-one orthologous genes with the highest orthology confidence value (1) were kept for defining the set of 1-to-1 orthologous genes across the five species (11,249 genes). Of these, 9,936 were annotated as protein-coding in all species and 7,850 were expressed (TPM $\geq$ 0.5) in at least one species.

To find orthologous regulatory regions, we followed the approach described in Supplementary Fig. 69. First, we mapped non-human regulatory elements to the human reference genome and considered orthologous regulatory elements those with a minimum overlap of 50% and for which the overlapping regions had similar size (50% of each other) (Supplementary Fig. 69a). In the case of several overlapping regions, we prioritized the regions with the largest overlap. For every orthologous region for which we could not recover an orthologous regulatory element in at least one species, we defined the primate orthologous regulatory region coordinates as a genomic region of size equal to the average size of the species orthologous elements and located in the middle of the collapsed orthologous elements coordinates. We used this primate orthologous regions to recover overlooked orthologous relationships, this is, regulatory elements that either overlapped or were close (200 bp) to the primate orthologous region (Supplementary Fig. 69b). If no regulatory element was recovered, the coordinates of the primate orthologous region were mapped to the corresponding species assembly (Supplementary Fig. 69c). For species-specific regulatory elements for which we found no orthologous regulatory elements in any of the other species, we mapped their coordinates to the other species reference genome assemblies and considered those regions as the corresponding orthologous region (Supplementary Fig. 69d). All inter-species projections were performed using the liftOver tool from the UCSCTOOLS/331 suite[16]. We used pairwise best reciprocal chains. For every inter-species projection, coordinates were mapped twice, going forward and backward, and only regions that could be properly mapped in both directions were kept. Overlaps were performed using the intersectBed tool from the BEDTools suite[17].

Note here that our dataset of orthologous regulatory regions is restricted to genomic regions that can be mapped across species (regions for which an orthologous region at the sequence level can be found). This is a restrictive protocol that ensures the comparability and balance between different species, which have very different levels of annotation and genome assembly qualities. However, this comparability comes at the price of not being able to study the epigenomic changes associated with genomic gains and losses.To assign a regulatory state to those orthologous regions not associated with a regulatory element we used the underlying enrichments in histone modifications and open chromatin (see Assignment of a regulatory state to regulatory elements). For those analyses comparing the regulatory state at orthologous regulatory regions associated with genes, we assigned a consensus regulatory component type to each orthologous regulatory region considering the different quality of the assemblies and gene annotations. Specifically, we assigned to the group of orthologues the type of regulatory component assigned in more species. In the case of a draw, we established the following hierarchy: human > chimpanzee > macaque > gorilla > orangutan.

### Normalization of gene expression and enrichment signals across species

First, we used batch correction to remove the technical variation derived from the integration of previously published data for the cell line GM12878. The batch effect was corrected using the ComBat function implemented in the R package sva (version 3.22.0)[18]. Then, we developed a method to normalize gene expression across samples. This method is based on the identification of a set of internal reference controls (IRCs) (genes) whose expression is constant across samples and from which a normalization factor can be derived (Supplementary Fig. 70). We defined internal reference controls as genes that were not outliers in any pairwise sample comparison, were outliers were defined as any gene with pairwise sample differences larger than the pairwise sample-specific median difference plus two median absolute deviations. We first standardized (robust standardization) each sample signal in IRCs. Then, we defined the representative IRCs values as the means of the standardized values of each IRC in the different samples. These representative IRCs are used to define a common scale for all the samples. We modeled the linear relationship between IRCs of each sample and the representative IRC values using Deming regressions. Sample-specific normalization factors are the result of dividing each sample-specific slope by the mean slope of all samples. To normalize the signal of all genes, we first scaled each sample signal in the robustly standardized distribution of IRCs using the corresponding sample median and median absolute deviation computed using these standardizations. Then, we proceeded to do the normalization with the corresponding sample normalization factor. Finally, to retrieve normalized signals, destandardization was performed using the mean and standard deviation of the samples' IRC median and median absolute deviations.

To assess the performance of the developed calibration method, we evaluated the effect of both the batch correction and expression signal normalization for every sample-pairwise comparison. To do so, we measured the angle to the identity line (x=y). If the normalization procedure removed noisy variance effectively, differences between samples were expected to be reduced. This would reflect in regression lines closer to the identity line (Supplementary Fig. 71). Our method effectively removed technical noise in the expression signal, the angles were successfully shrunk both at IRCs and pairwise orthologous expressed protein-coding genes excluding IRCs (Supplementary Fig. 72).

We further compared our normalization method with the widely used quantile normalization[19] and found that our method outperformed the latter particularly at normalizing the tails of the distributions (Supplementary Figs. 73 and 74). Normalization of the tails is a known problem associated with quantile normalization, which cannot properly handle the greater inter-samples differences at extreme values. Our parametric approach overcomes this limitation and correctly calibrates signal values throughout the whole distribution.

We applied to the same method to normalize the enrichment signals of the histone modifications at regulatory elements associated with orthologous protein-coding genes.

### Analysis of whole-genome sequencing data (WGS)

#### WGS mapping

151 bp PE reads were mapped to the corresponding species reference genome assemblies: human (hg38), chimpanzee (panTro5), gorilla (gorGor4), orangutan (ponAbe2) and macaque (rheMac8) along with the reference genome of the Epstein-Barr virus. Read alignments were carried out using BWA-MEM (version 0.7.8-r455)[20] with default parameters. Low-quality and multiple-mapping reads were removed using Samtools v1.6[8] with options '-q30 -F 1804 -f 2' and potential PCR duplicates were removed using PICARD v1.91 (http://picard.sourceforge.net).

#### Genotyping

WGS was used for genotyping every cell line and hard-filtering criteria were applied to identify SNPs, Indels and STRs. Variant discovery was performed using GATK version v3.7[21]. 'HaplotypeCaller' was run for each sample independently with default parameters using the '--emitRefConfidence GVCF' mode. Subsequently, both biological replicates were jointly genotyped using 'GenotypeGVCFs'. Hard filtering was conducted following GATK's recommendations. For SNPs, positions were removed if: 'QD < 2', 'MQ < 40', 'FS > 60', 'SOR > 3'. For Indels, positions were removed if: 'QD < 2', 'FS > 200', 'SOR > 10'. We further filtered variants based on their sample-specific callability, determined with the tool 'CallableLoci' with the following parameters: --minDepth 4, --maxDepth 100. Finally, mappable variants were kept based on 151-kmers species-specific mappability tracks (Additional File 5).

We performed principal component analysis (PCA) with EIGENSOFT (version 7.2.1)[22] using single nucleotide variants on chromosome 21 to evaluate how our samples related to the great ape individuals genotyped in the GAGP2[23]. Variants were mapped to hg38 coordinates and chromosome 21 variants were selected for the analysis. For the mitochondrial PCA, sequences from several studies were compiled to create a mitochondrial reference panel (Additional File 3). Each sequence was mapped to hg38 mitochondrial sequence with bwa-mem[9] and variants were called with VarScan2[24]. The obtained clusterings confirmed the species of the samples included in this study (Supplementary Fig. 75) and further allowed us to infer the corresponding subspecies of individual non-human great ape samples (Additional File 2). Macaque samples were of known Burmese origin. In addition, the sample-specific X chromosome coverage was used to confirm the sex of every sample included in the study (Additional File 3). The number of reported SNPs, Indels and STRs fall within the expected range considering the species-specific genomic diversity and the inferred subspecies (Additional File 2). Short tandem repeats (STRs) were called using HipSTR[25] with default stutter models and setting to 15 the minimum number of reads required to genotype a locus. STR filtering was performed using the recommended parameters: --min-call-qual 0.9 --max-call-flank-indel 0.15 --max-call-stutter 0.15 --min-call-allele-bias -2 --min-call-strand-bias -2 (Additional File 5).

### Analysis of whole-genome bisulfite sequencing data (WGBS)

Bisulfite converted sequencing data was used to estimate CpG methylation values genome-wide. Individual CpG methylation levels are inferred as the fraction of aligned methylated reads. In this regard, it is important to note the impact of nucleotide variants, genomic positions where the reference genome and the sequence of the sample under study differ. SNPs can introduce biases due to incorrect estimation of the methylation state: heterozygous loci might be assigned intermediate methylation values, whereas homozygous sites would result in an unmethylated call. Matching whole-genome sequence data allows the avoidance of this bias.

151 bp PE reads were mapped to the *in silico* bisulfite-converted species reference genome assemblies along with the *in silico* bisulfite-converted genome of the Epstein-Barr virus using Bismark (version 0.16.1)[26]. Low-quality and multiple-mapping reads were removed using Samtools (version v1.2)[8] with options '-q30 -F 1804 -f 2'. Potential PCR duplicates were also removed using Bismark's deduplicate_bismark program. Inspection of M-bias plots[27] revealed a methylation bias towards the end of the reads, so measurement at the last 15 positions was excluded from further analysis.

Custom Bash scripts were used to filter mappable CpG sites based on 151-kmers species-specific mappability tracks. Individual CpG methylation levels were summarized for each sample computing the ratio of unmodified Cs (methylated) to bisulfite converted Ts (unmethylated) using the R package bsseq (version 1.8.2)[28]. Only autosomal CpGs with a minimum of 4x coverage were considered for downstream analysis.

On average, a CpG coverage of ~16X was achieved after filtering CpGs located in non-mappable species-specific regions, removing sample-specific identified polymorphic CpGs and setting a minimum threshold of 4 sequencing reads. Methylation values were then inferred in approximately 17.4 million CpG sites per sample. Overall, the five species exhibited similar levels of CpG methylation, with an average value of ~61%. These values are expected and comparable to previous estimates in LCLs[29], which are, in general, lower than those observed in non-transformed tissues, e.g., ~72% global methylation in great-ape blood[30]. Nonetheless, orangutan sample O1 stood out, due to its unusually low global methylation (average sample-specific methylation value of 48.6%). As a result, Pearson correlation values between orangutan replicates were much lower (64%) than among human (81%), chimpanzee (77%), gorilla (77%) and macaque (81%) samples.

The R package MethylseekR (version 1.12.0)[31] was used to identify unmethylated regions (UMRs) and low methylated regions (LMRs). Methylation levels and FDR parameters were inferred from the data, as suggested by the MethylseekR workflow.

UMRs tend to be CpG-rich regions and are commonly regarded as proximal regulatory elements; LMRs, CpG-poor, have been associated with the binding of transcription factors, which cause the local reduction of otherwise high methylation levels, and are largely considered as distal regulatory elements, highly dynamic and tissue-specific[32]. Overall, similar numbers of UMRs and LMRs were found across samples and species (Supplementary Fig. 76). Notably, sample O1 showed a discordant pattern, with particularly elevated numbers of both UMRs and LMRs.

The relationship between UMRs and LMRs and chromatin states was explored, and the expected trends were observed. On the one hand, UMRs showed an enrichment pattern highly akin to that of CGI. On the other hand, LMRs overlap with active intergenic enhancer states (chromatin states E5, E6 and E7, Supplementary Figs. 2 and 66 and Additional File 4).

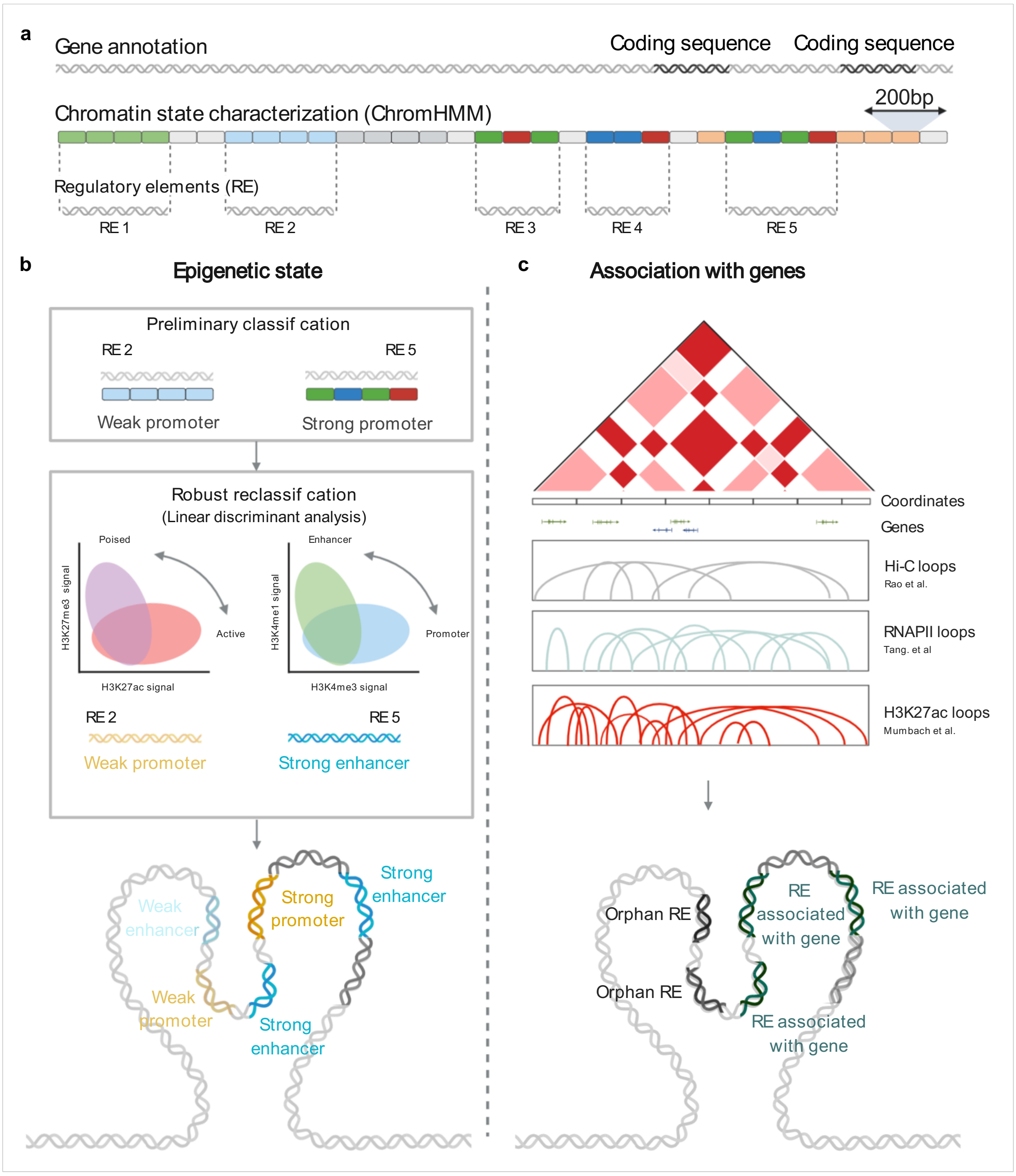


**Supplementary Figure 1. Schematic illustration of the approach followed to annotate and classify regulatory elements. a,** DNA strand represents a gene annotation track, wherein dark grey regions correspond to coding annotated regions. The second row represents the binarized output from ChromHMM, wherein each box corresponds to a 200 bp bin. Light grey indicates bins without evidence of promoter or enhancer states, whereas the different colors represent the different learned chromatin states (Methods, Supplementary Fig. 2). Shorter DNA strands represent the genomic coordinates defined for regulatory elements that result from merging adjacent 200 bp bins with epigenetic signals associated with promoter or enhancer states. We defined species regulatory elements from the union of the regulatory elements detected in each biological replicate (Methods). **b,** We established a hierarchy between chromatin states based on the combination of chromatin marks found within each regulatory region and classified regulatory elements into epigenetic promoter (P) and enhancer (E) states with three different activity levels: strong (s), weak (w) or poised (p) (Methods). Then, we applied a linear discriminative analysis (LDA) (Methods) using normalized histone and open chromatin enrichments to refine this epigenetic classification (Supplementary Methods). **c,** We linked regulatory elements to genes based on gene proximity and using previously published 3D chromatin maps in GM12878 cells we recovered physical interactions between regulatory elements. We additionally assigned regulatory elements associated with genes to a type of regulatory component. We classified regulatory elements into genic promoters (gP), genic enhancers (gE), proximal enhancers (prE), promoter-interacting enhancers (PiE) and enhancer-interacting enhancers (EiE) (Fig. 4a).

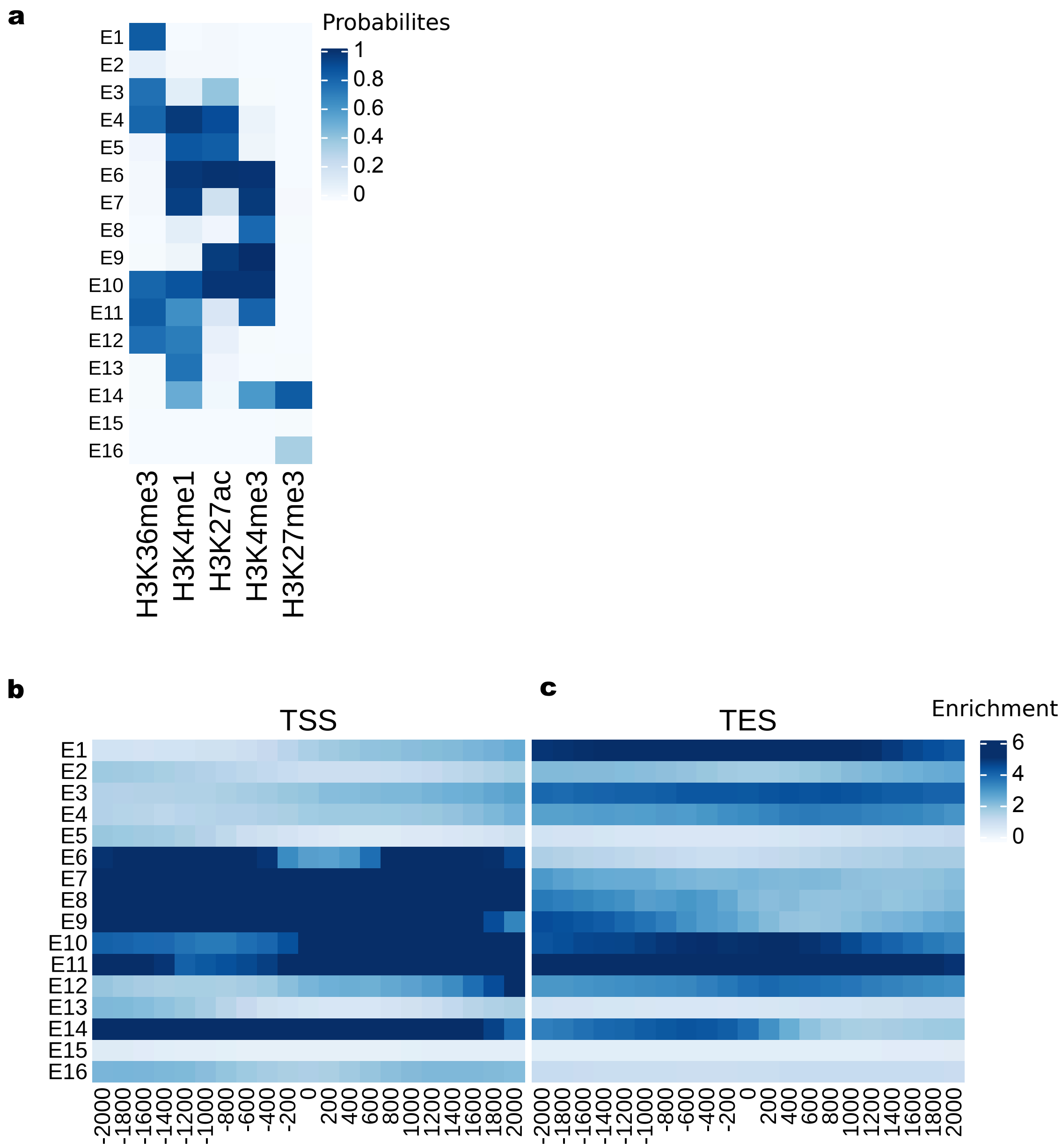


**Supplementary Figure 2. Chromatin states are enriched in biologically meaningful genomic regulatory regions.** Each row corresponds to one of the 16 chromatin states jointly learnt using the combined information of the histone marks H3K4me3, H3K4me1, H3K27ac, H3K36me3 and H3K27me3. **a,** Emission parameters form ChromHMM. The darker the blue the higher the probability of observing a given mark in a given state. **b** and **c,** Overlap and neighborhood enrichment analyses in functionally defined regions. Cell values represent the average fold enrichment across samples over several different functional annotations **(b)** and distances from transcriptional start and end sites (TSS and TES) **(c).** The 16 chromatin states recovered major regulatory regions including active promoters (E8, E9) and enhancers (E6) states, flanking upstream and downstream promoters (E7, E11) and enhancers (E5, E10), bivalent states (E14), elongation states (E1 and E2), heterochromatin (E16) and low signal (E15). Chromatin states did not reach the required resolution to distinguish between poised promoters and enhancers, although both types of regulatory elements are present in the data (E14). Cell values represent the average fold enrichment across samples.

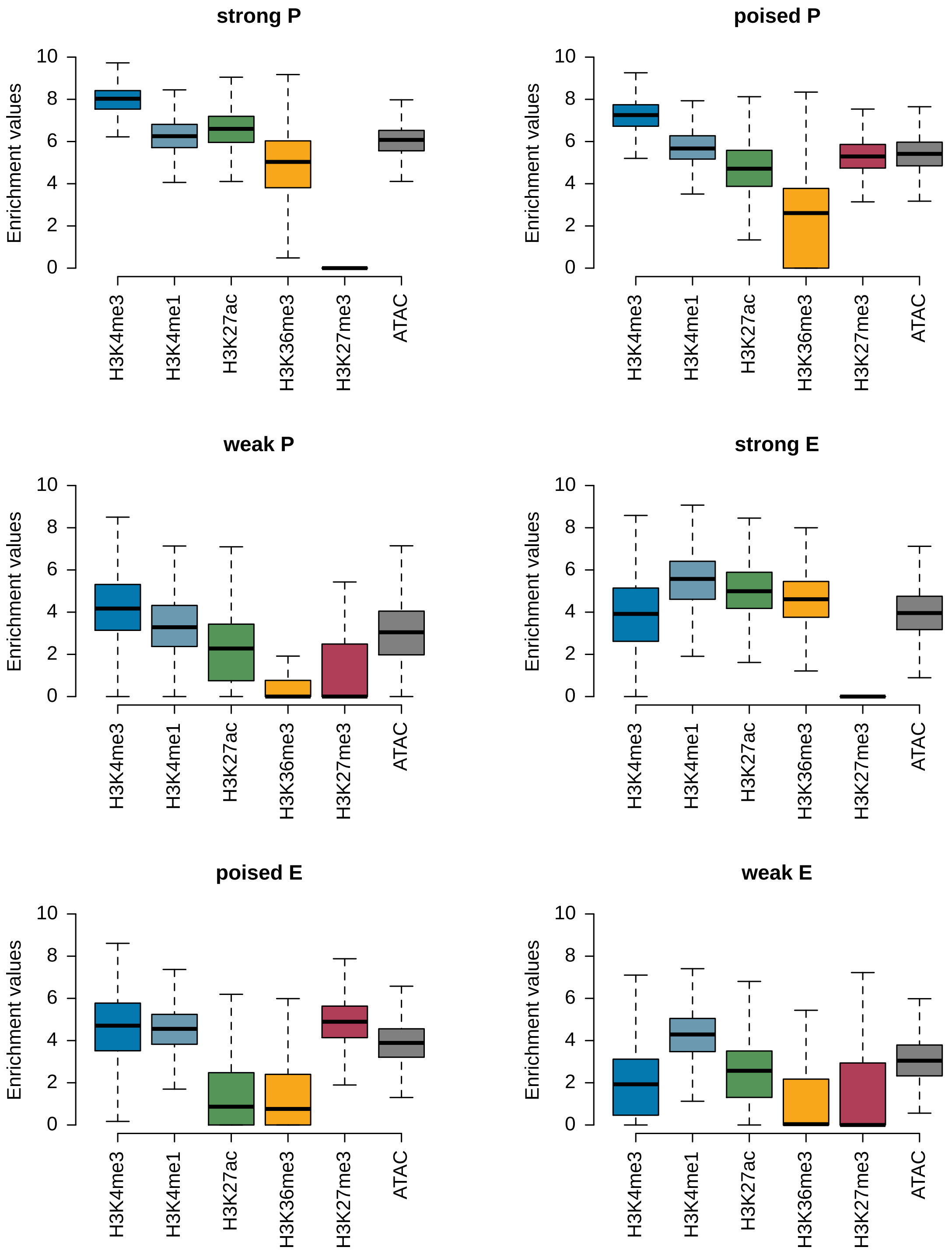


**Supplementary Figure 3. Distribution of normalized enrichment values associated with histone marks and open chromatin signals in each regulatory state.**

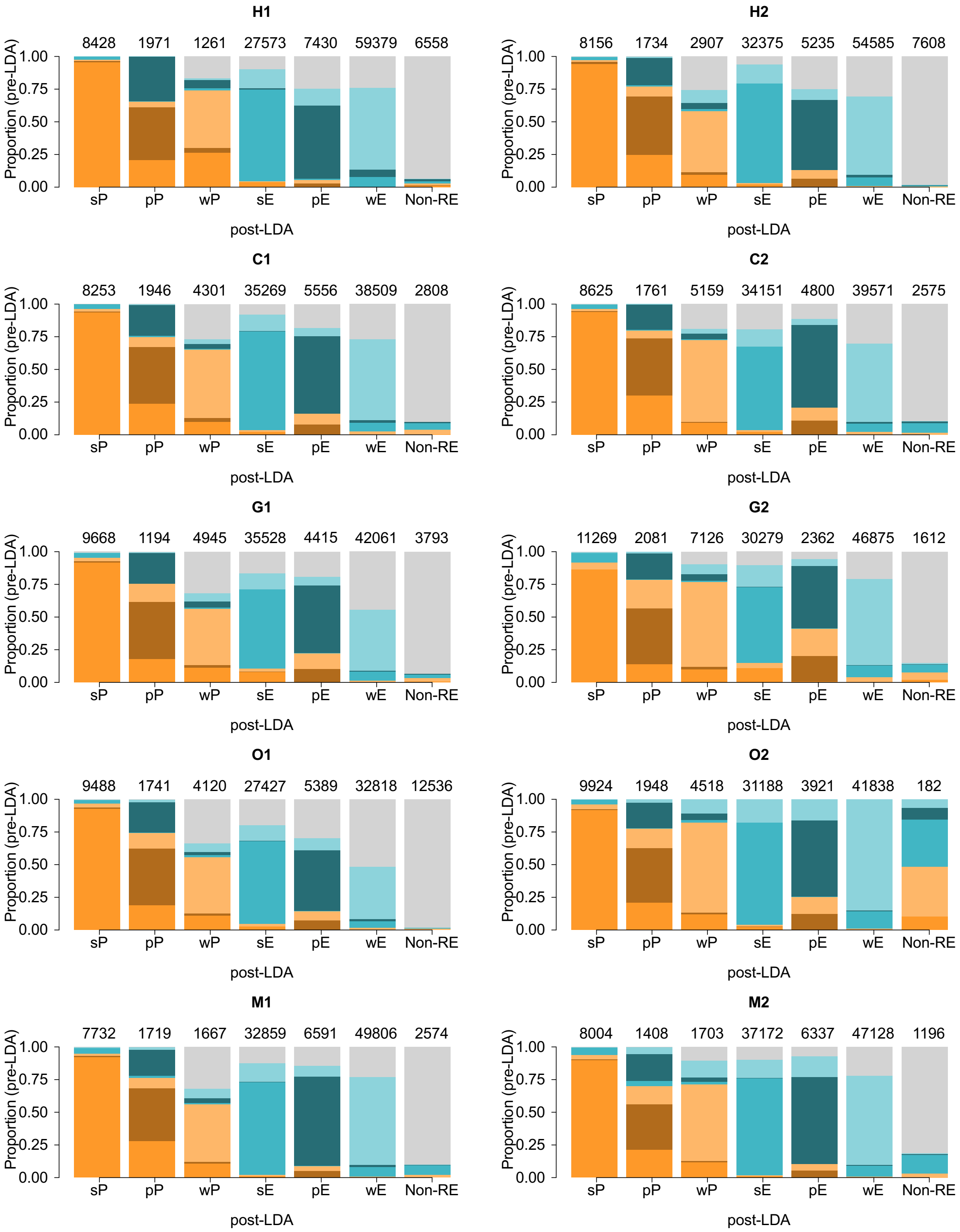


**Supplementary Figure 4. Linear discriminant analyses (LDA) of regulatory elements.** Each barplot shows, for regulatory elements, assigned the x-axis regulatory state through the LDA analysis, the proportion of pre-LDA regulatory states. Numbers on top of the bars indicate the number of regulatory elements with each regulatory state in every sample.

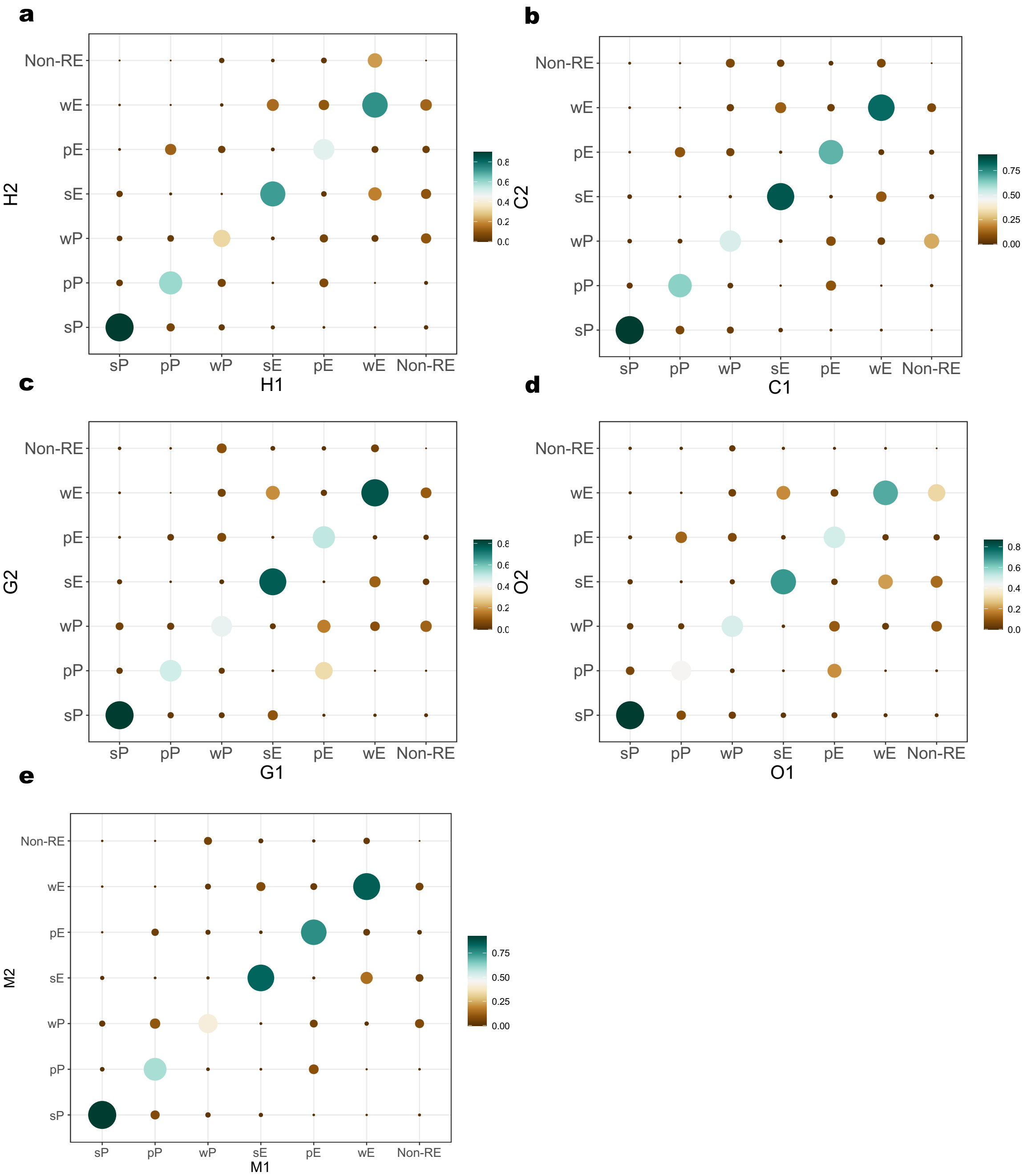


**Supplementary Figure 5. Similarity in the regulatory states assigned to regulatory elements between biological replicates.** To evaluate the similarity in the regulatory states between biological replicates, we calculated the Sørensen-Dice similarity coefficient (DSC) as DSC=2·x∩yx+y wherein x and y are the numbers of regulatory elements with a given regulatory state in each of the replicates. **a** to **e** Similarity between human, chimpanzee, gorilla orangutan and macaque replicates, respectively.

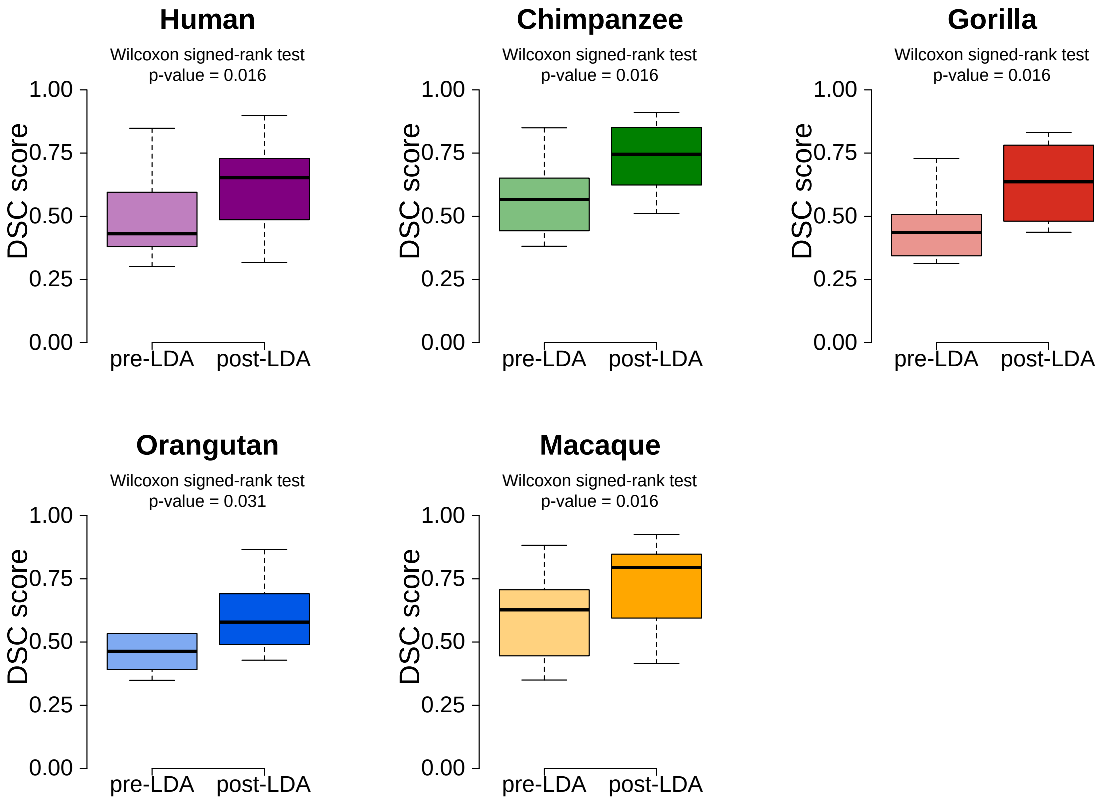


**Supplementary Figure 6. Comparison of the similarity between biological replicates before and after the linear discriminant analysis (LDA).** Distribution of the DSC scores corresponding to regulatory elements with the same regulatory state in both biological replicates (diagonal from Supplementary Figure 4).

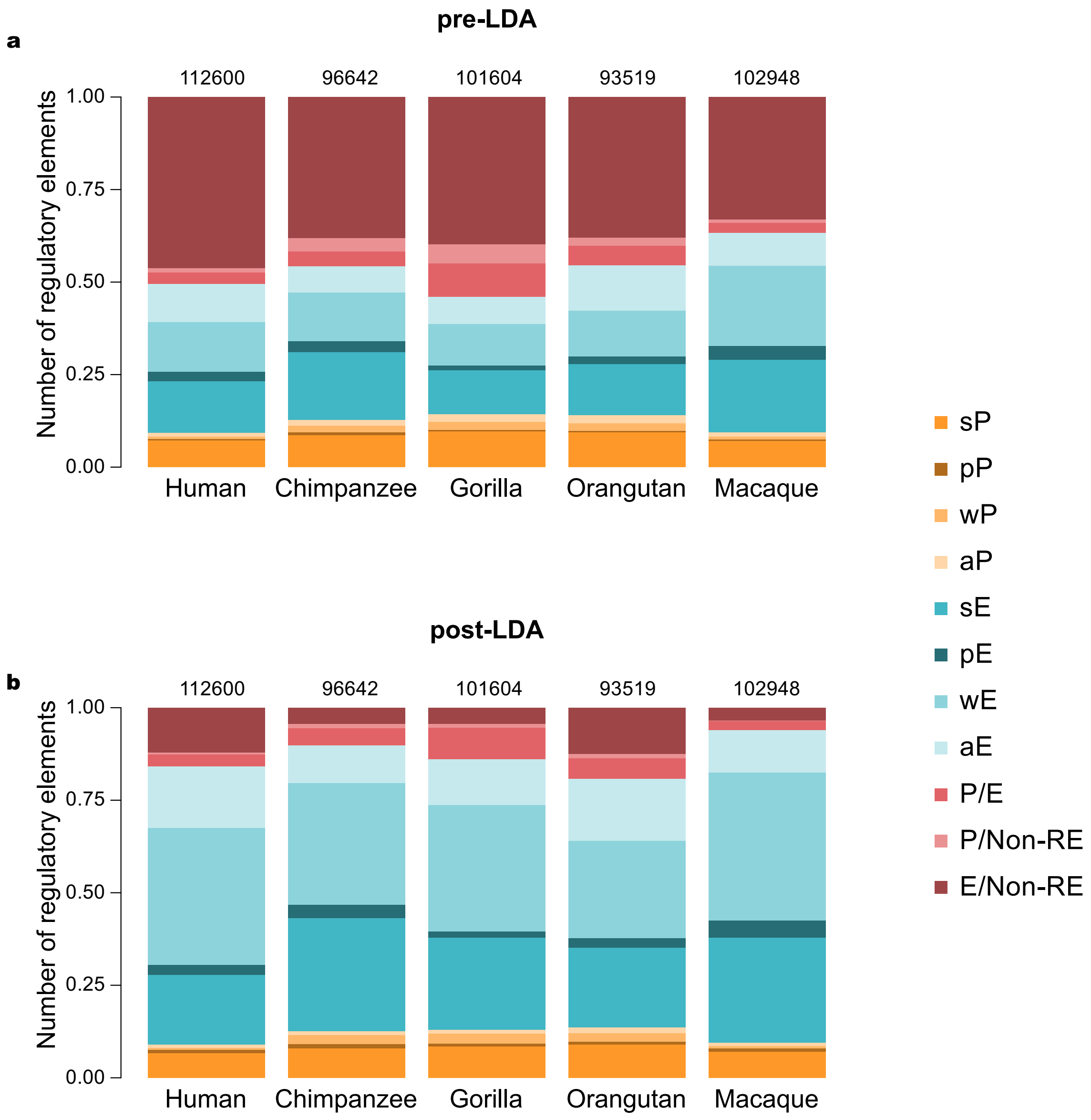


**Supplementary Figure 7. Comparison of the regulatory state of the regulatory elements before and after the linear discriminant analysis (LDA).** Each barplot shows the proportion of regulatory elements with a given regulatory state. Numbers on top of the bars indicate the total number of regulatory elements annotated per species. Color-coded the different regulatory states **a,** pre-LDA composition **b,** post-LDA composition. aP and aE corresponded to regulatory elements with either a promoter or enhancer state but with different activities between biological replicates. P/E correspond to regulatory elements with a promoter state in one replicate and an enhancer state in the other replicate. P/Non-RE and E/Non-RE correspond to regulatory elements with a promoter or enhancer state in one replicate and no evidence of regulatory activity in the other replicate.

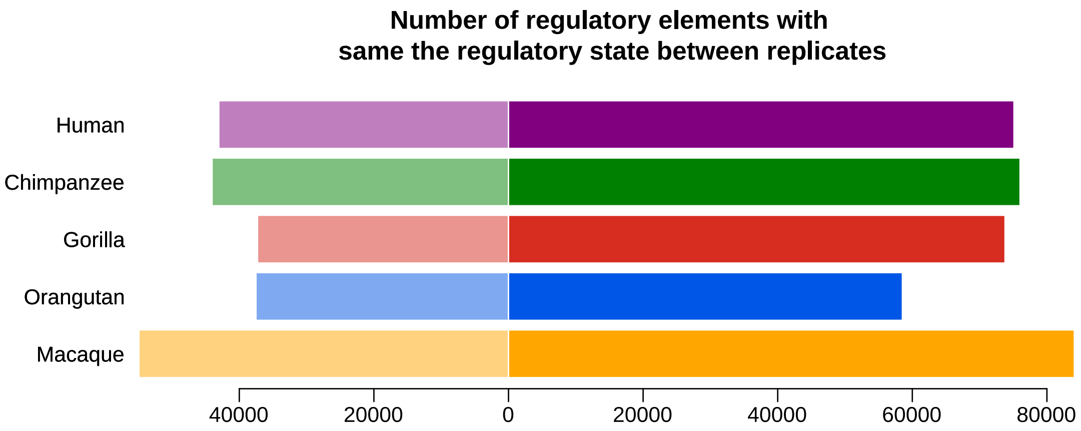


**Supplementary Figure 8. Linear discriminant analyses increased the number of regulatory elements with the same regulatory state.** Left and right barplots show, for every species, the number of regulatory elements with the same regulatory state before(left) and after (right) the LDA.

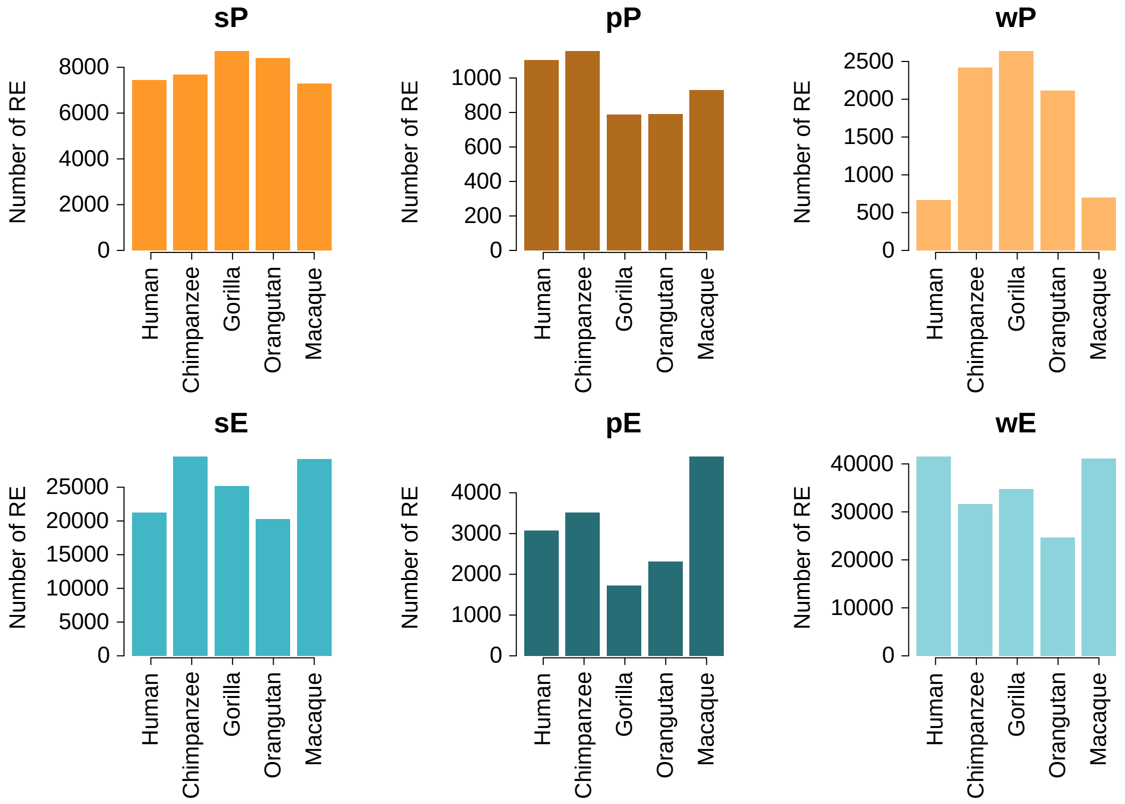


**Supplementary Figure 9. Number of regulatory elements with the same regulatory state between replicates in every species.** Regulatory states are color-coded as in Figure 2. sP: strong promoter state; pP: poised promoter state; wP; weak promoter state; sE: strong enhancer state; pE: poised enhancer state; wE: weak enhancer state.

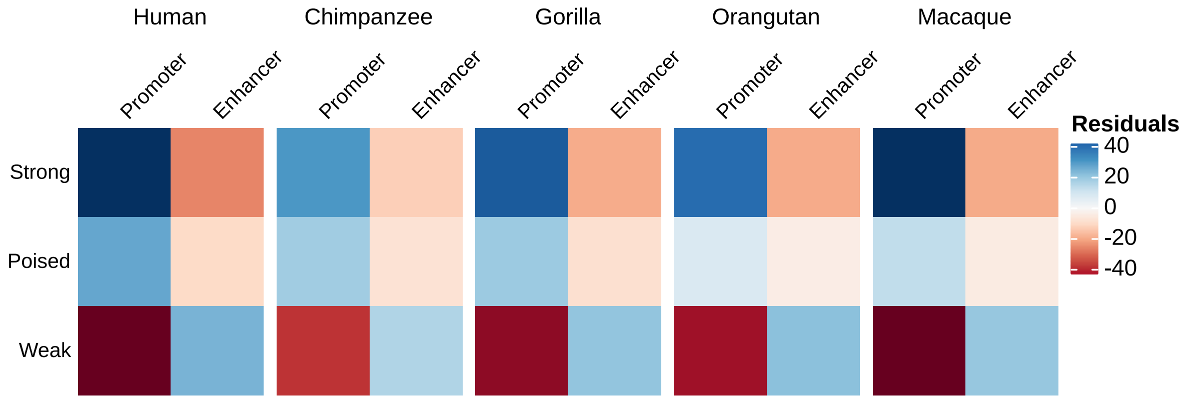


**Supplementary Figure 10. Strong and poised activities are significantly more associated with promoter regulatory states**. Heatmap shows the residual values from corresponding Chi-square tests.

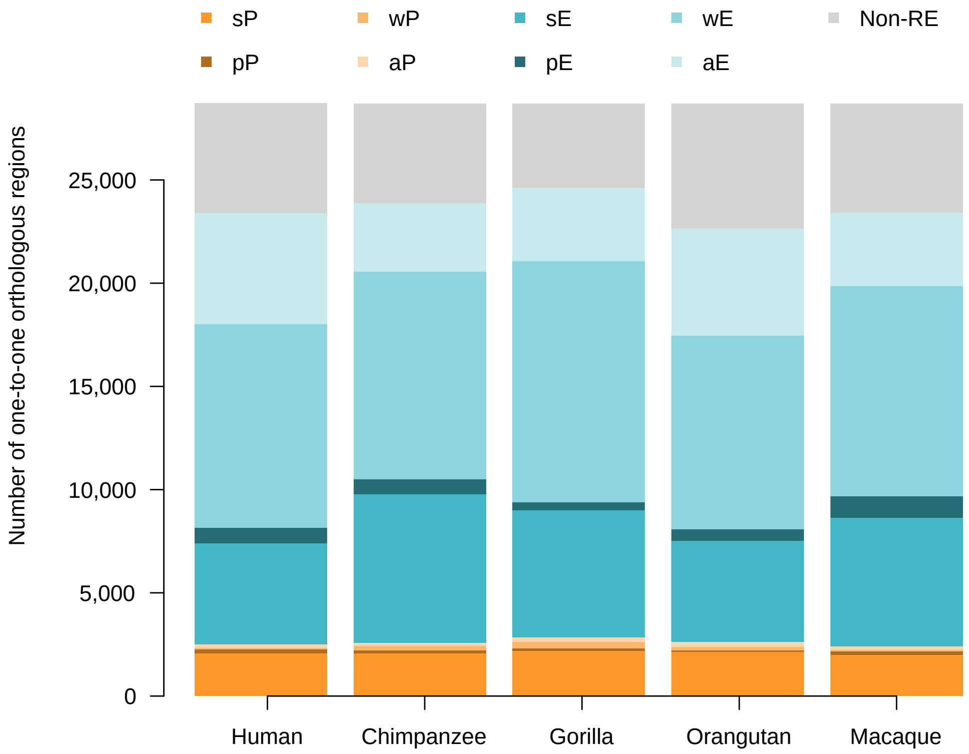


**Supplementary Figure 11. Regulatory states at orthologous regulatory regions.** Each bar shows for each species, the percentage of orthologous regulatory regions with the corresponding color-coded regulatory state.

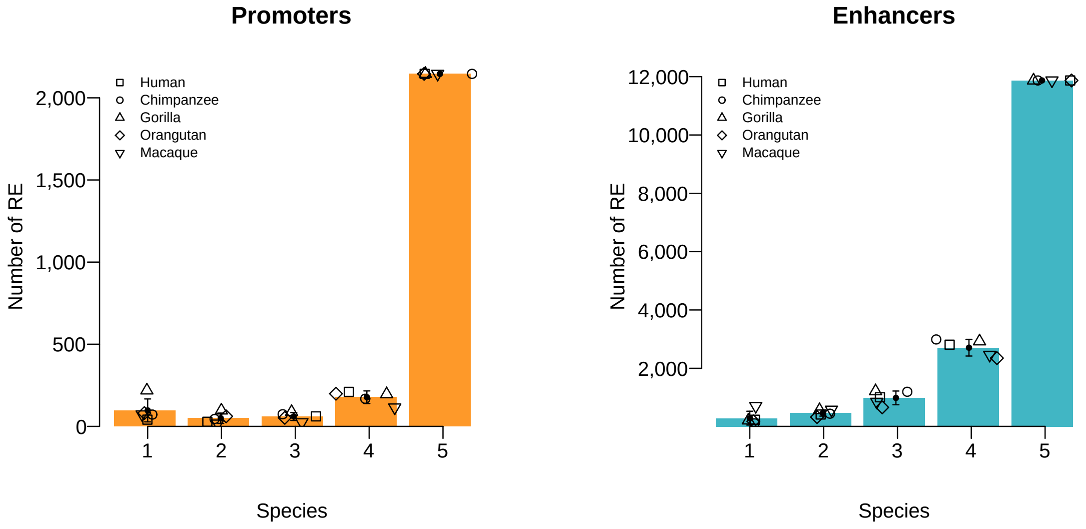


**Supplementary Figure 12. Promoter states at orthologous regulatory regions are more evolutionarily conserved than enhancer states.** Barplots show the average number of orthologous regulatory regions with a promoter or enhancer statee in 1, 2, 3, 4 or 5 species. Error bars show the standard deviation across species. Fisher's exact test, odds ratio = 1.48, $P < 2.2 \times 10^{-16}$.

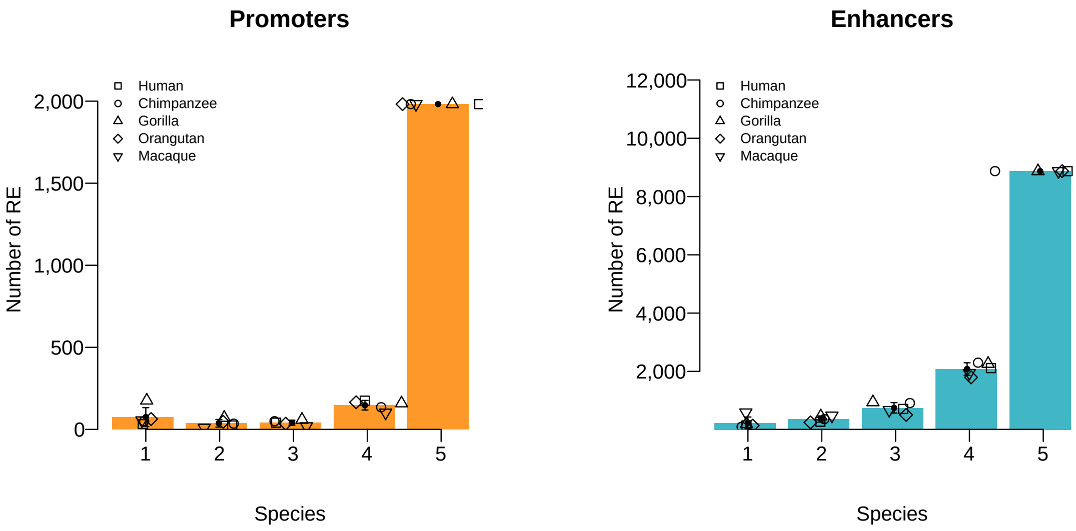


**Supplementary Figure 13. Promoter states at orthologous regulatory regions associated with human protein-coding genes are more evolutionarily conserved than enhancer states.** Figure caption as in Figure S12. Fisher exact test, odds ratio = 1.84, $P < 2.2 \times 10^{-16}$.

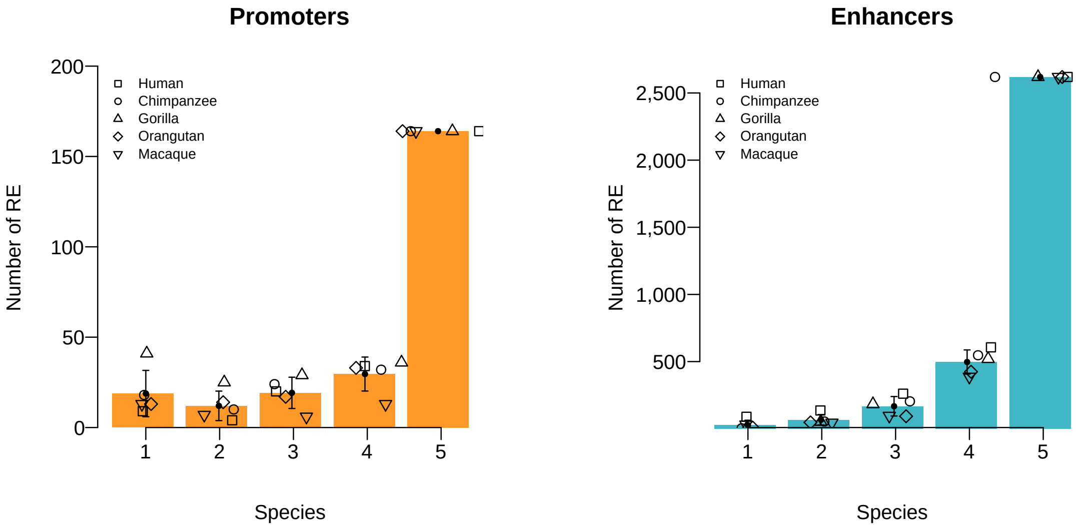


**Supplementary Figure 14. Promoter states at orthologous regulatory regions associated with human non-coding genes are less evolutionarily conserved than enhancer states.** Figure caption as in Figure S12. Fisher exact test, odds ratio = 0.39, $P < 2.2 \times 10^{-16}$.

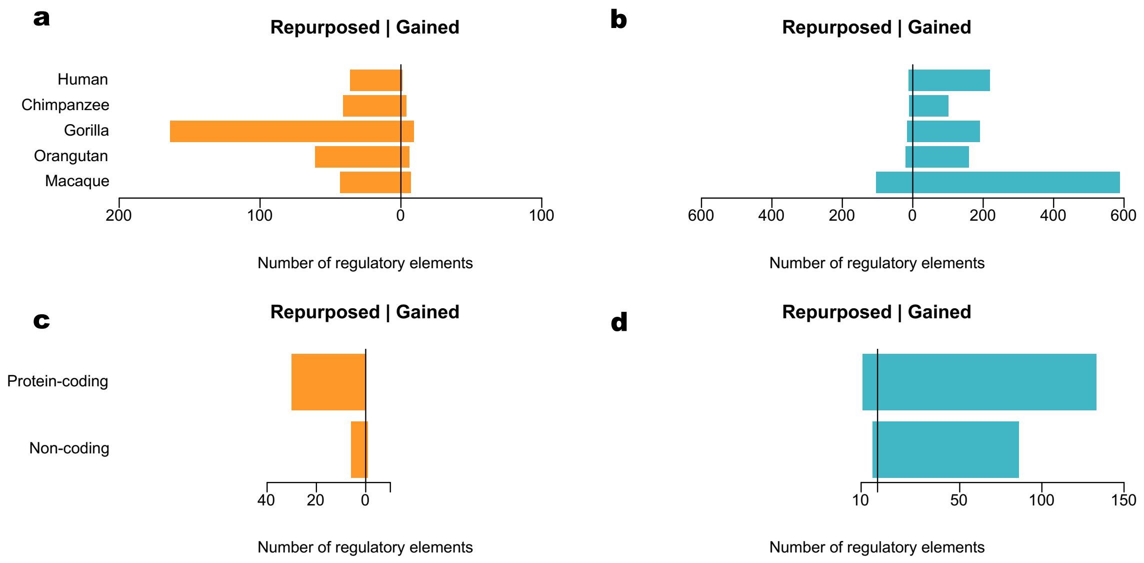


**Supplementary Figure 15. Promoters are gained through repurposing events where enhancers emerge from orthologous regions with no regulatory activity.** Barplots show, for promoters and enhancers, the number of regulatory elements gained through repurposing events (left side barplots) and acquired de novo at orthologous regions with no regulatory activity on the other species (right side barplots).

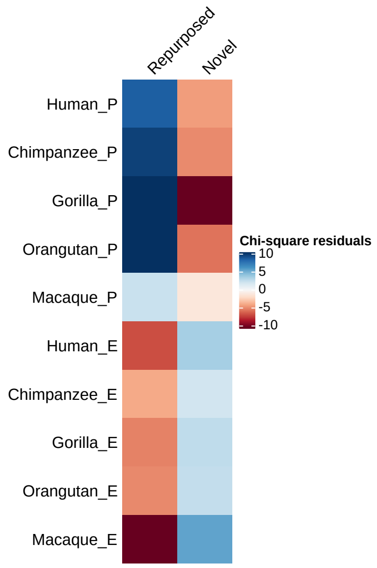


**Supplementary Figure 16. Promoters are gained through repurposing events where enhancers emerge from orthologous regions with no regulatory activity**. Cell values correspond to the residuals from a Chi-square test.

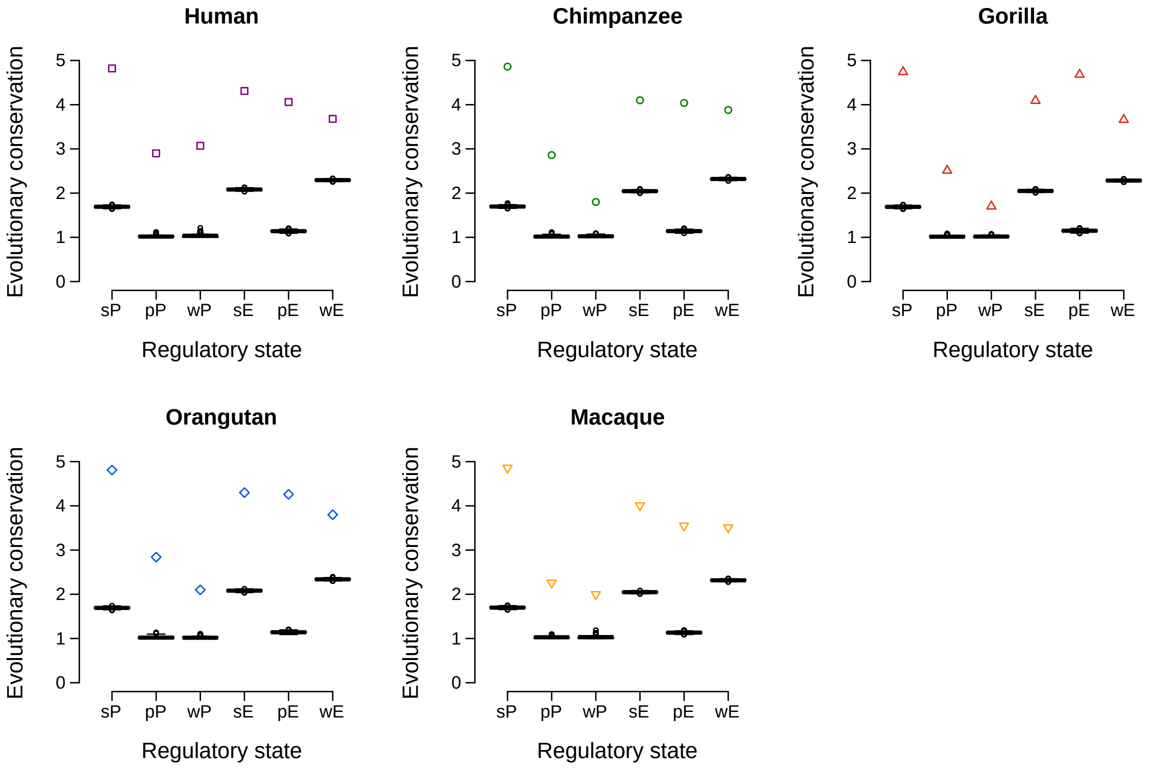


**Supplementary Figure 17. Randomization analyses show that conserved regulatory states are found more frequently than expected by chance.** Evolutionary conservation was defined as the average number of species in which a regulatory state was conserved. For each species and regulatory state, boxplots show the distributions obtained in 1,000 randomizations whereas the points indicate the average evolutionary conservation observed for each species and regulatory state.

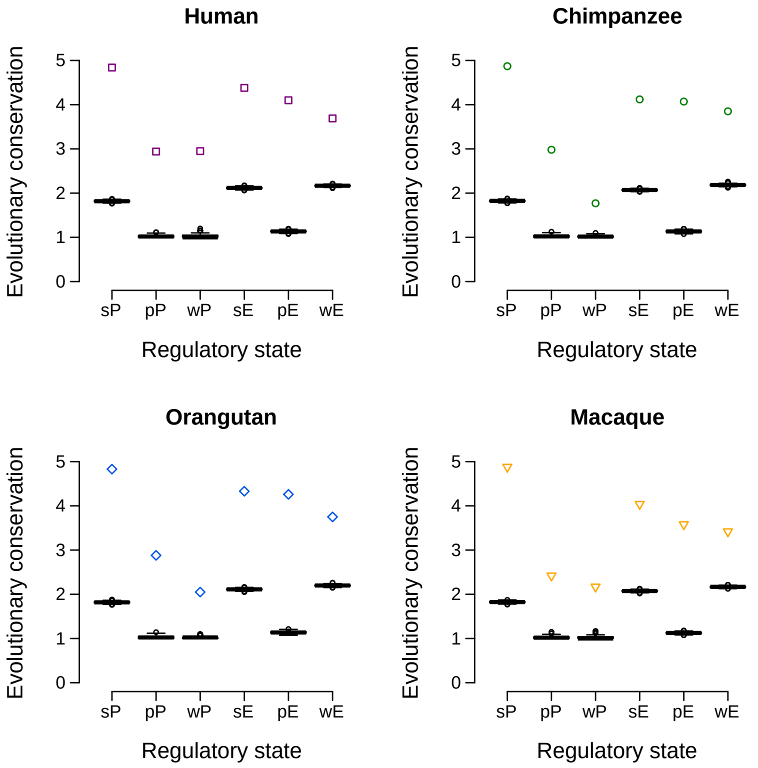


**Supplementary Figure 18. Randomization analyses show that conserved regulatory states are found more frequently than expected by chance in orthologous regulatory regions associated with human protein-coding genes.** Figure caption as in Supplementary Figure 17.

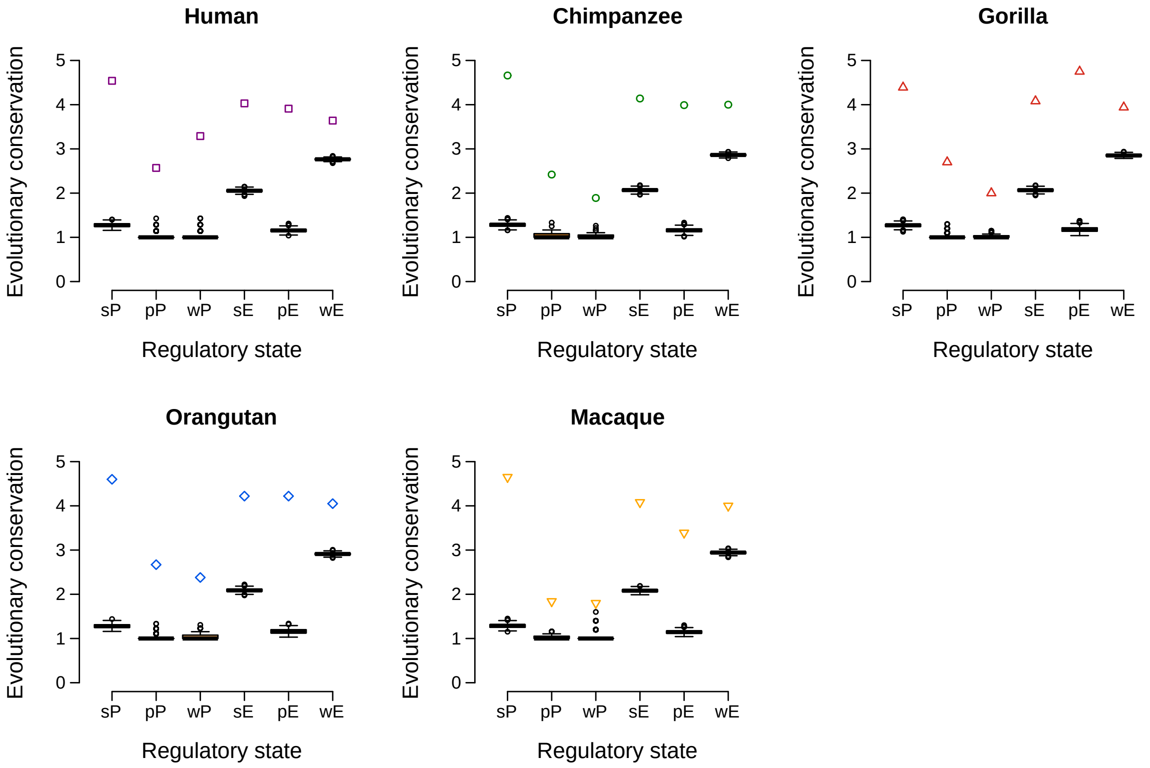


**Supplementary Figure 19. Randomization analyses show that conserved regulatory states are found more frequently than expected by chance in orthologous regulatory regions associated with human non-coding genes.** Figure caption as in Supplementary Figure 17.

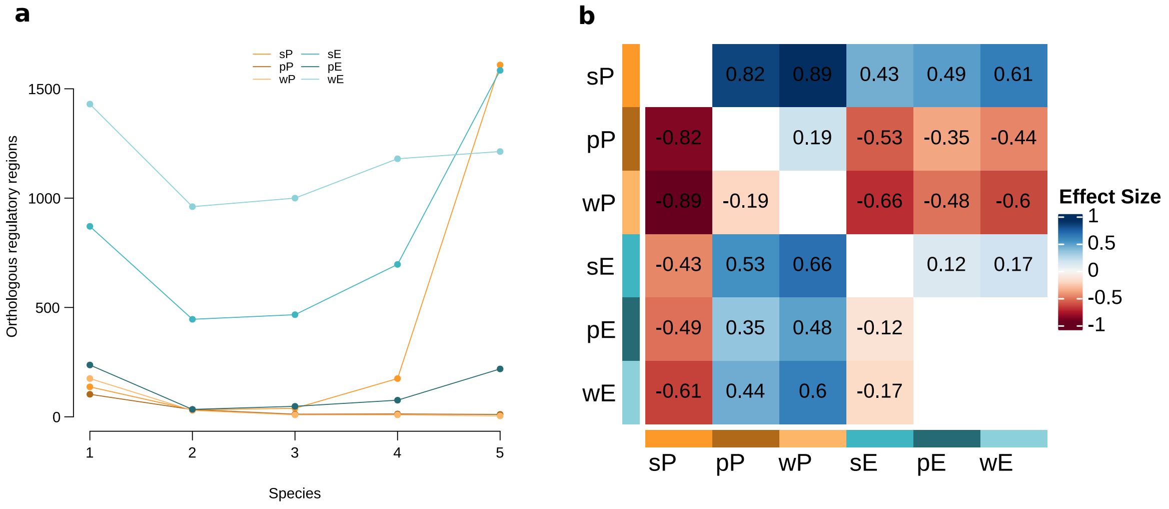


**Supplementary Figure 20. Different regulatory states show different evolutionary dynamics.** Analysis restricted to those orthologous regulatory regions associated with human protein-coding genes. Patterns of evolutionary conservation for the different regulatory states. Kruskal-Wallis test, $P < 2.2 \times 10^{-16}$. Effect sizes of the pairwise comparison of the evolutionary conservation patterns between any two regulatory states (Dwass-Steel-Critchlow-Fligner test). Only the effect sizes for significantly different comparisons are shown ($P < 0.05$).

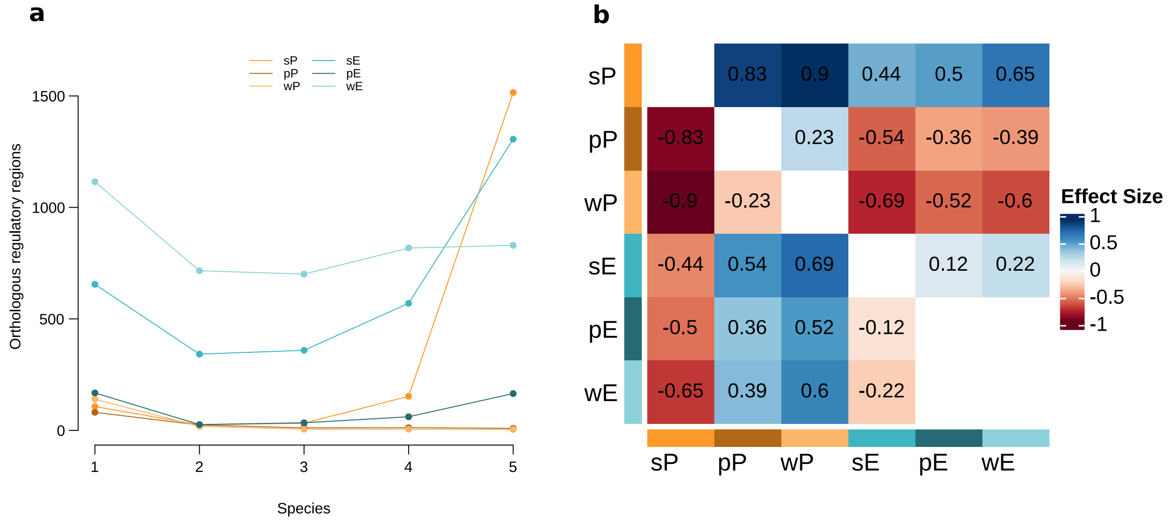


**Supplementary Figure 21. Different regulatory states show different evolutionary dynamics.** Analysis restricted to those orthologous regulatory regions associated with human non-coding genes. Patterns of evolutionary conservation for the different regulatory states. Kruskal-Wallis test, $P < 2.2 \times 10^{-16}$. Effect sizes of the pairwise comparison of the evolutionary conservation patterns between any two regulatory states (Dwass-Steel-Critchlow-Fligner test). Only the effect sizes for significantly different comparisons are shown ($P < 0.05$).

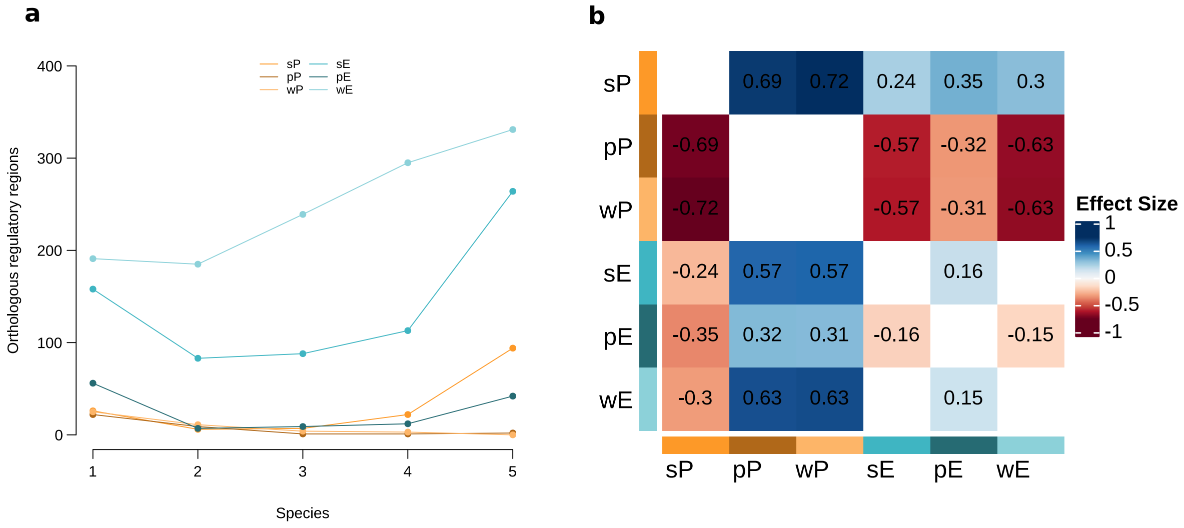


**Supplementary Figure 22. Different regulatory states show different evolutionary dynamics.** Analysis restricted to those orthologous regulatory regions associated with human non-coding genes. Patterns of evolutionary conservation for the different regulatory states. Kruskal-Wallis test, $P < 2.2 \times 10^{-16}$. Effect sizes of the pairwise comparison of the evolutionary conservation patterns between any two regulatory states (Dwass-Steel-Critchlow-Fligner test). Only the effect sizes for significantly different comparisons are shown ($P < 0.05$).

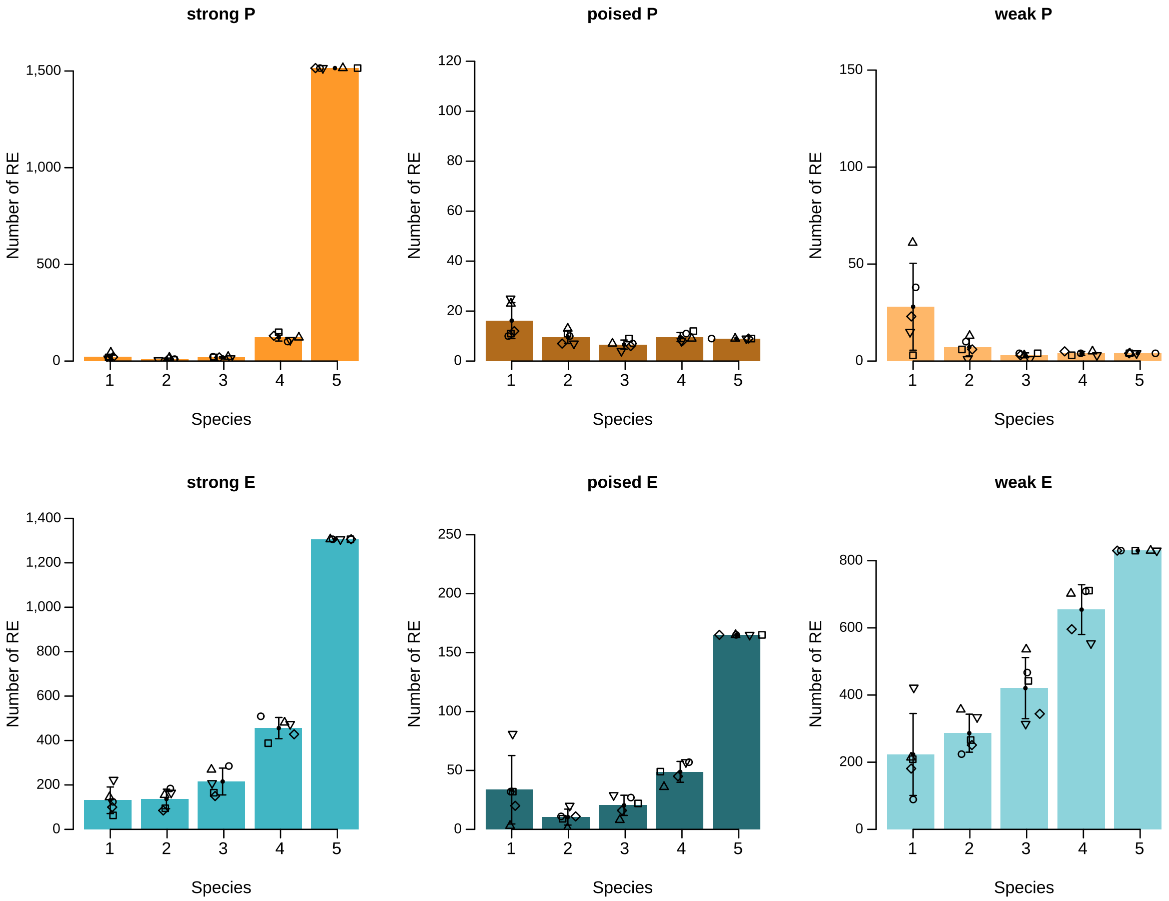


**Supplementary Figure 23. Patters of evolutionary conservation of each regulatory state at orthologous regions associated with human protein-coding genes.** Barplots show the average number of orthologous regulatory regions across species with the corresponding color-coded epigenetic state conserved in 1, 2, 3, 4 or 5 species. Error bars show the standard deviation across species and differently shaped dots show the number of regulatory regions with this conservation for each species.

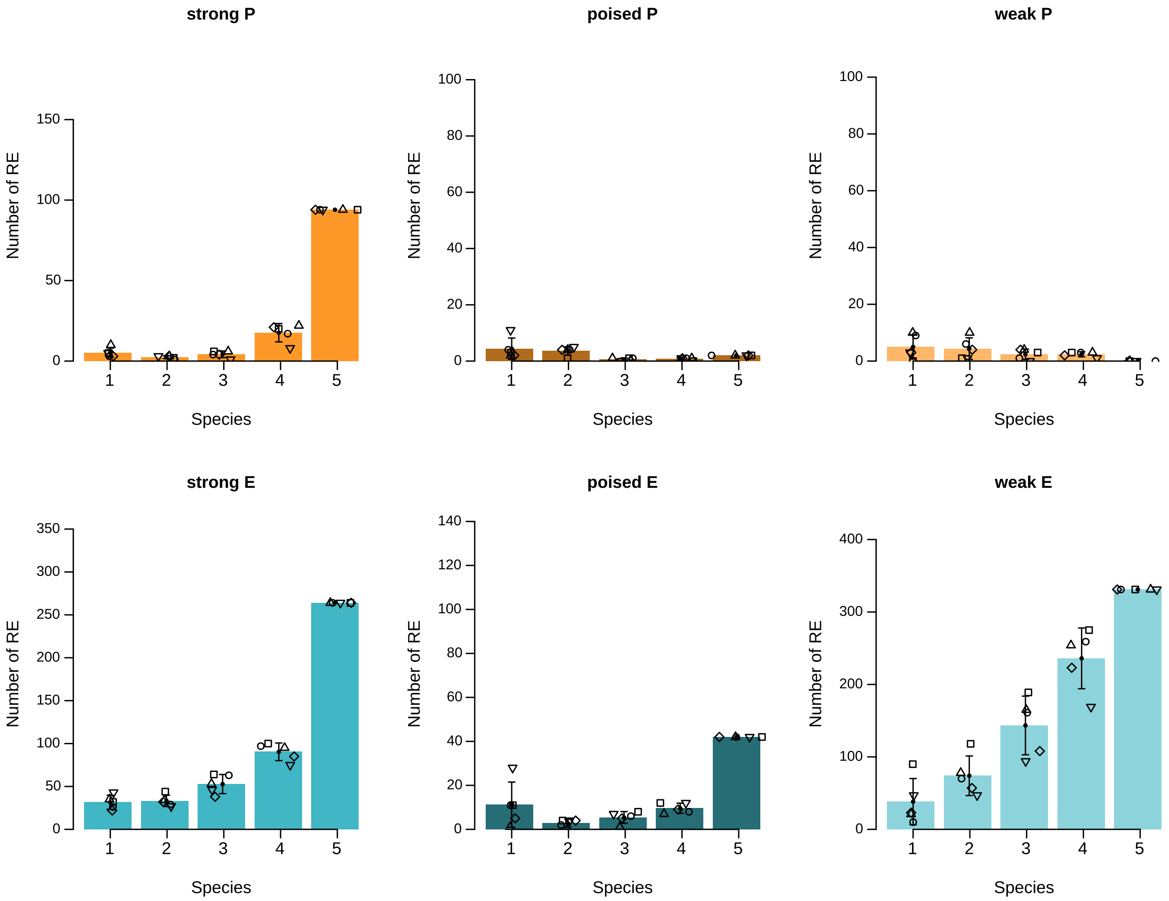


**Supplementary Figure 24. Patters of evolutionary conservation of each regulatory state at orthologous regions associated with human non-coding genes.** Barplots show the average number of orthologous regulatory regions across species with the corresponding color-coded epigenetic state conserved in 1, 2, 3, 4 or 5 species. Error bars show the standard deviation across species and differently shaped dots show the number of regulatory regions with this conservation for each species.

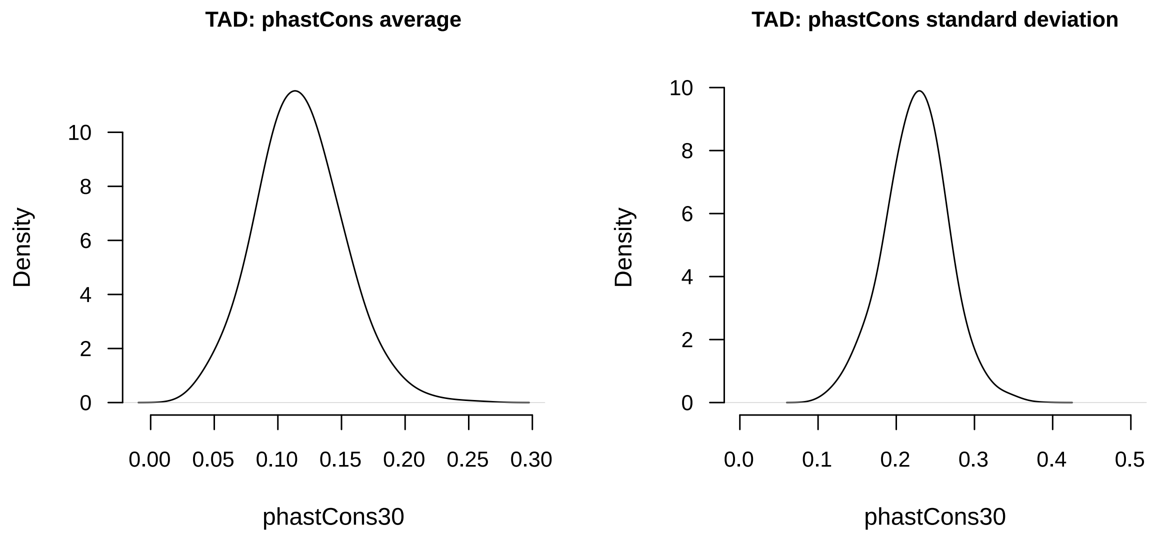


**Supplementary Figure 25. Distribution of the average and standard deviation phatsCons30 scores associated with TADs defined in cell the line GM12878.**

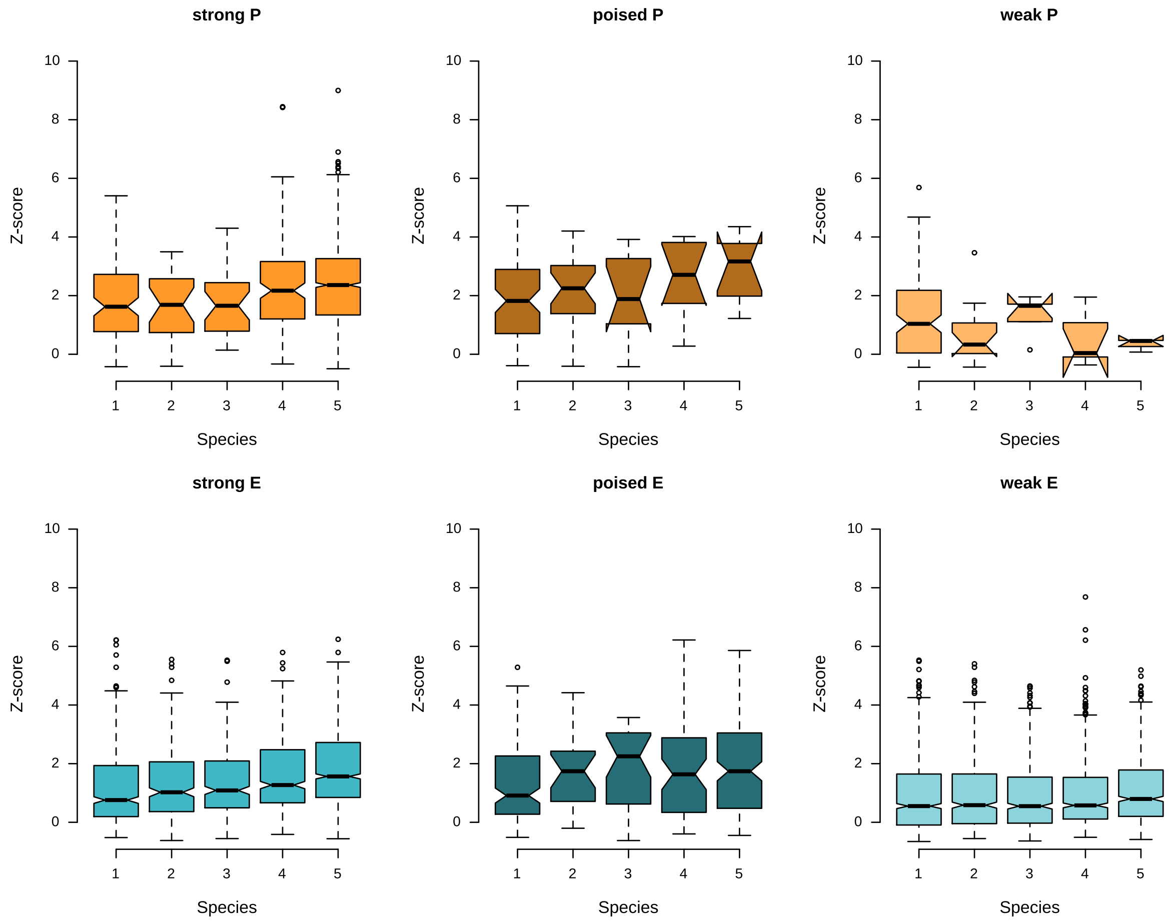


**Supplementary Figure 26. Distribution of sequence conservation scores of orthologous regulatory regions associated with human protein-coding genes in 1, 2, 3, 4 or 5 species.** Different regulatory states are color-coded.

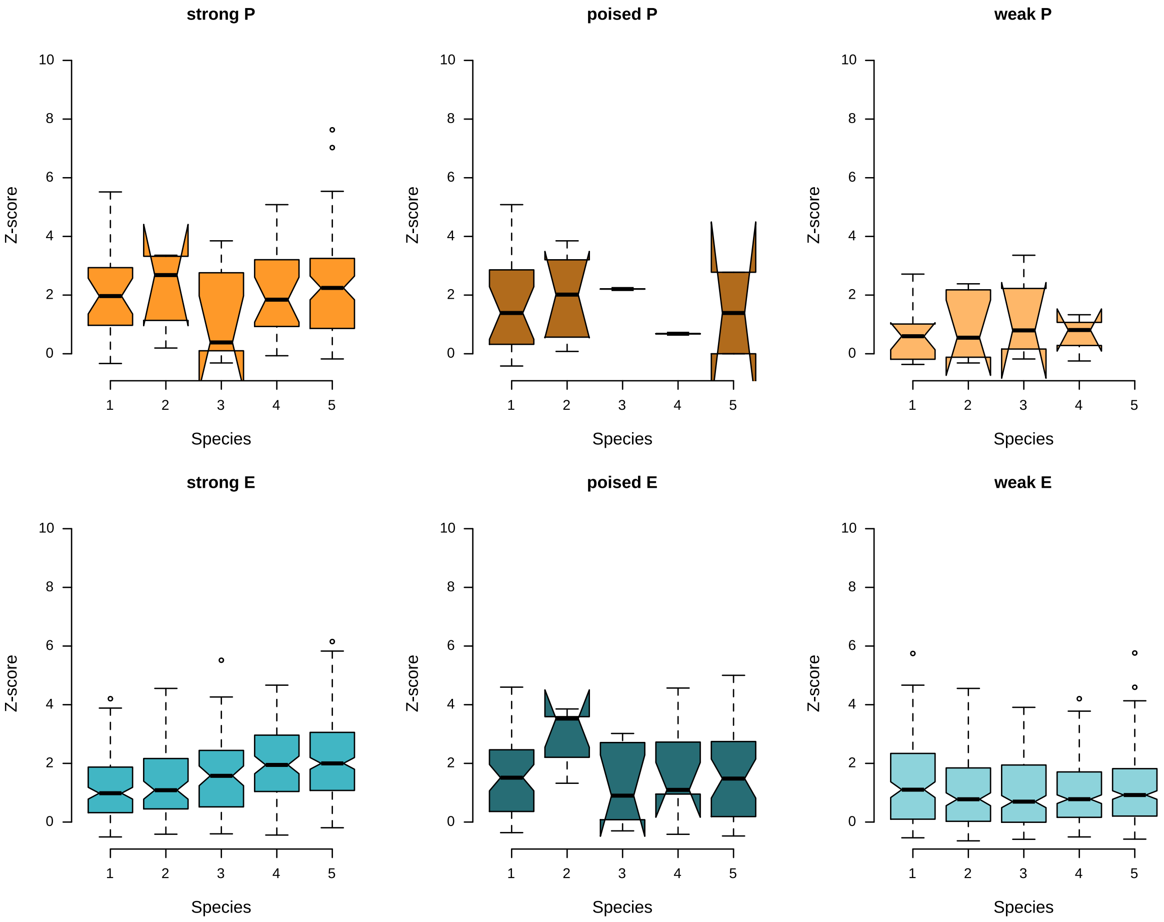


**Supplementary Figure 27. Distribution of sequence conservation scores of orthologous regulatory regions associated with human non-coding genes in 1, 2, 3, 4 or 5 species.** Different regulatory states are color-coded.

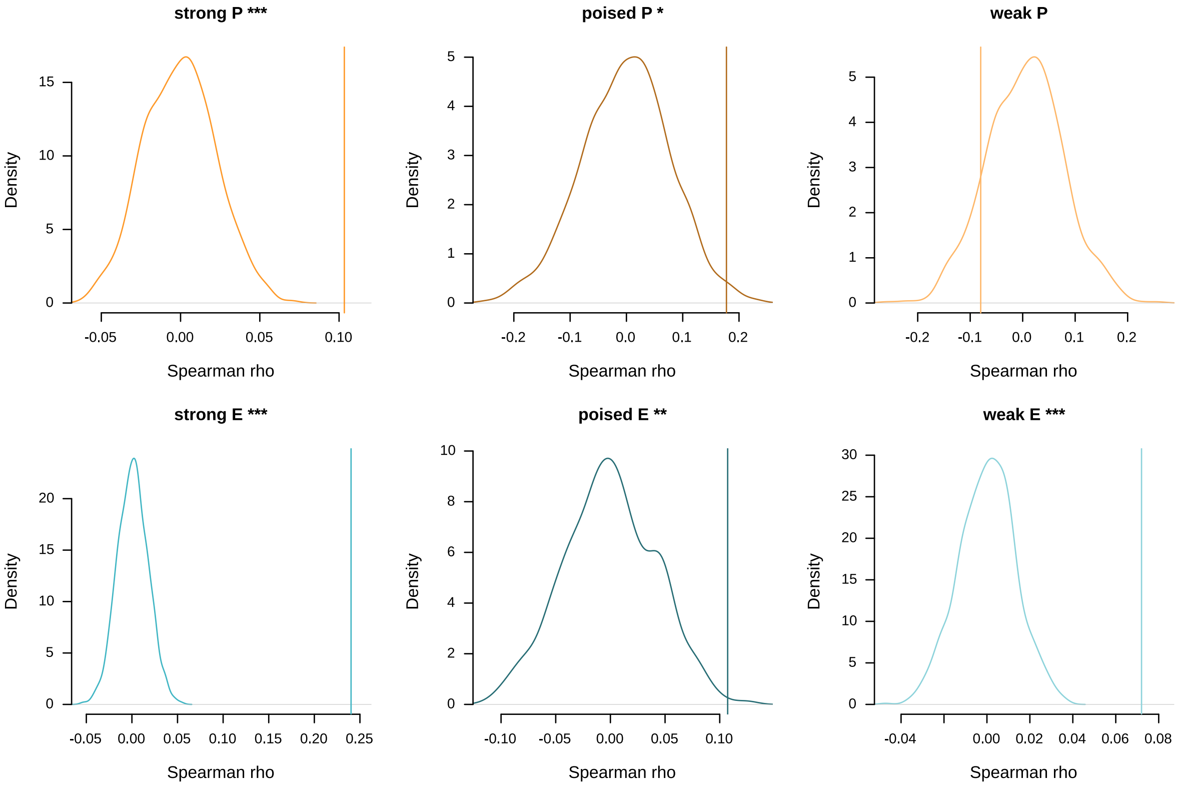


**Supplementary Figure 28. The evolutionary conservation of the regulatory state at orthologous regulatory regions is significantly positively correlated with the evolutionary conservation of the underlying sequence.** Density plots show the distribution of the Spearman rho correlation values obtained in 1,000 randomizations. Vertical lines indicate the Spearman rank correlation $\rho$ value of the real data. Asterisk represent associated P-values (* P-value < 0.05; ** P-value < 0.01; *** P-value < 0.001).

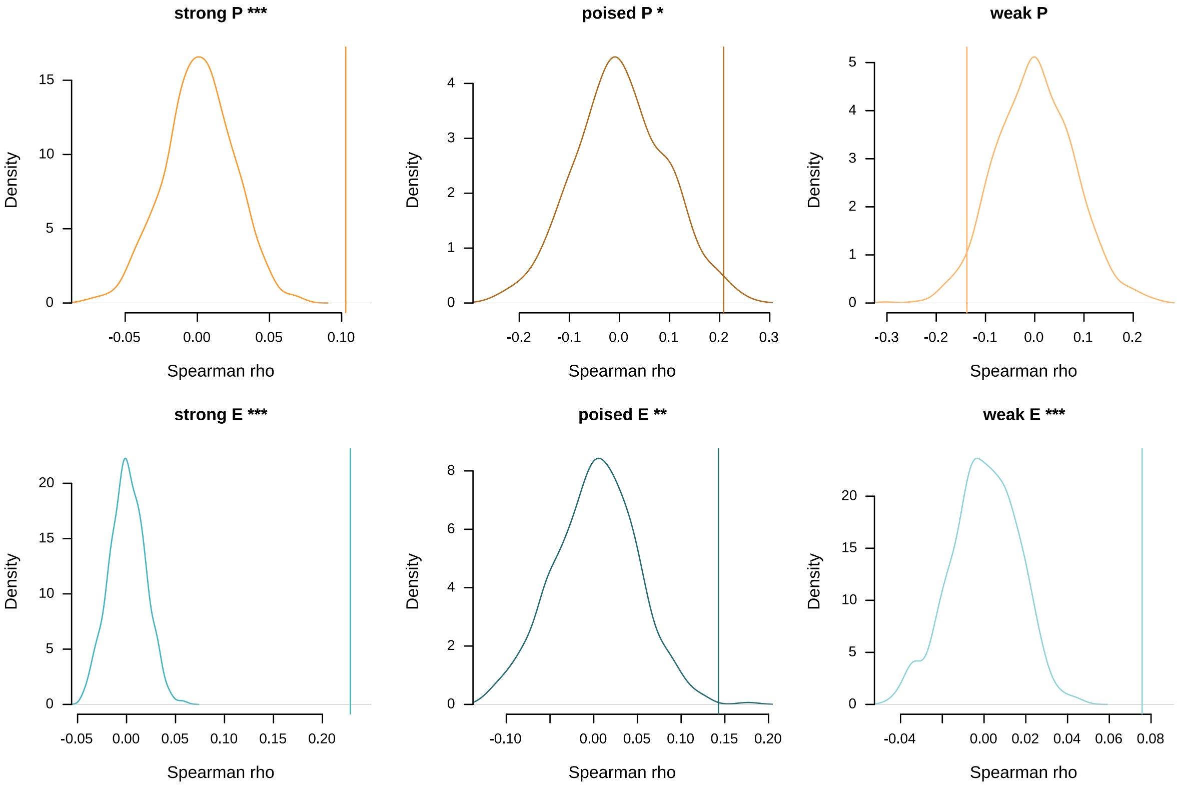


**Supplementary Figure 29. The evolutionary conservation of the regulatory state at orthologous regulatory regions associated with human protein-coding is significantly positively correlated with the evolutionary conservation of the underlying sequence.** Density plots show the distribution of the Spearman rank correlation ρ values obtained in 1,000 randomizations. Vertical lines indicate the Spearman rank correlation ρ value of the real data. Asterisk represent associated P-values (* P < 0.05; ** P < 0.01; *** P < 0.001).

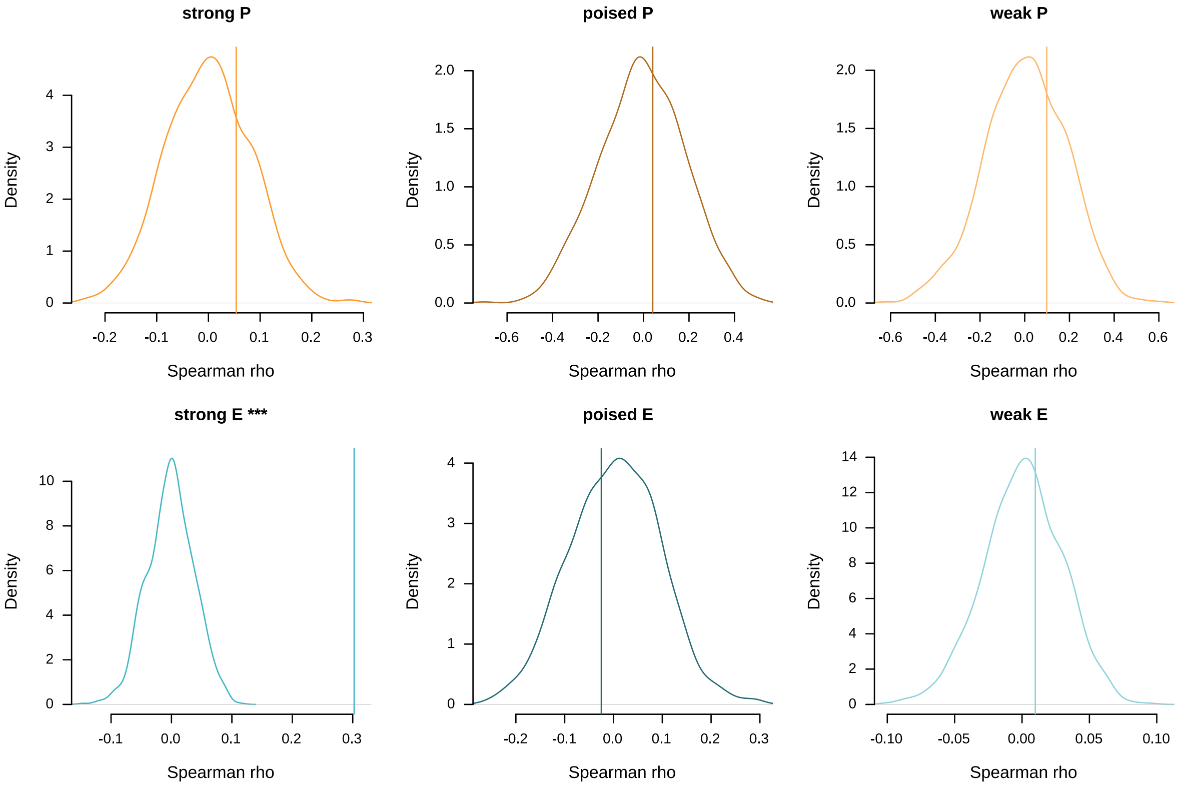


**Supplementary Figure 30. The evolutionary conservation of the regulatory state at orthologous regulatory regions associated with human non-coding genes is only significantly positively correlated with the evolutionary conservation of the underlying sequence for strong enhancer sates**. Density plots show the distribution of the Spearman rank correlation ρ values obtained in 1,000 randomizations. Vertical lines indicate the Spearman rank correlation *ρ* value of the real data. Asterisk represent associated P-values (* P-value < 0.05; ** P-value < 0.01; *** P-value < 0.001).

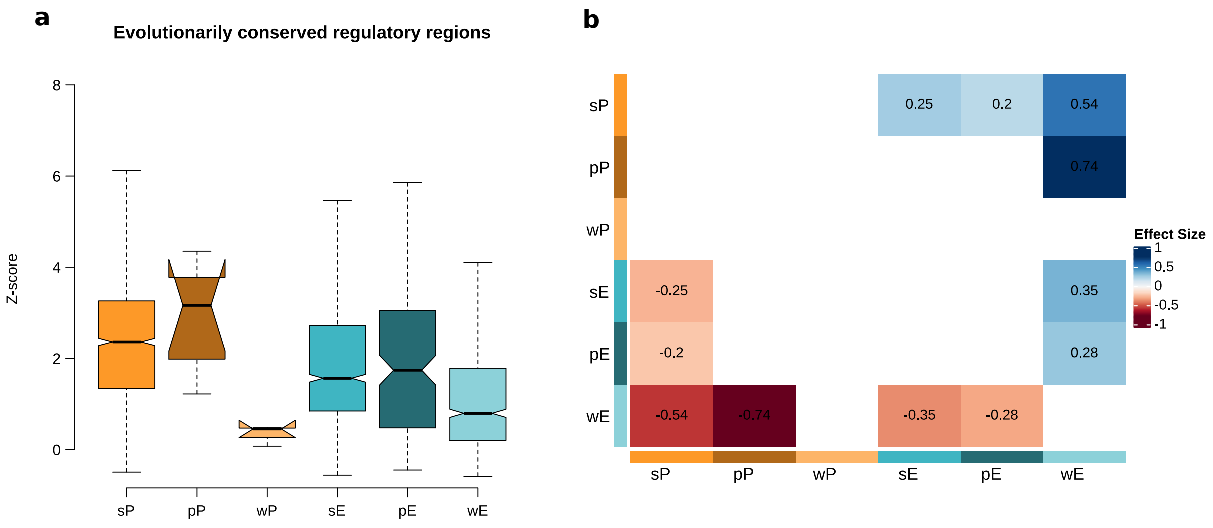


**Supplementary Figure 31. Fully conserved strong promoters have the highest evolutionarily conserved sequences. a,** Distribution of z-score values of evolutionarily conserved regulatory regions associated with human protein-coding genes (Kruskal-Wallis test, $P < 2.2 \times 10^{-16}$). **b,** Effect sizes correspond to significant pairwise comparisons from (a) (Dwass-Steel-Critchlow-Fligner test, $P < 0.05$).

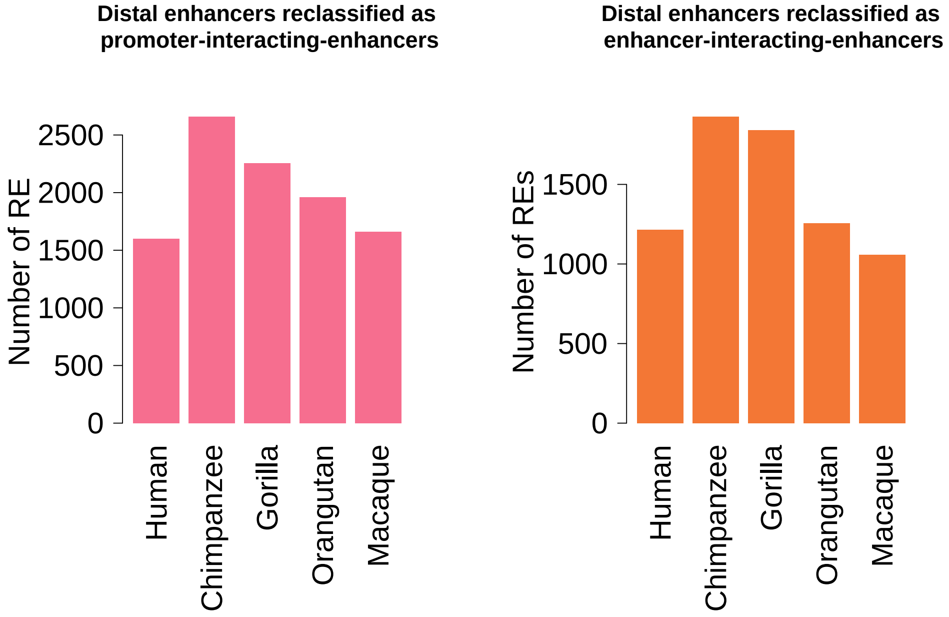


**Supplementary Figure 32. Number of distal enhancers annotated as PiE and EiE per species through the integration of 3D chromatin data.** Promoter-interacting-enhancers are gene-associated enhancers that interact with genic promoters. Enhancer-interacting-enhancers are gene-associated enhancers that interact between them.

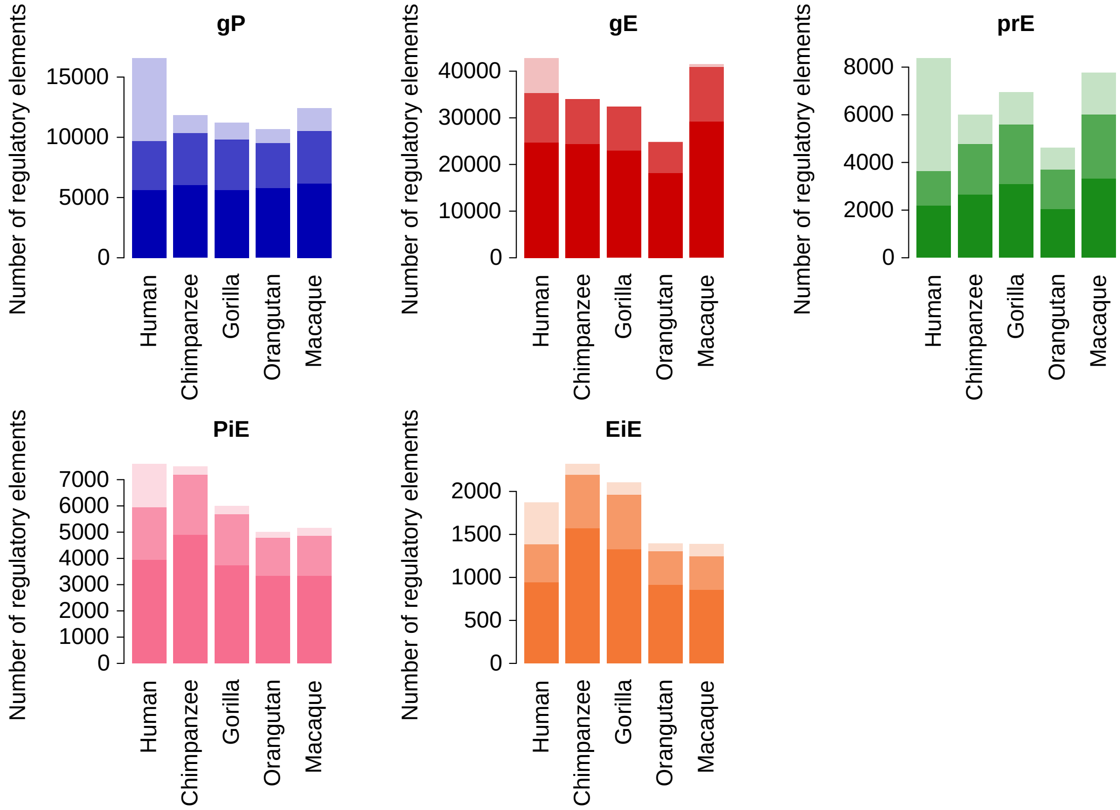


**Supplementary Figure 33. Number of regulatory elements annotated as genic promoter (gP), intragenic enhancers (gE), proximal enhancers (prE), promoter-interactingenhancers (PiE) and enhancer-interacting enhancers (EiE).** Dark, medium and light shades indicate the number of regulatory elements associated with 1-to-1 orthologous protein-coding genes, protein-coding genes or non-coding genes, respectively

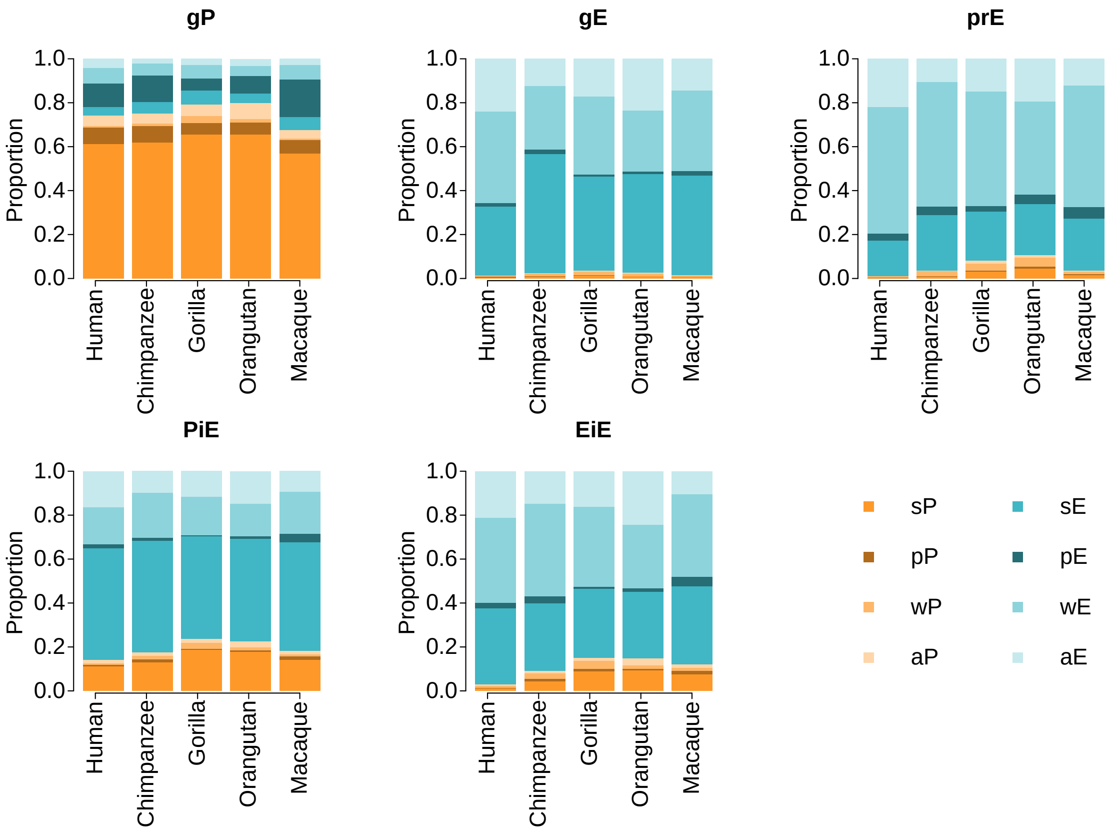


**Supplementary Figure 34. Correspondence between regulatory state and component of regulatory elements associated with orthologous protein-coding genes.** Each bar shows, for regulatory elements assigned a given regulatory component, the proportion of regulatory elements with the color-coded regulatory state. P: promoter; E: enhancer; s: strong; p: poised; w: weak; a: ambiguous (different activity states between biological replicates).

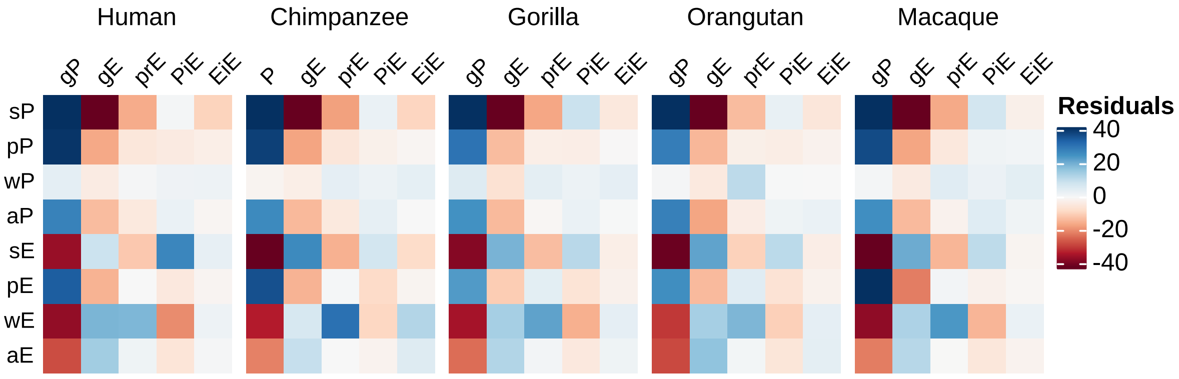


**Supplementary Figure 35. Genic promoters are significantly more associated with promoter regulatory states, while gene-associated enhancers are significantly more associated with enhancer regulatory states.** Heatmaps show the residuals values of a Chisquare test run separately for each species ($P < 2.2 \times 10^{-16}$ in all species).

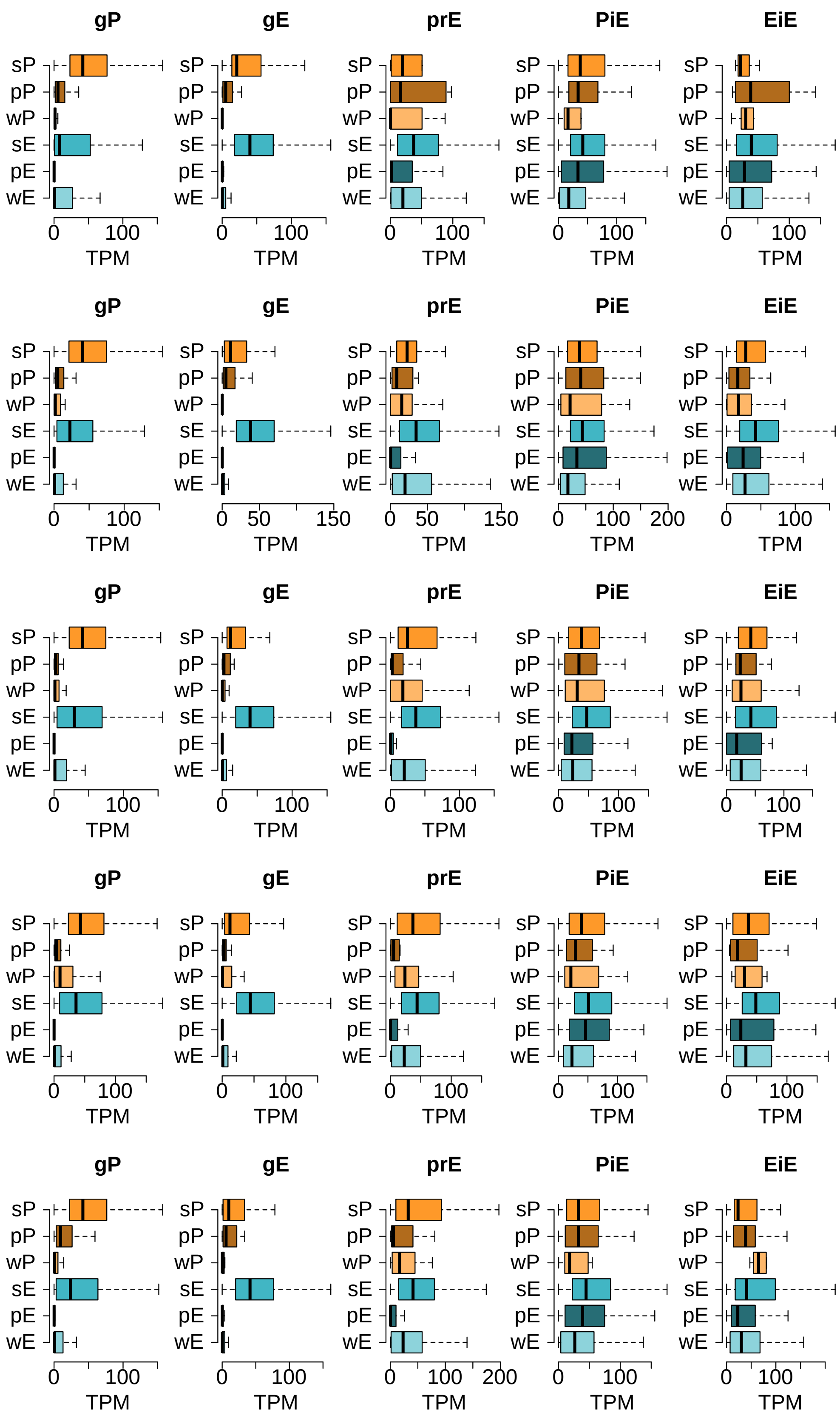


**Supplementary Figure 36. Gene expression levels are associated with the regulatory state and the type of regulatory component of the associated regulatory elements.** Boxplots show, for every species and regulatory role, the distribution of the TPM expression levels of the genes they regulate. **a to e,** Distributions observed for human, chimpanzee, gorilla, orangutan and macaque.

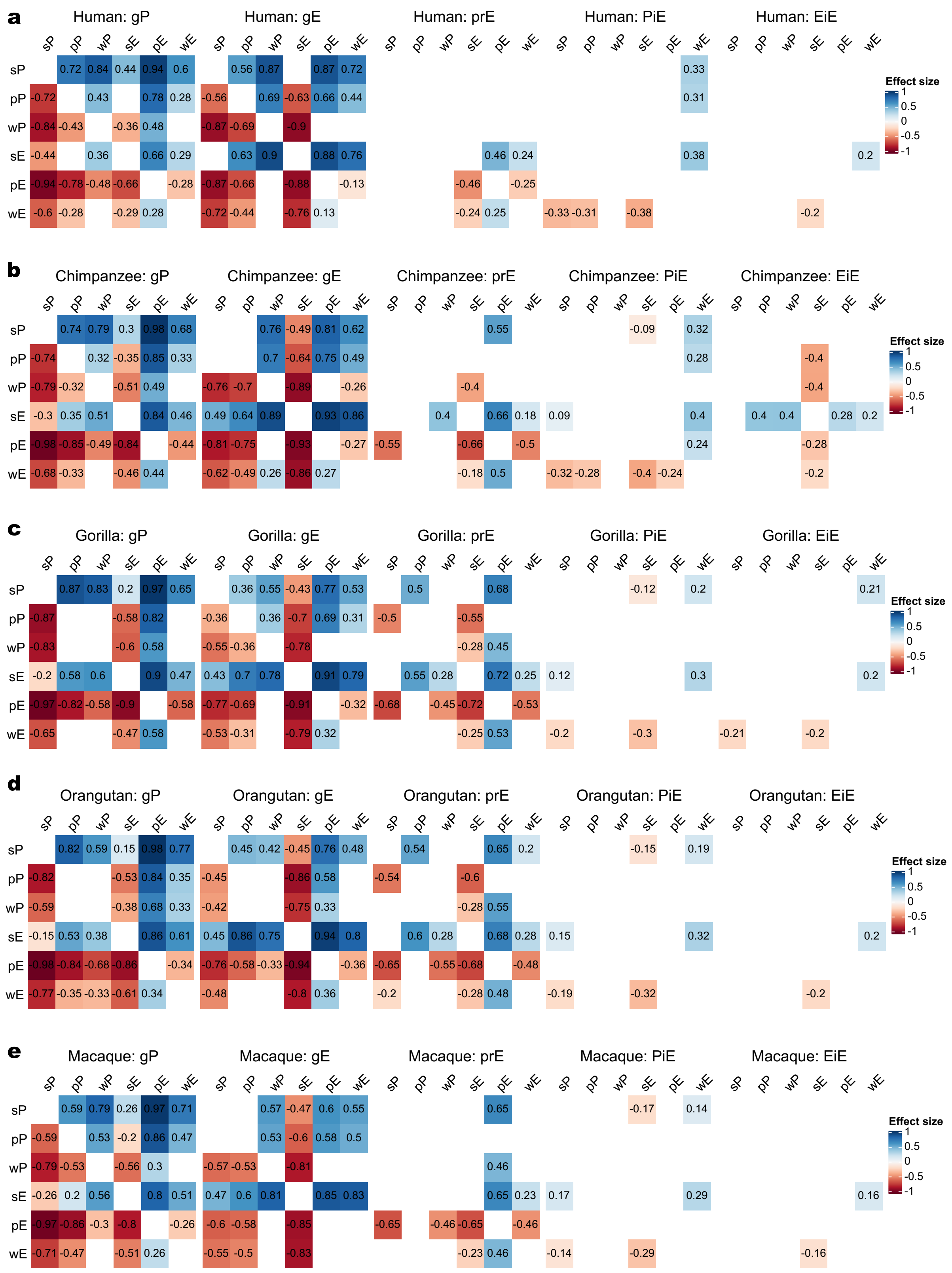

**Supplementary Figure 37. The expression levels of genes associated with regulatory elements with strong activities are significantly higher than those of genes associated with regulatory elements with poised activities, which showed significantly lower expression levels.** For each species and regulatory component, we compared the expression levels of the associated genes based on the presence of regulatory elements with strong, poised or weak regulatory promoter or enhancer states (Kruskal-Wallis, $P < 0.05$ in all combinations) (Methods). Cell values show the effect size for those significantly different pairwise comparisons (Dwass-Steel-Critchlow-Fligner test. $P < 0.05$). **a to e,** Values for human, chimpanzee, gorilla, orangutan and macaque.

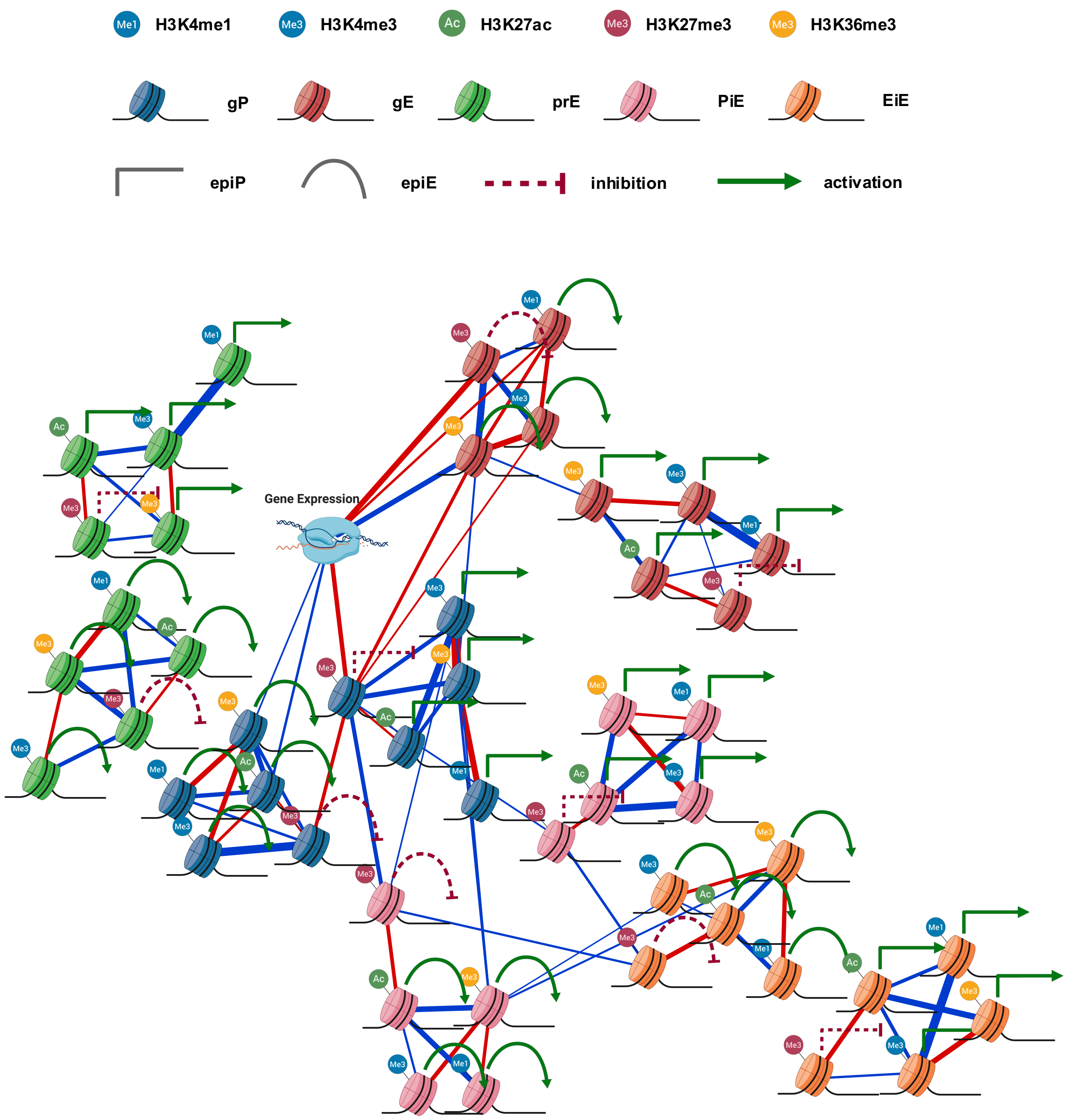


**Supplementary Figure 38. Global residuals Sparse Partial Correlation Network.** Blue edges represent positive partial correlations and red edges negative ones. Edge widths are proportional to absolute partial correlation values within each network. Only nodes for values with significant and relevant partial correlations (see Methods) were represented (minimal partial correlation = -0.323; maximal partial correlation = 0.354; all partial correlations Benjamini-Hochberg's Q-value < $3.4 \times 10^{-112}$).

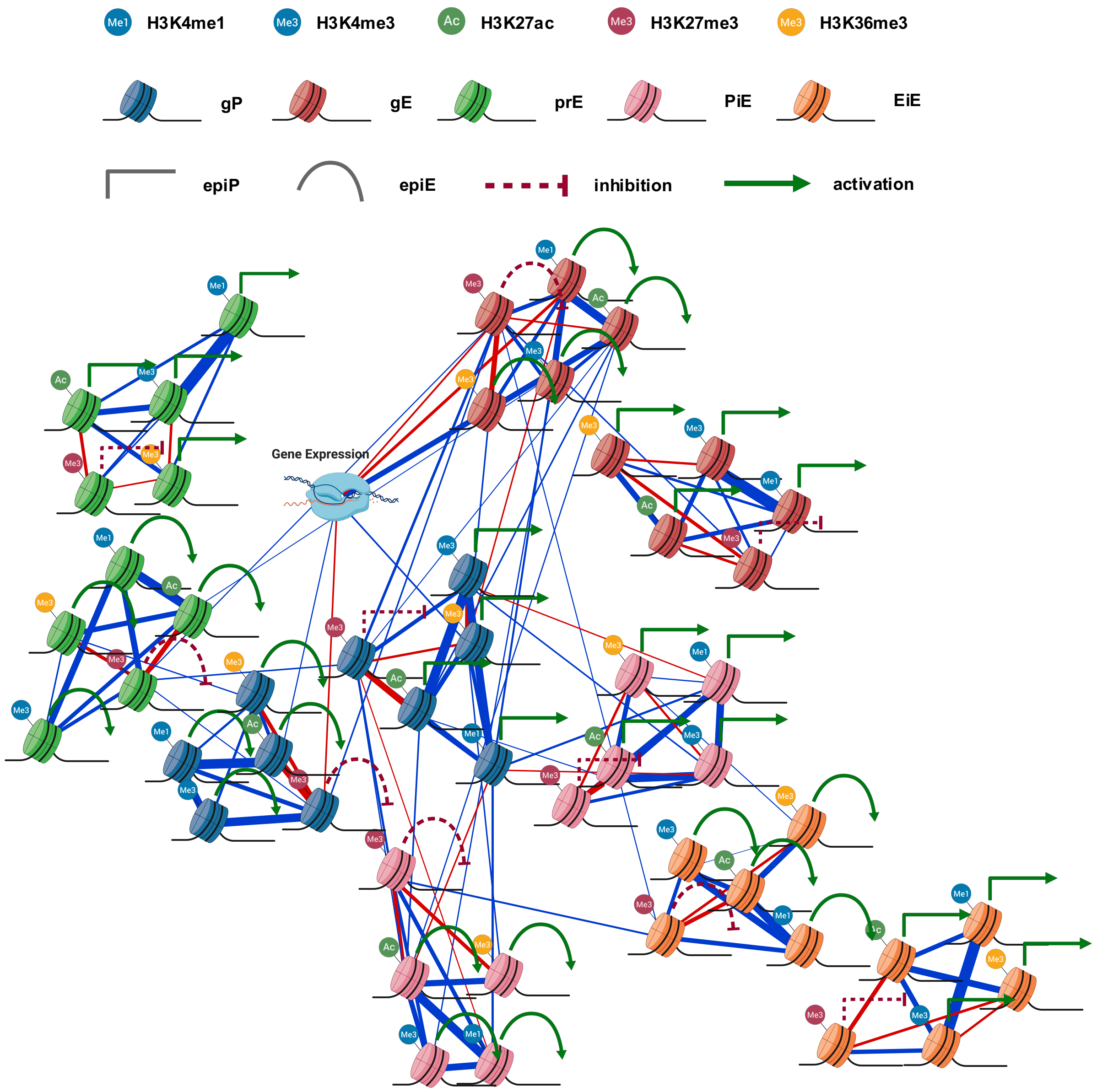


**Supplementary Figure 39. Global Sparse Partial Correlation Network**. Blue edges represent positive partial correlations and red edges negative ones. Edge widths are proportional to absolute partial correlation values within each network. Only nodes for values with significant and relevant partial correlations (see Methods) were represented (minimal partial correlation = -0.499; maximal partial correlation = 0.788; all partial correlations Benjamini-Hochberg's Q-value < $7.4 \times 10^{-22}$).

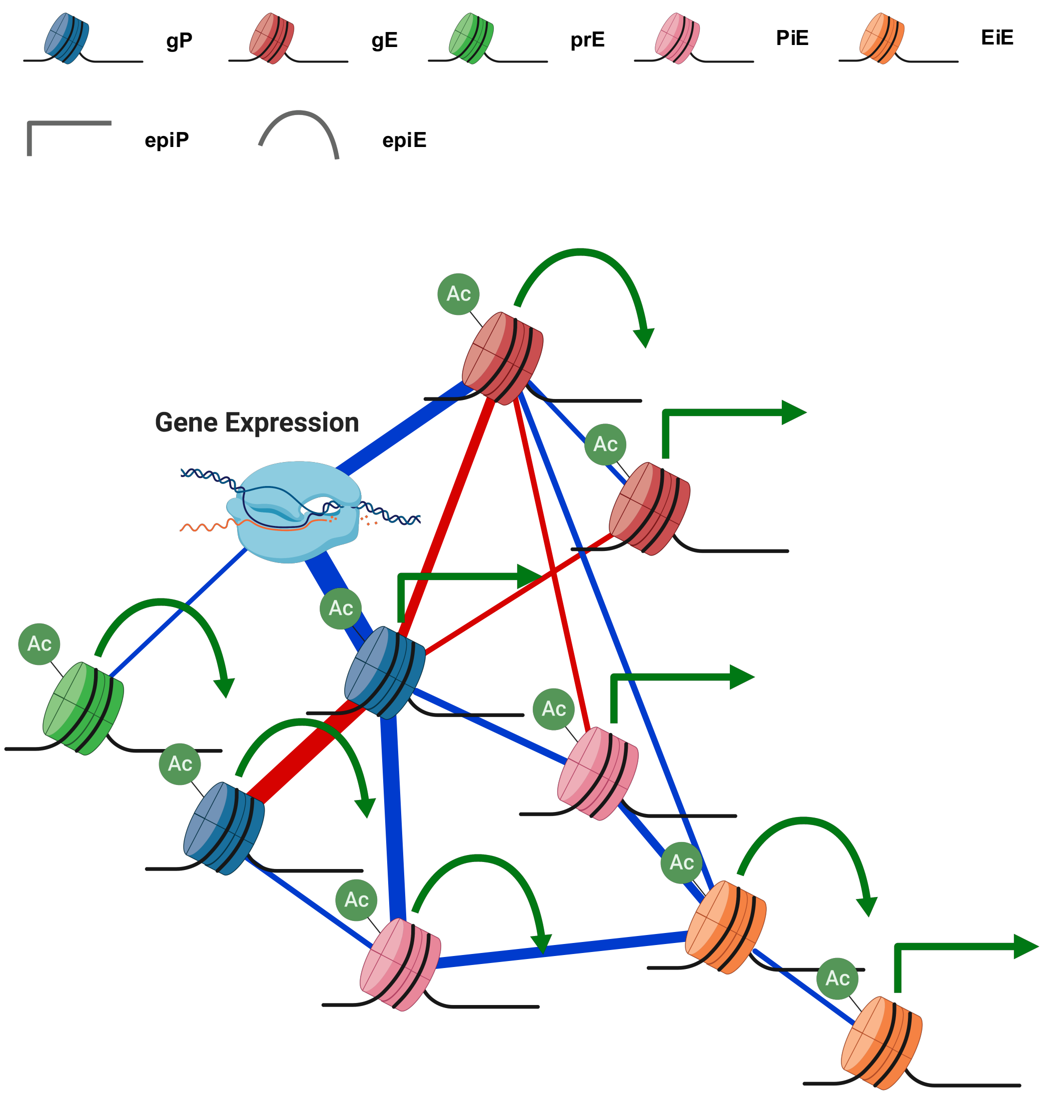


**Supplementary Figure 40. H3K27ac Sparse Partial Correlation Network**. Blue edges represent positive partial correlations and red edges negative ones. Edge widths are proportional to absolute partial correlation values within each network. Only nodes for values with significant and relevant partial correlations (see Methods) were represented (minimal partial correlation = -0.4; maximal partial correlation = 0.49; all partial correlations Benjamini-Hochberg's Q-value < $1.4 \times 10^{-109}$).

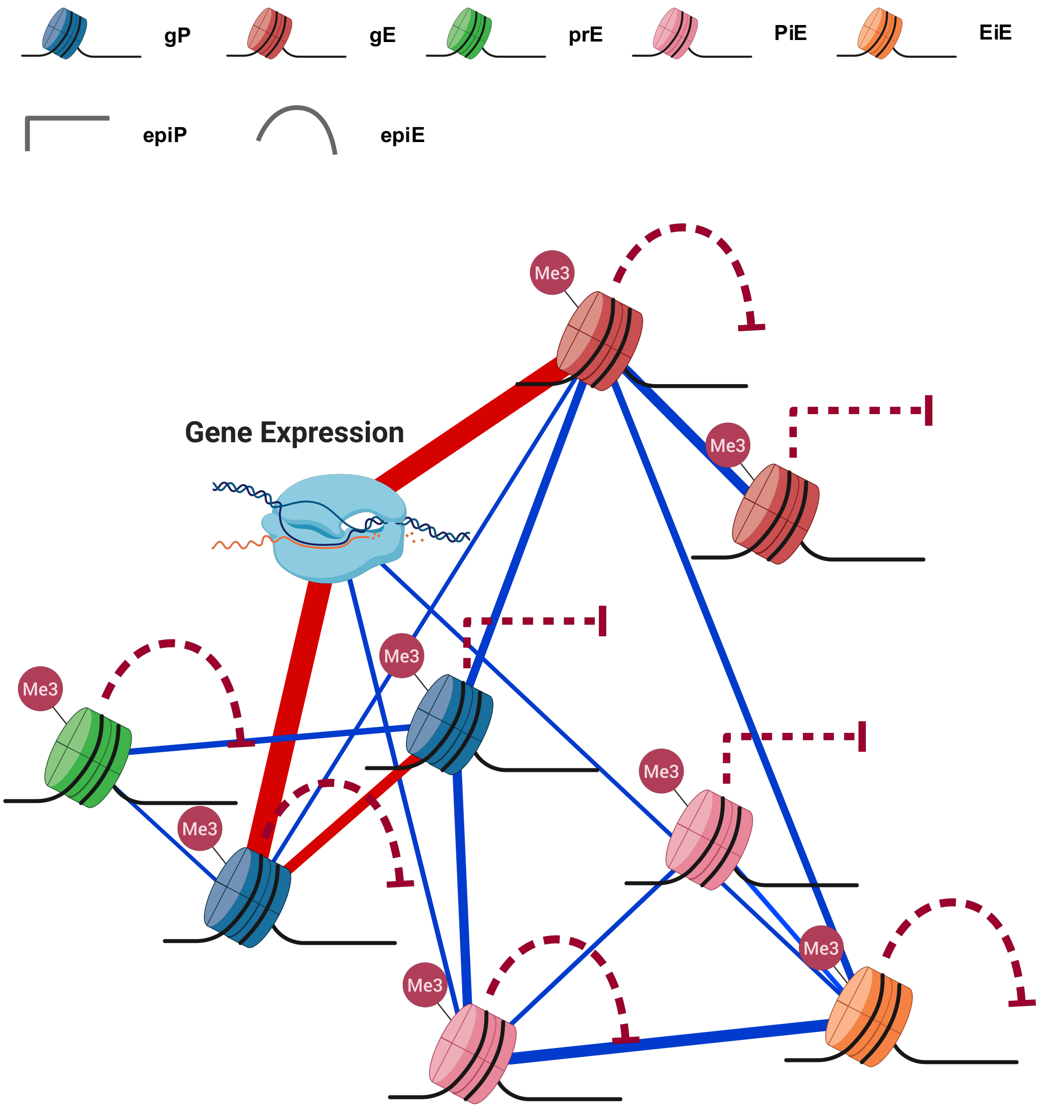


**Supplementary Figure 41. H3K27me3 Sparse Partial Correlation Network**. Blue edges represent positive partial correlations and red edges negative ones. Edge widths are proportional to absolute partial correlation values within each network. Only nodes for values with significant and relevant partial correlations (see Methods) were represented (minimal partial correlation = -0.4; maximal partial correlation = 0.2; all partial correlations Benjamini-Hochberg's Q-value < 3.9 x $10^{-57}$).

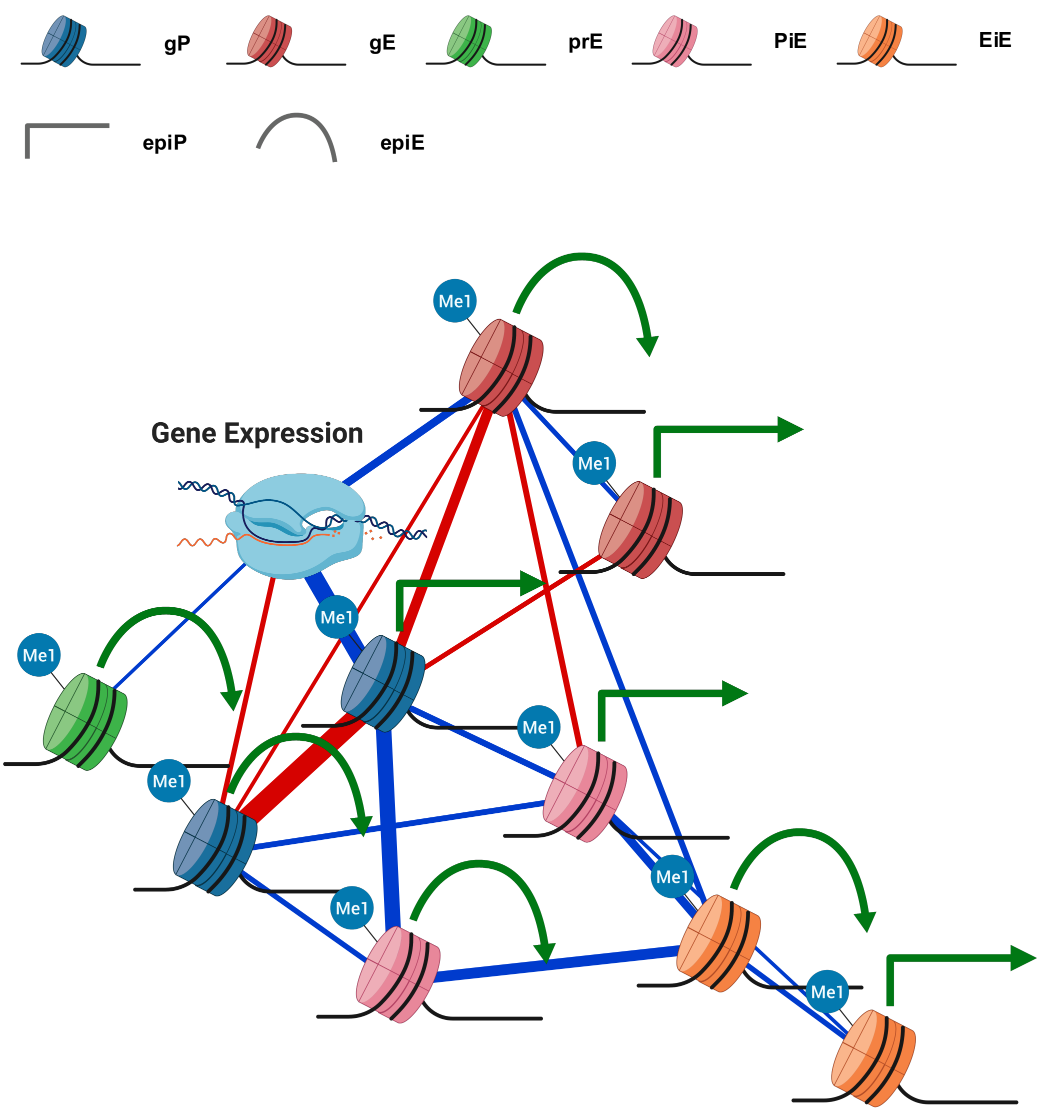


**Supplementary Figure 42. H3K4me1 Sparse Partial Correlation Network**. Blue edges represent positive partial correlations and red edges negative ones. Edge widths are proportional to absolute partial correlation values within each network. Only nodes for values with significant and relevant partial correlations (see Methods) were represented (minimal partial correlation = -0.431; maximal partial correlation = 0.366; all partial correlations Benjamini-Hochberg's Q-value < 7.7 x $10^{-67}$).

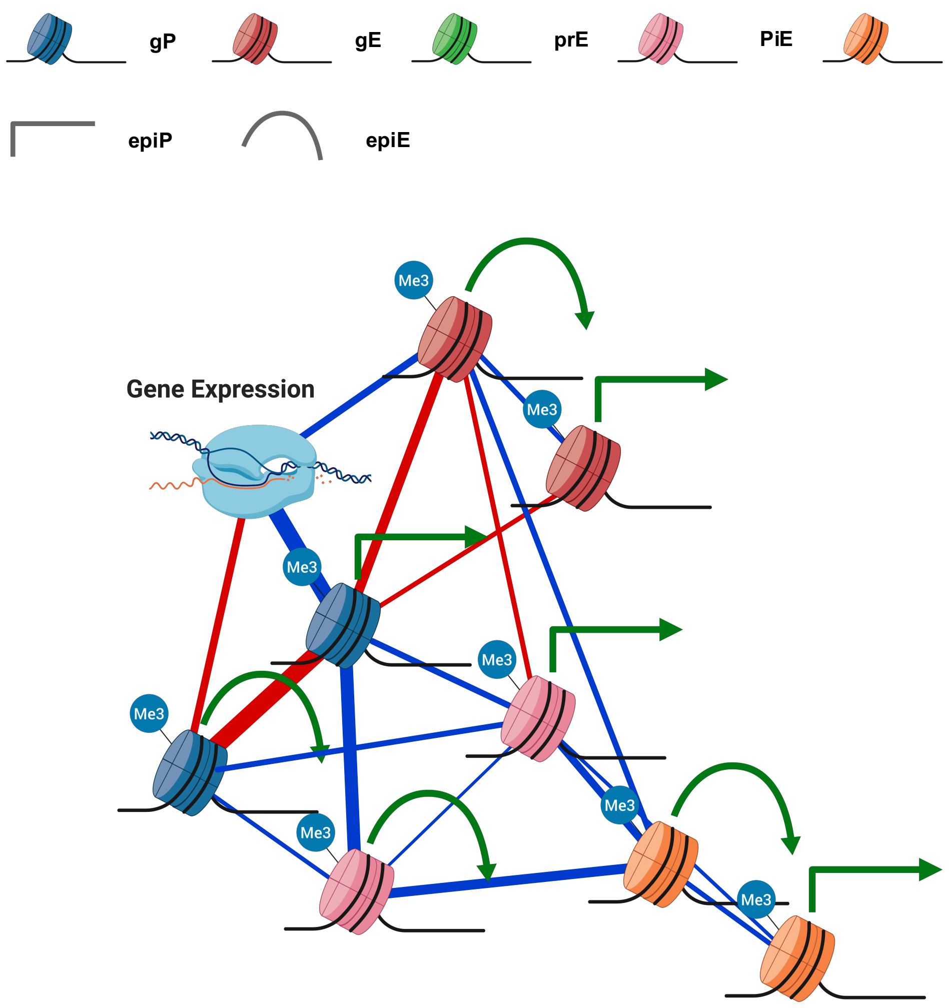


**Supplementary Figure 43. H3K4me3 Sparse Partial Correlation Network**. Blue edges represent positive partial correlations and red edges negative ones. Edge widths are proportional to absolute partial correlation values within each network. Only nodes for values with significant and relevant partial correlations (see Methods) were represented (minimal partial correlation = -0.388; maximal partial correlation = 0.358; all partial correlations Benjamini-Hochberg's Q-value < 1.2 x $10^{-56}$).

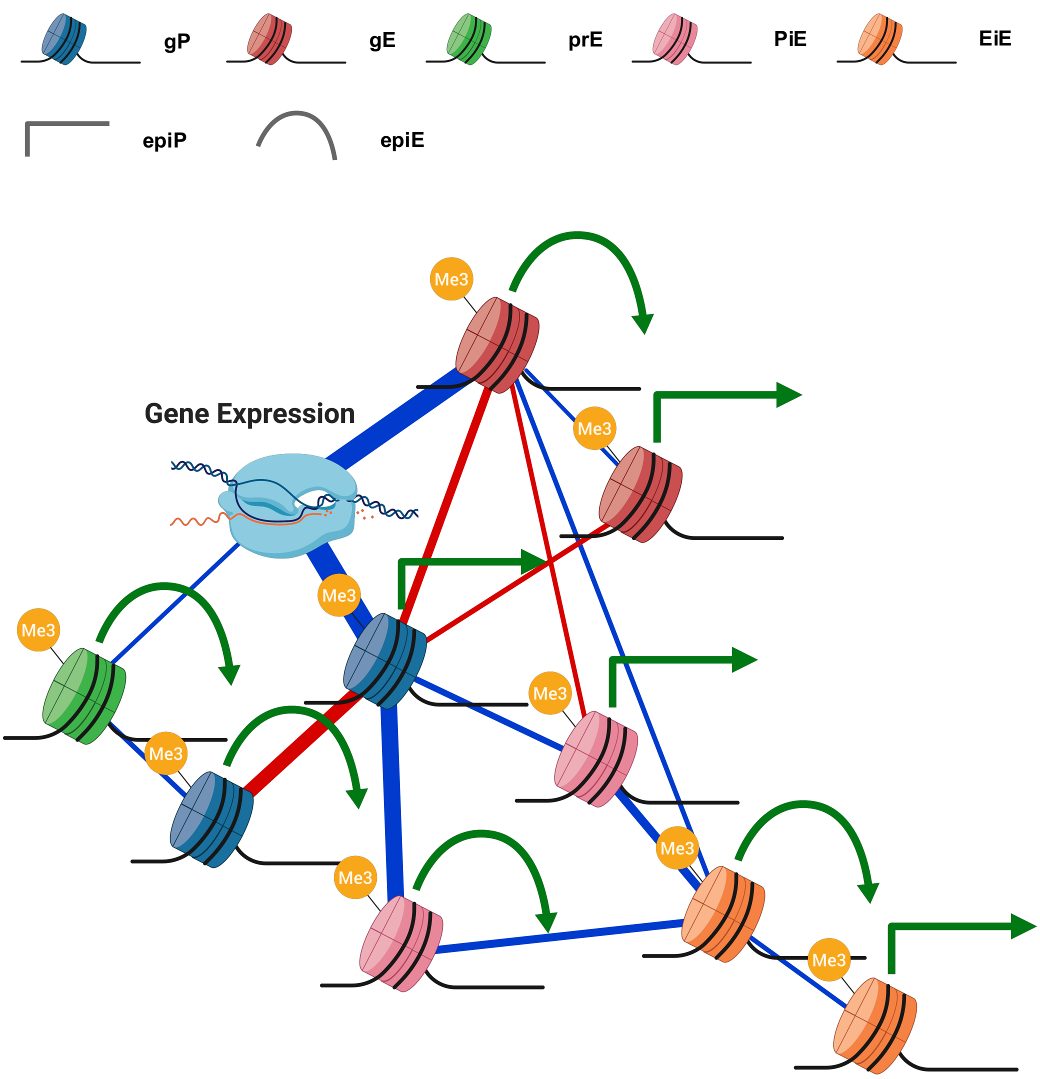


**Supplementary Figure 44. H3K36me3 Sparse Partial Correlation Network**. Blue edges represent positive partial correlations and red edges negative ones. Edge widths are proportional to absolute partial correlation values within each network. Only nodes for values with significant and relevant partial correlations (see Methods) were represented (minimal partial correlation = -0.289; maximal partial correlation = 0.498; all partial correlations Benjamini-Hochberg's Q-value < 1.5 x $10^{-84}$).

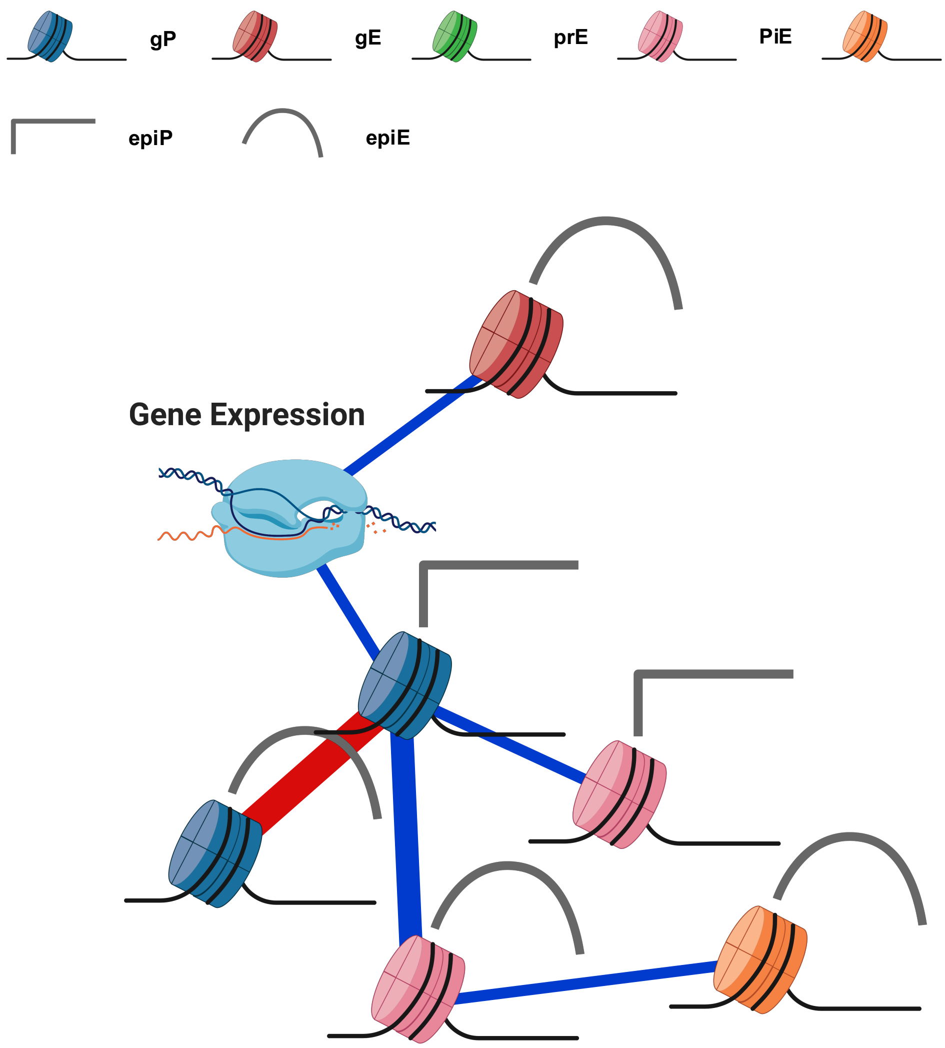


**Supplementary Figure 45. Global eigencomponents Sparse Partial Correlation Network for genes with a full architecture**. Blue edges represent positive partial correlations and red edges negative ones. Edge widths are proportional to absolute partial correlation values within each network. Only nodes for values with significant and relevant partial correlations (see Methods) were represented (minimal partial correlation = -0.581; maximal partial correlation = 0.448; all partial correlations Benjamini-Hochberg's Q-value < $3.3 \times 10^{--97}$).

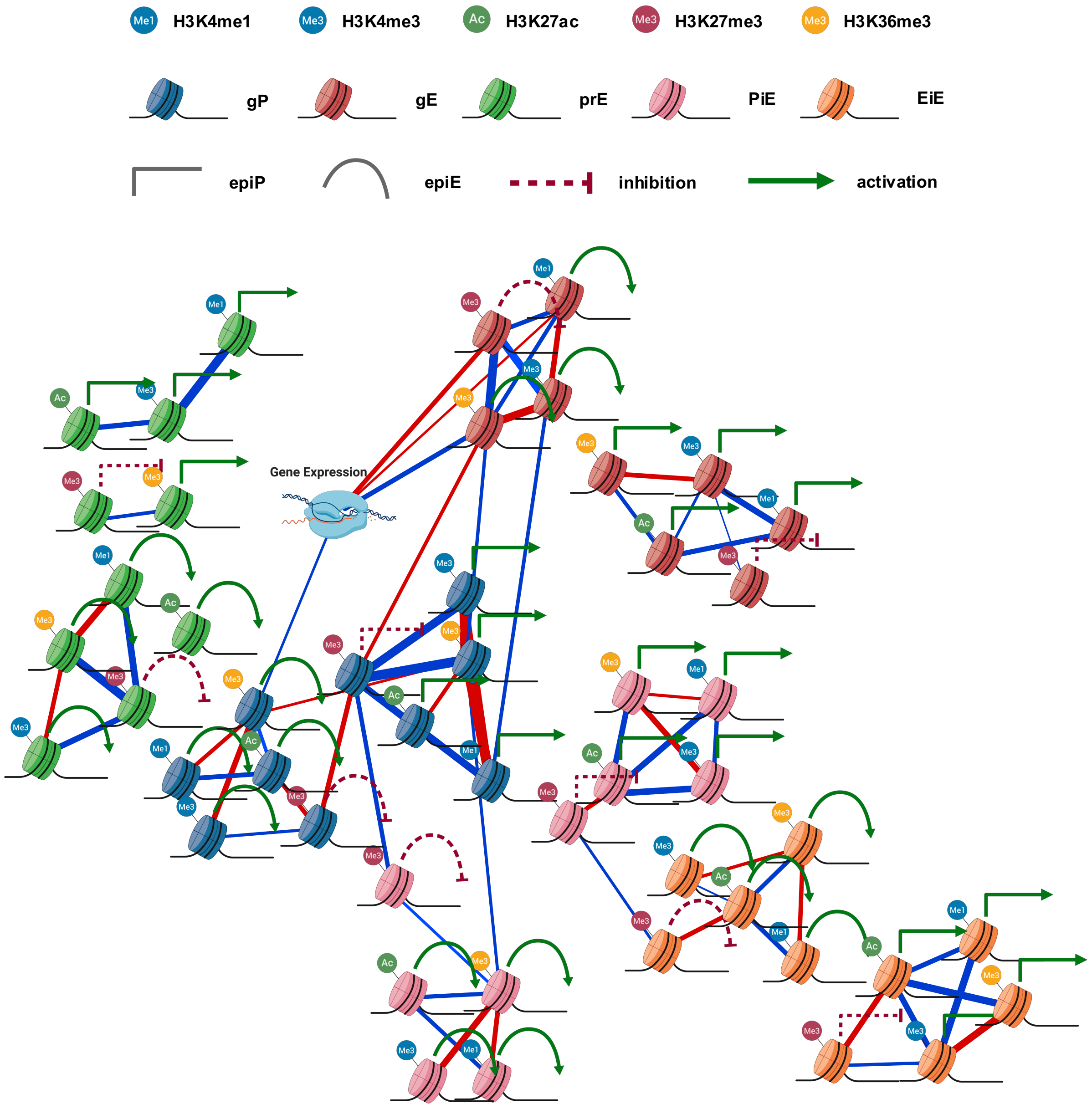


**Supplementary Figure 46. Global residuals Sparse Partial Correlation Network for genes with a full architecture** (minimal partial correlation = -0.581; maximal partial correlation = 0.448; all partial correlations Benjamini-Hochberg's Q-value < 3.3 x $10^{--97}$). Blue edges represent positive partial correlations and red edges negative ones. Edge widths are proportional to absolute partial correlation values within each network. Only nodes for values with significant and relevant partial correlations (see Methods) were represented.

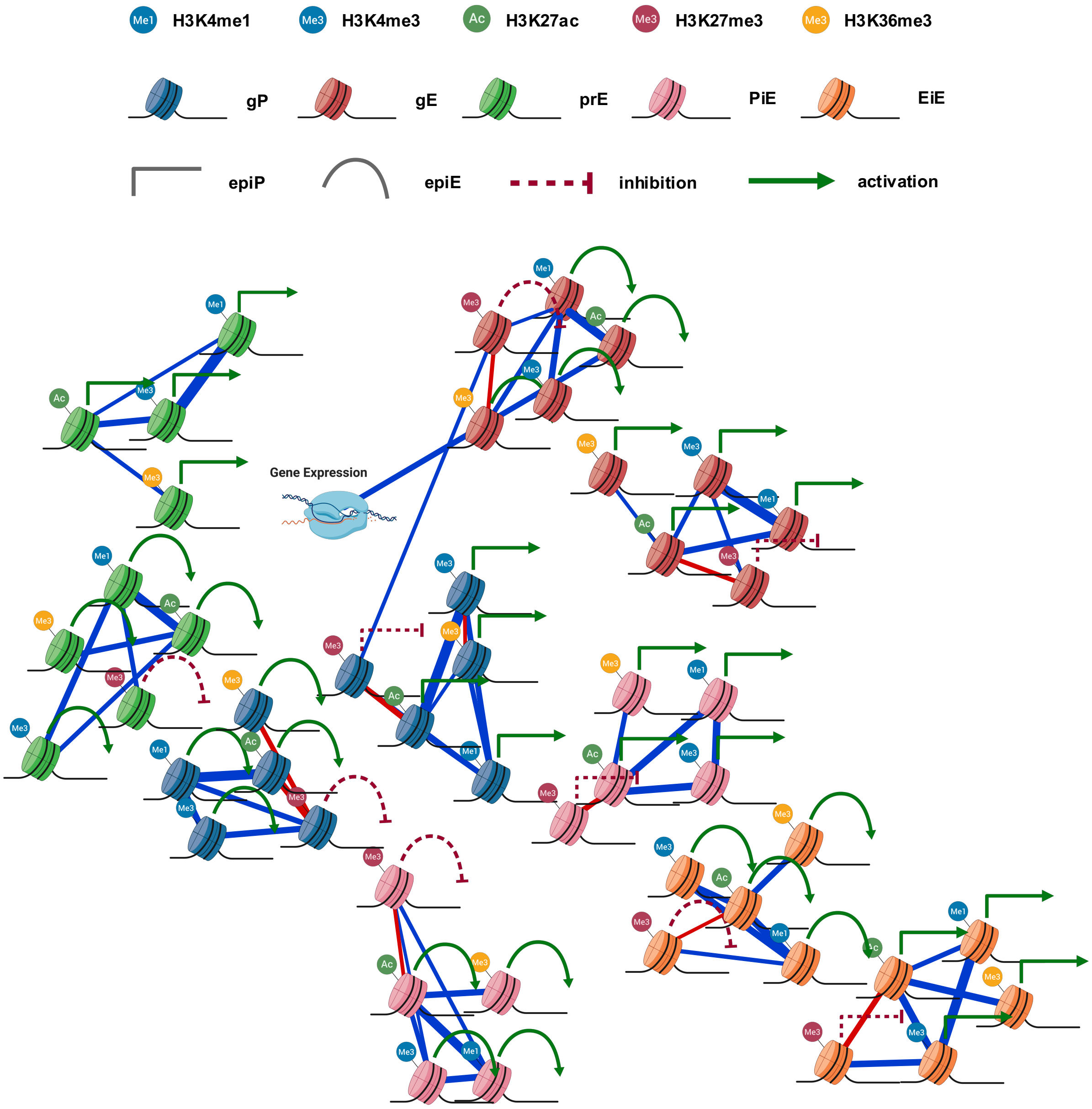


**Supplementary Figure 47. Global Sparse Partial Correlation Network for genes with a full architecture**. Blue edges represent positive partial correlations and red edges negative ones. Edge widths are proportional to absolute partial correlation values within each network. Only nodes for values with significant and relevant partial correlations (see Methods) were represented (minimal partial correlation = -0.442; maximal partial correlation = 0.734; all partial correlations Benjamini-Hochberg's Q-value < $4.7 \times 10^{-110}$).

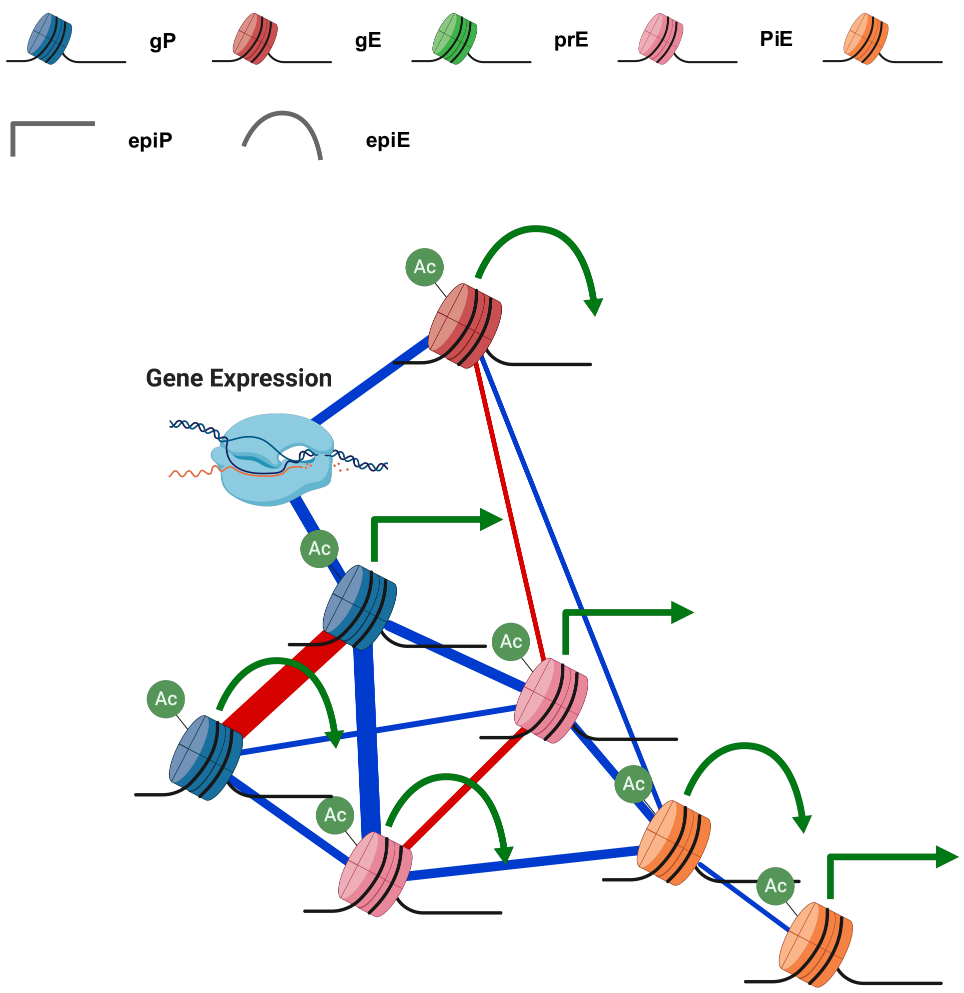


**Supplementary Figure 48. H3K27ac Sparse Partial Correlation Network for genes with a full architecture**. Blue edges represent positive partial correlations and red edges negative ones. Edge widths are proportional to absolute partial correlation values within each network. Only nodes for values with significant and relevant partial correlations (see Methods) were represented (minimal partial correlation = -0.568; maximal partial correlation = 0.444; all partial correlations Benjamini-Hochberg's Q-value < $9.5 \times 10^{-26}$).

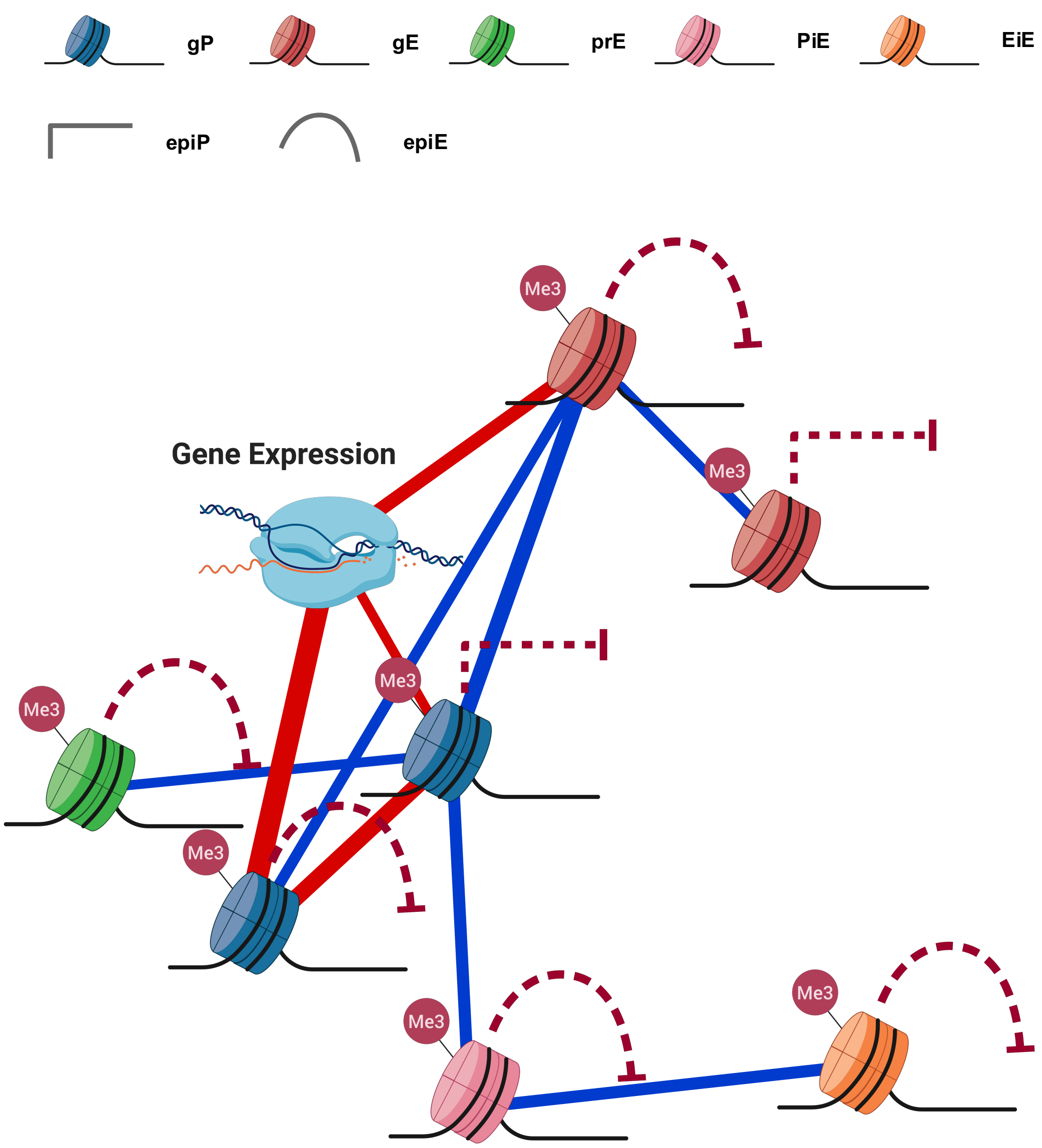


**Supplementary Figure 49. H3K27me3 Sparse Partial Correlation Network for genes with a full architecture**. Blue edges represent positive partial correlations and red edges negative ones. Edge widths are proportional to absolute partial correlation values within each network. Only nodes for values with significant and relevant partial correlations (see Methods) were represented (minimal partial correlation = -0.363; maximal partial correlation = 0.259; all partial correlations Benjamini-Hochberg's Q-value < $2.4 \times 10^{-57}$).

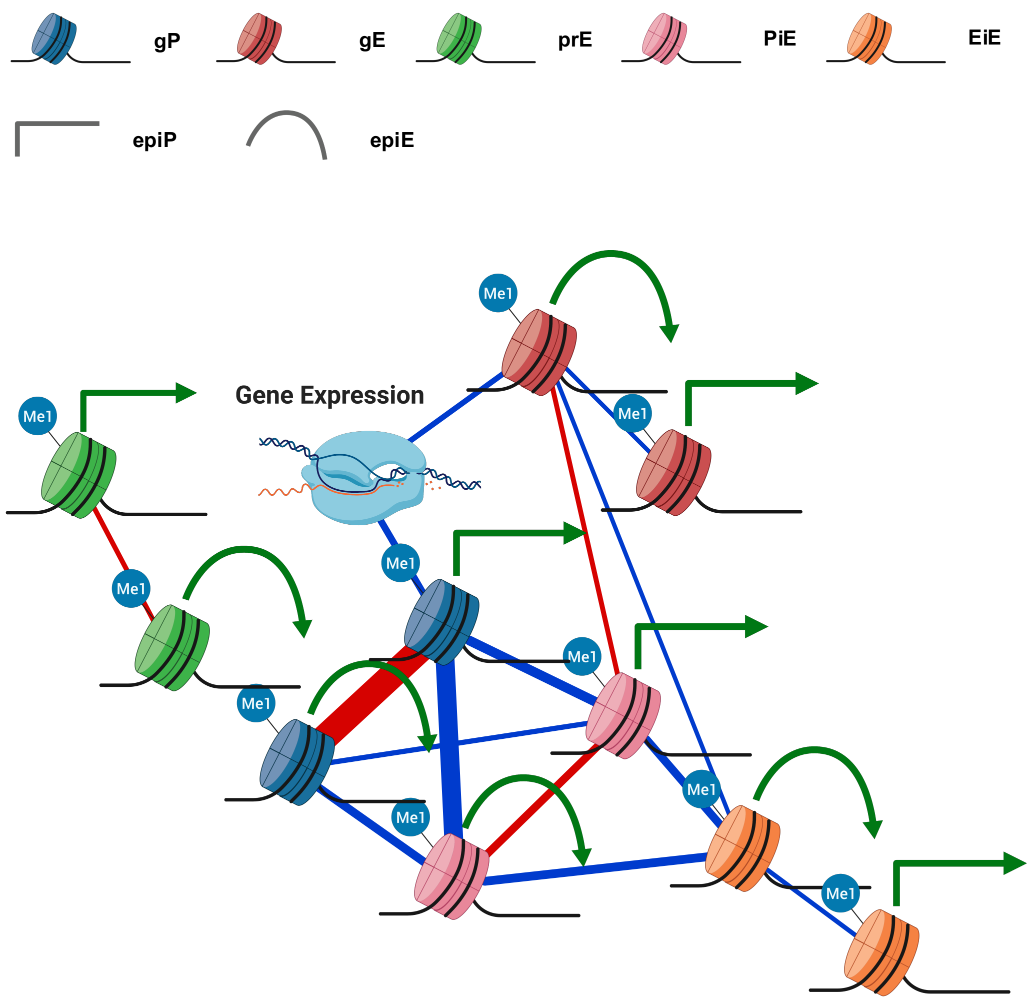


**Supplementary Figure 50. H3K4me1 Sparse Partial Correlation Network for genes with a full architecture**. Blue edges represent positive partial correlations and red edges negative ones. Edge widths are proportional to absolute partial correlation values within each network. Only nodes for values with significant and relevant partial correlations (see Methods) were represented (minimal partial correlation = -0.604; maximal partial correlation = 0.444; all partial correlations Benjamini-Hochberg's Q-value < $1.8 \times 10^{-22}$).

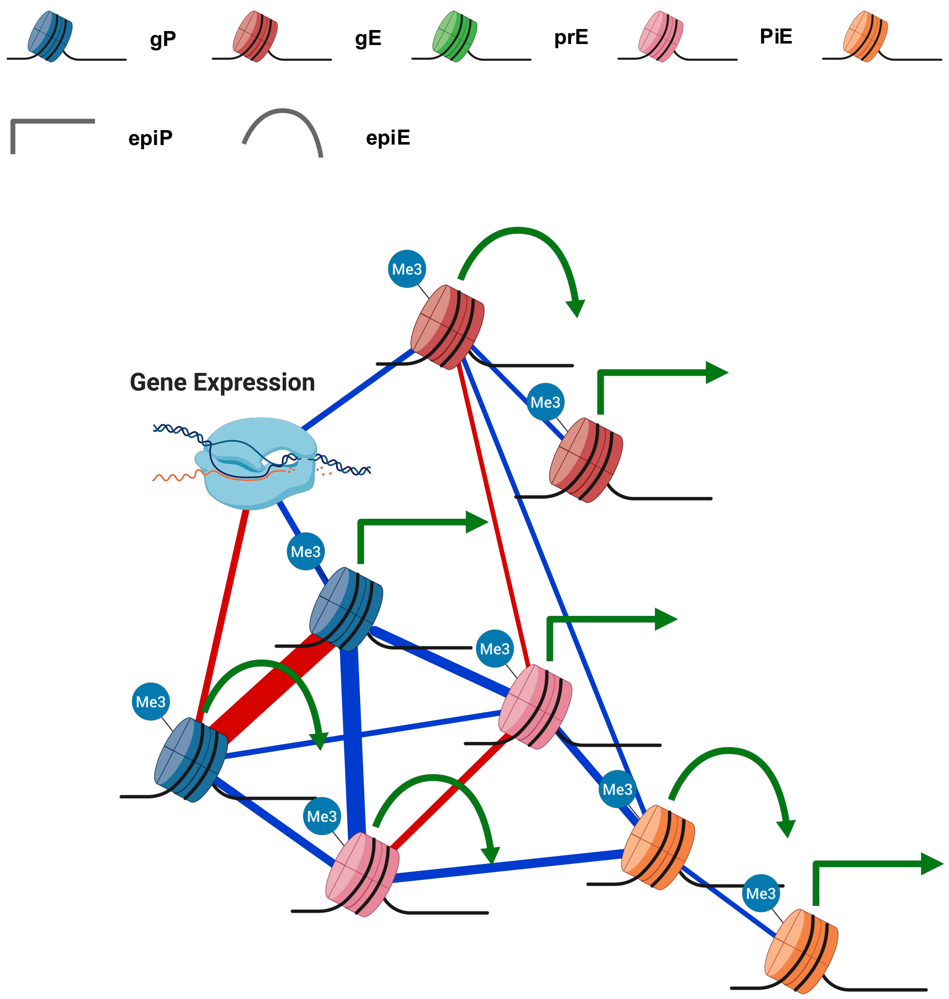


**Supplementary Figure 51. H3K4me3 Sparse Partial Correlation Network for genes with a full architecture**. Blue edges represent positive partial correlations and red edges negative ones. Edge widths are proportional to absolute partial correlation values within each network. Only nodes for values with significant and relevant partial correlations (see Methods) were represented (minimal partial correlation = -0.572; maximal partial correlation = 0.414; all partial correlations Benjamini-Hochberg's Q-value < $8.5 \times 10^{-23}$).

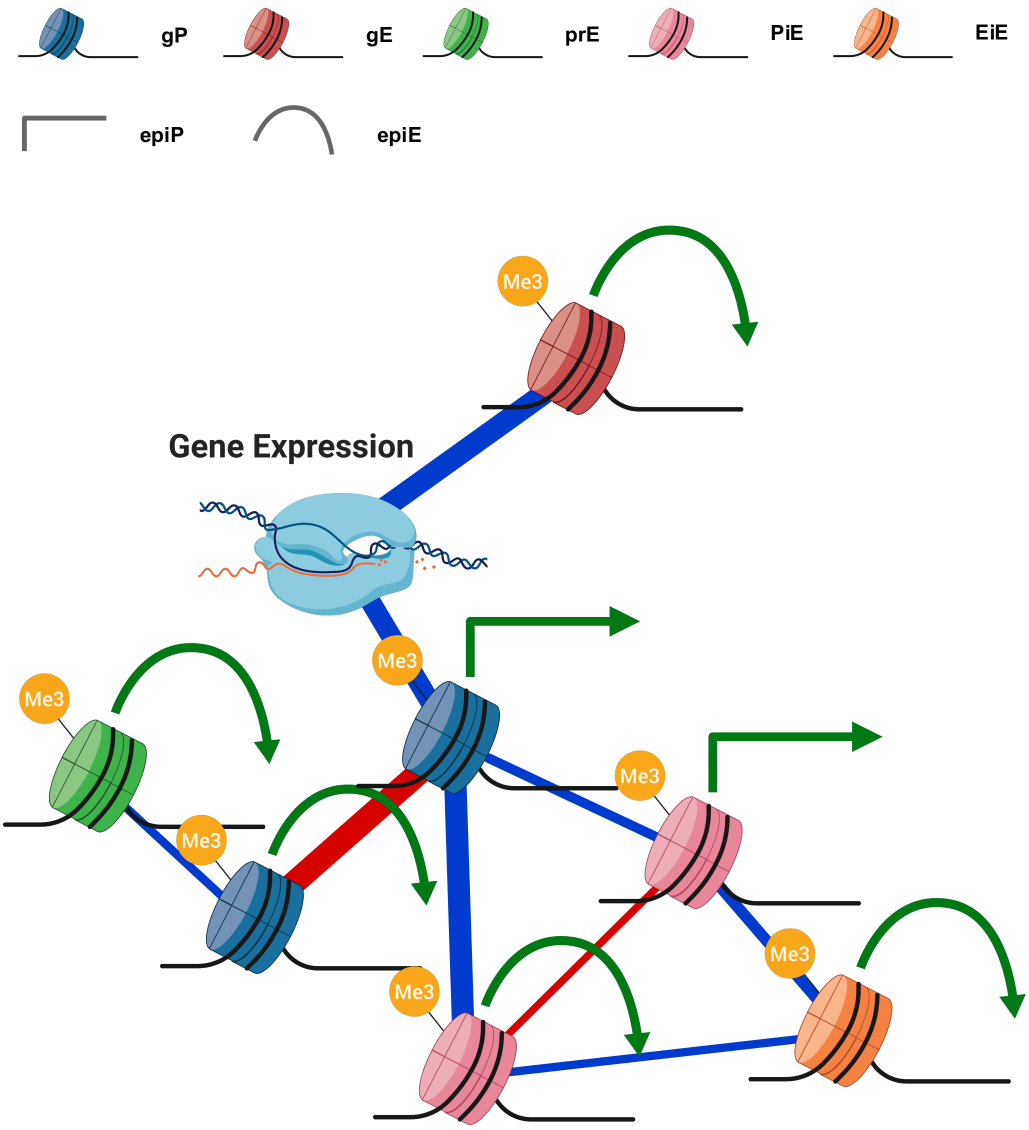


**Supplementary Figure 52. H3K36me3 Sparse Partial Correlation Network for genes with a full architecture**. Blue edges represent positive partial correlations and red edges negative ones. Edge widths are proportional to absolute partial correlation values within each network. Only nodes for values with significant and relevant partial correlations (see Methods) were represented (minimal partial correlation = -0.414; maximal partial correlation = 0.399; all partial correlations Benjamini-Hochberg's Q-value < 2.3 x $10^{-37}$).

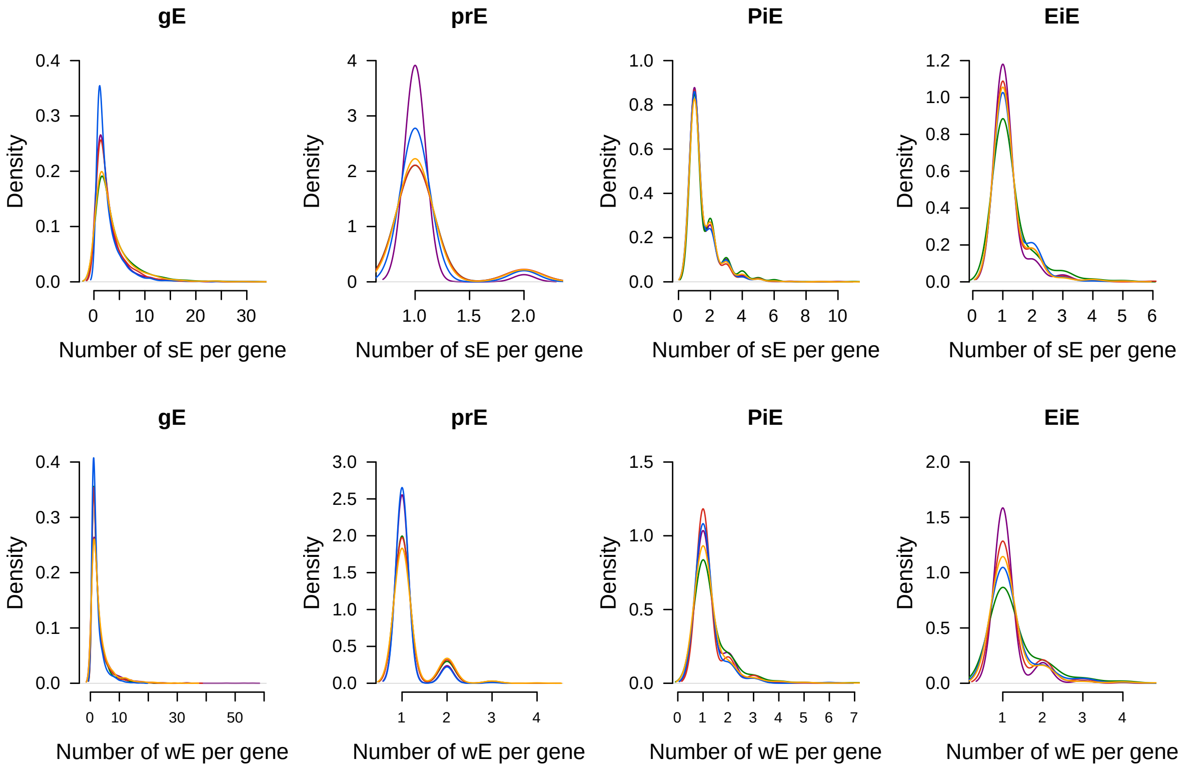


**Supplementary Figure 53. Distribution of the number of regulatory elements with strong and weak enhancers states associated with a gene, stratified by their regulatory component. a,** Number of sE in each enhancer-like regulatory component **b,** Number of wE in each enhancer-like regulatory component. Species are color-coded (Human: purple; Chimpanzee: green; Gorilla: red Orangutan: Blue; Macaque: Yellow).

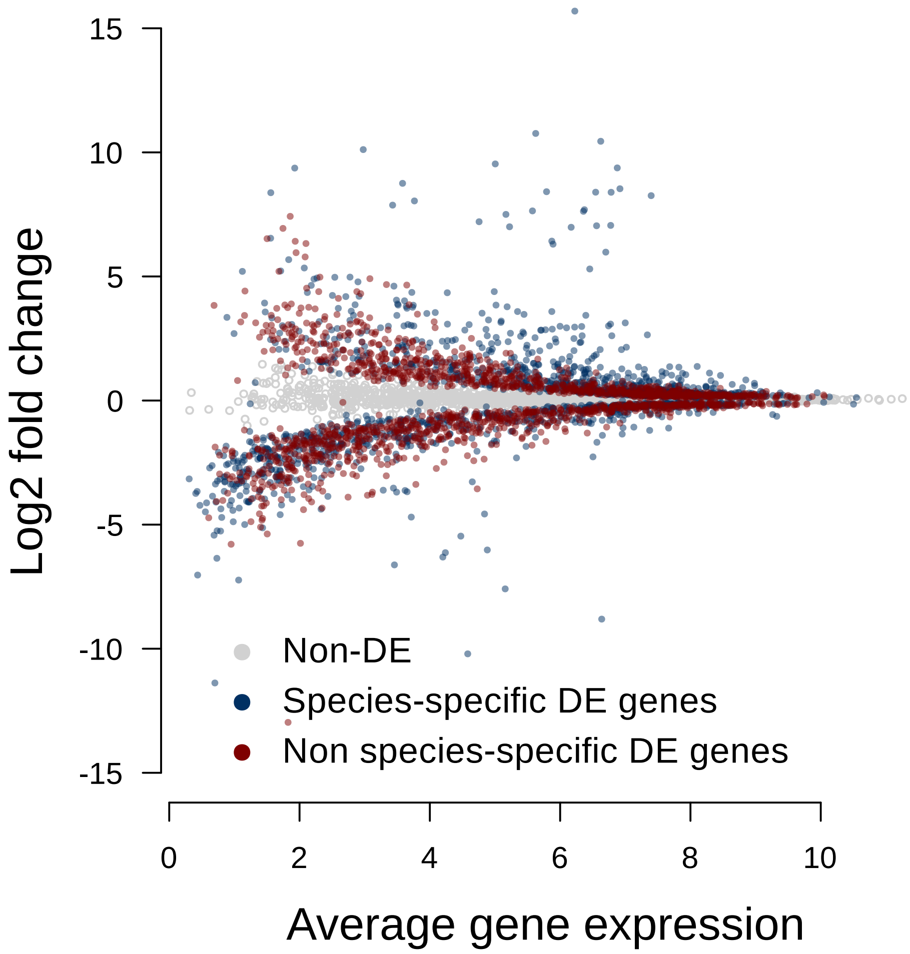


**Supplementary Figure 54. Genes with expression changes between species.** MA plot showing the average gene expression versus the expression change (log2 fold change) for genes non-differentially expressed between species (grey), genes with species-specific expression changes (blue) and genes with non-species-specific expression changes (red).

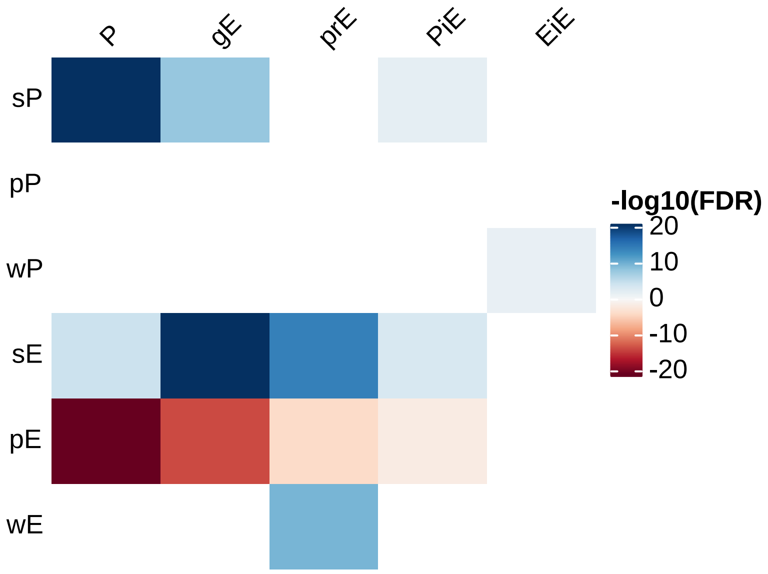


**Supplementary Figure 55. Gene expression differences between species are significantly associated with changes in the number of regulatory elements with strong and poised activities, mainly at genic promoters and intragenic enhancers.** Cell values in the heatmap show the adjusted P-value from a Wilcox signed rank-test testing whether a higher number of regulatory elements in the corresponding component-state combinations are significantly associated with higher (blue) or lower (red) expression levels in genes with expression differences across species.

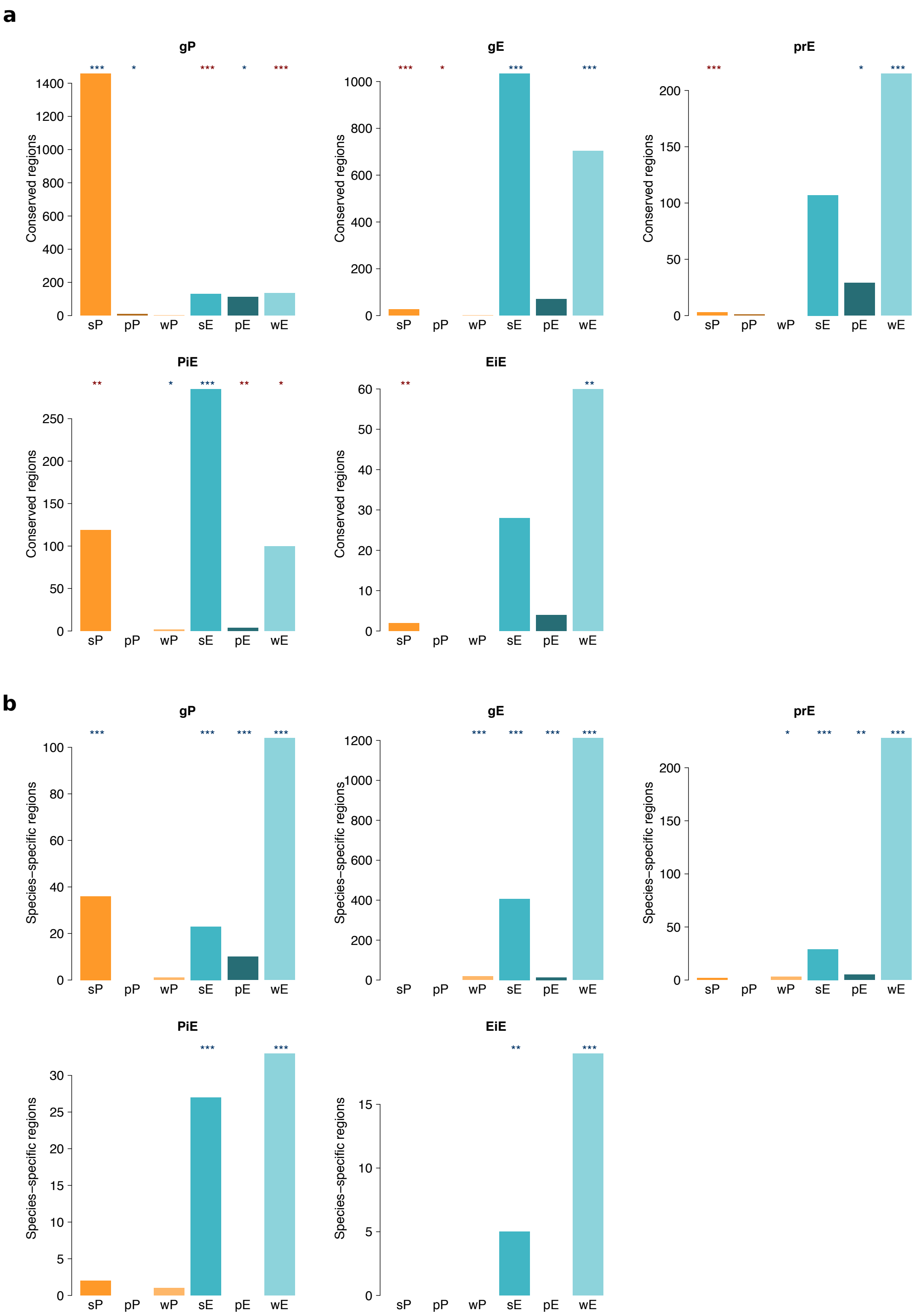


**Supplementary Figure 56. Evolutionarily conserved and species-specific regulatory states are enriched in particular regulatory components**. For each type of component, barplots show the number of **(a)** evolutionarily conserved or **(b)** species-specific orthologous regulatory regions with the corresponding color-coded regulatory state fully conserved. Asterisks represent the magnitude of a Chi-square test residuals, where blue and red indicate there are either more or fewer elements, respectively, in that particular component-state combination than expected by random chance (* abs(residulas) > 2; ** abs(residulas) > 4; *** abs(residuals) > 8; Chi-square test, $P < 2.2 \times 10^{-16}$).

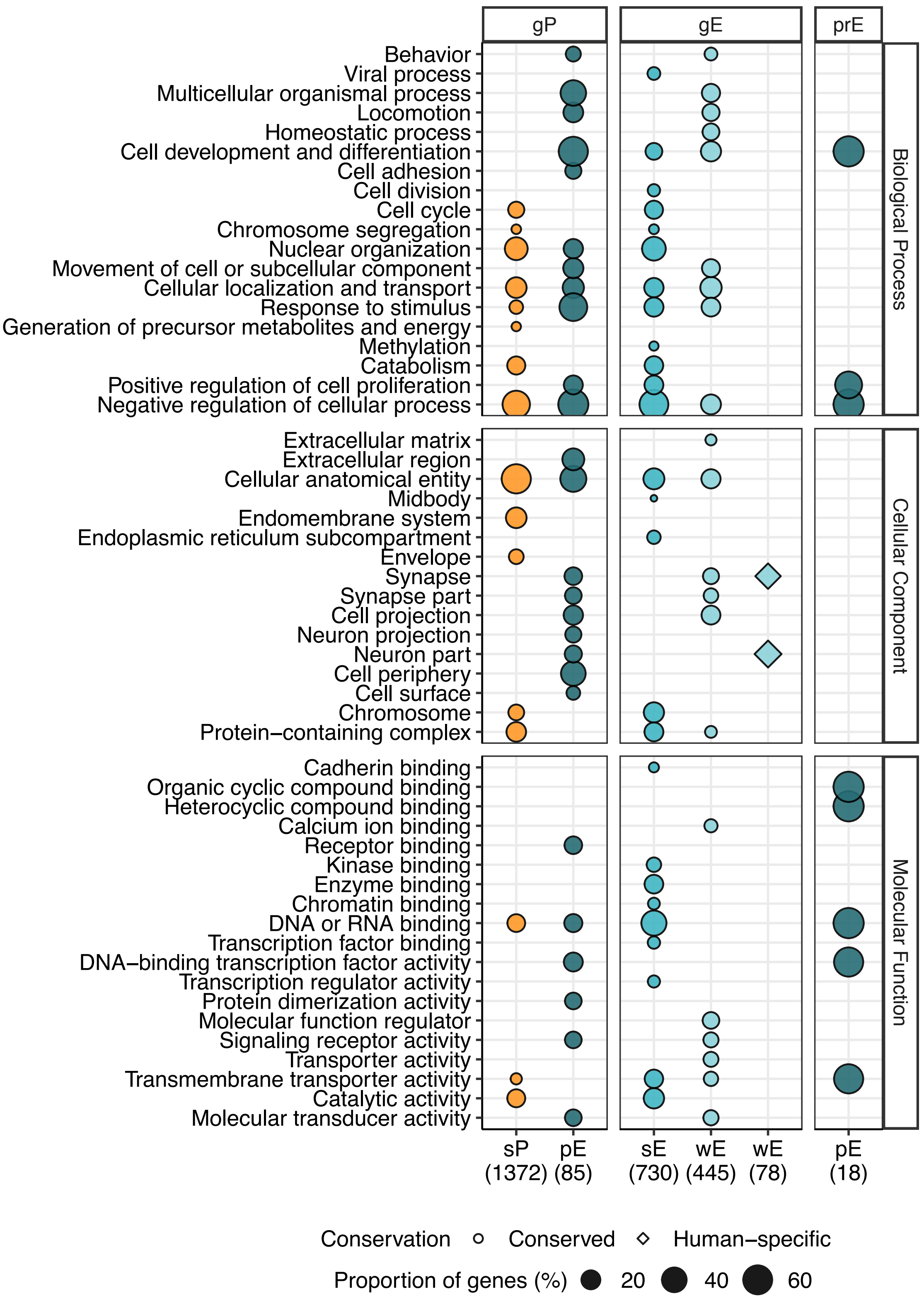


**Supplementary Figure 57. Functional enrichment of genes associated with fully conserved and species-specific epigenetic state/component combinations.** Functional enrichment of significantly enriched groups in Supplementary Figure 58. The size of circles/diamonds indicates the proportion of genes included in each functional category out of the total number of genes contained in the corresponding regulatory group (shown in brackets).

A simplified version of this figure including Biological Process and Cellular Component enrichments for genic promoter and intragenic enhancer combinations is shown in Fig. 5a.

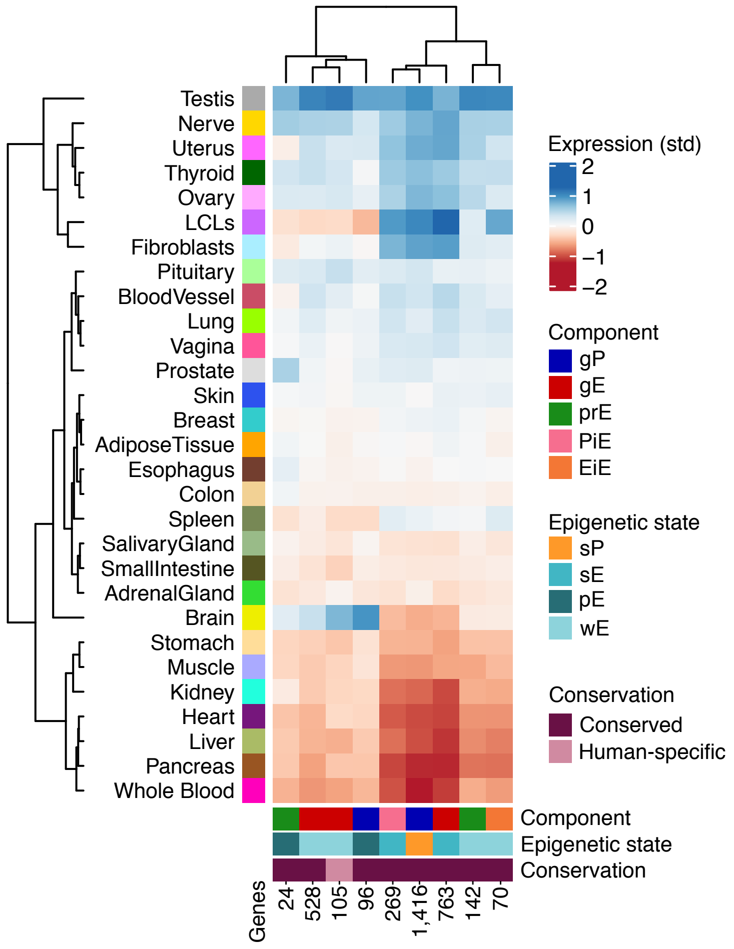


**Supplementary Figure 58. Heatmap of standardised expression of genes associated with fully conserved and species-specific epigenetic state/component combinations**. Epigenetic state/component combinations of enriched groups in Supplementary Figure 58. Standardised tissue expression pattern from bulk RNA-seq data obtained from GTEx (v8). A simplified version of this figure including combinations with significant functional enrichments is shown in Fig. 5b.

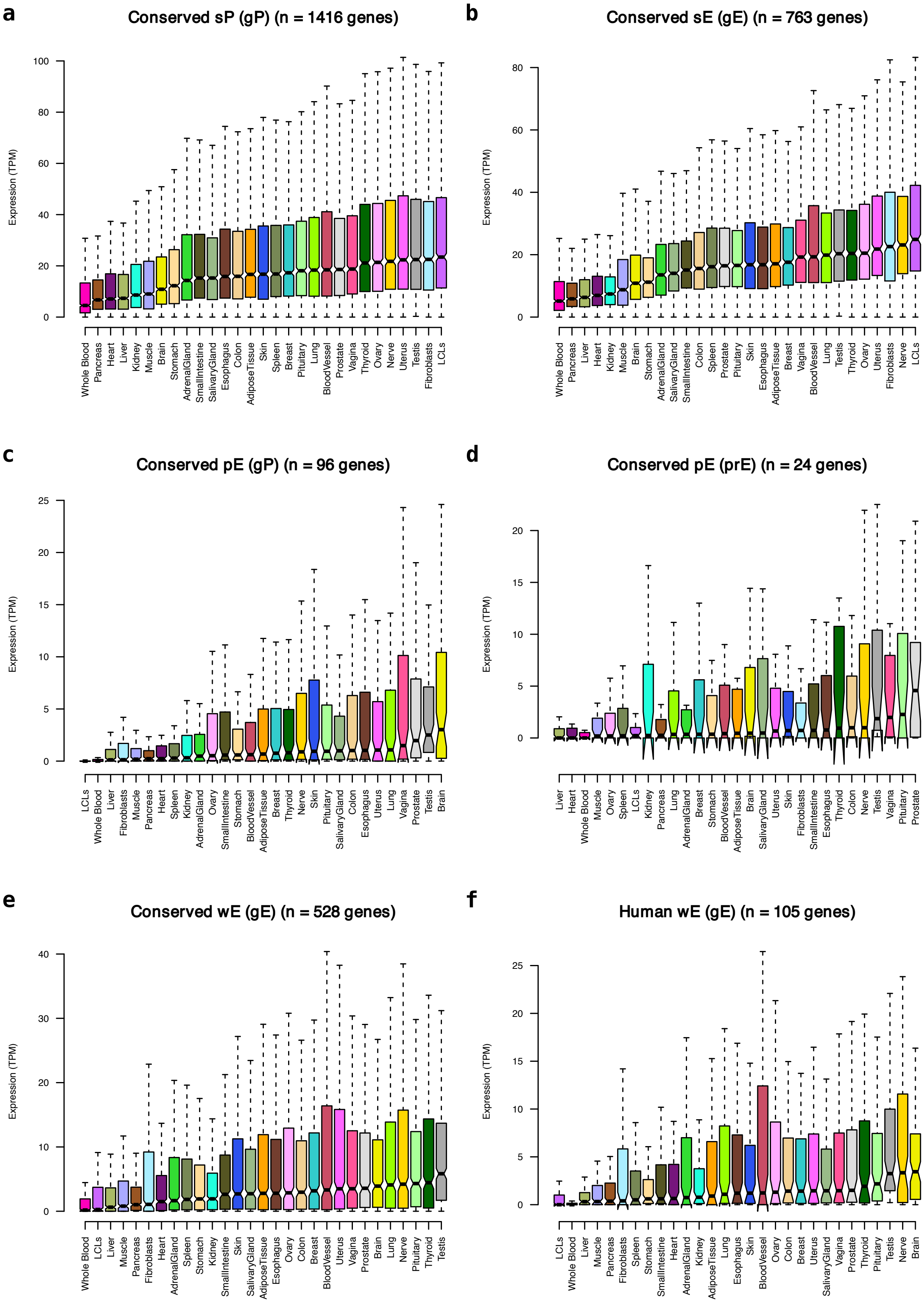


**Supplementary Figure 59. Tissue expression patterns of genes associated with particular conserved or human-specific component-state combinations with functional enrichments.** Tissue expression patterns (median TPM) from bulk RNA-seq data obtained from the latest GTEx release (v8).

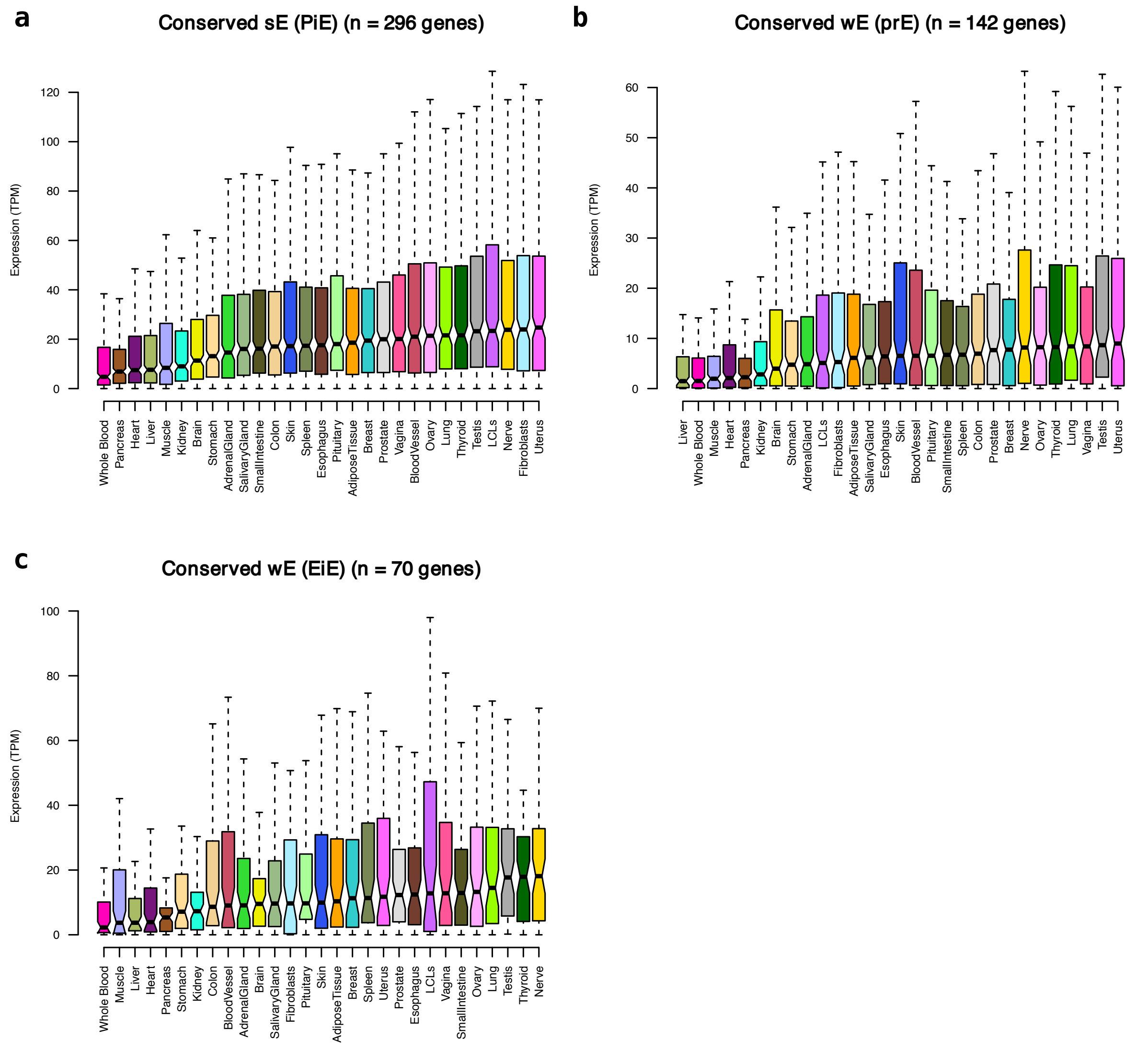


**Supplementary Figure 60. Tissue expression patterns of genes associated with particular conserved or human-specific component-state combinations without significant functional enrichments.** Tissue expression patterns (median TPM) from bulk RNA-seq data obtained from the latest GTEx release (v8).

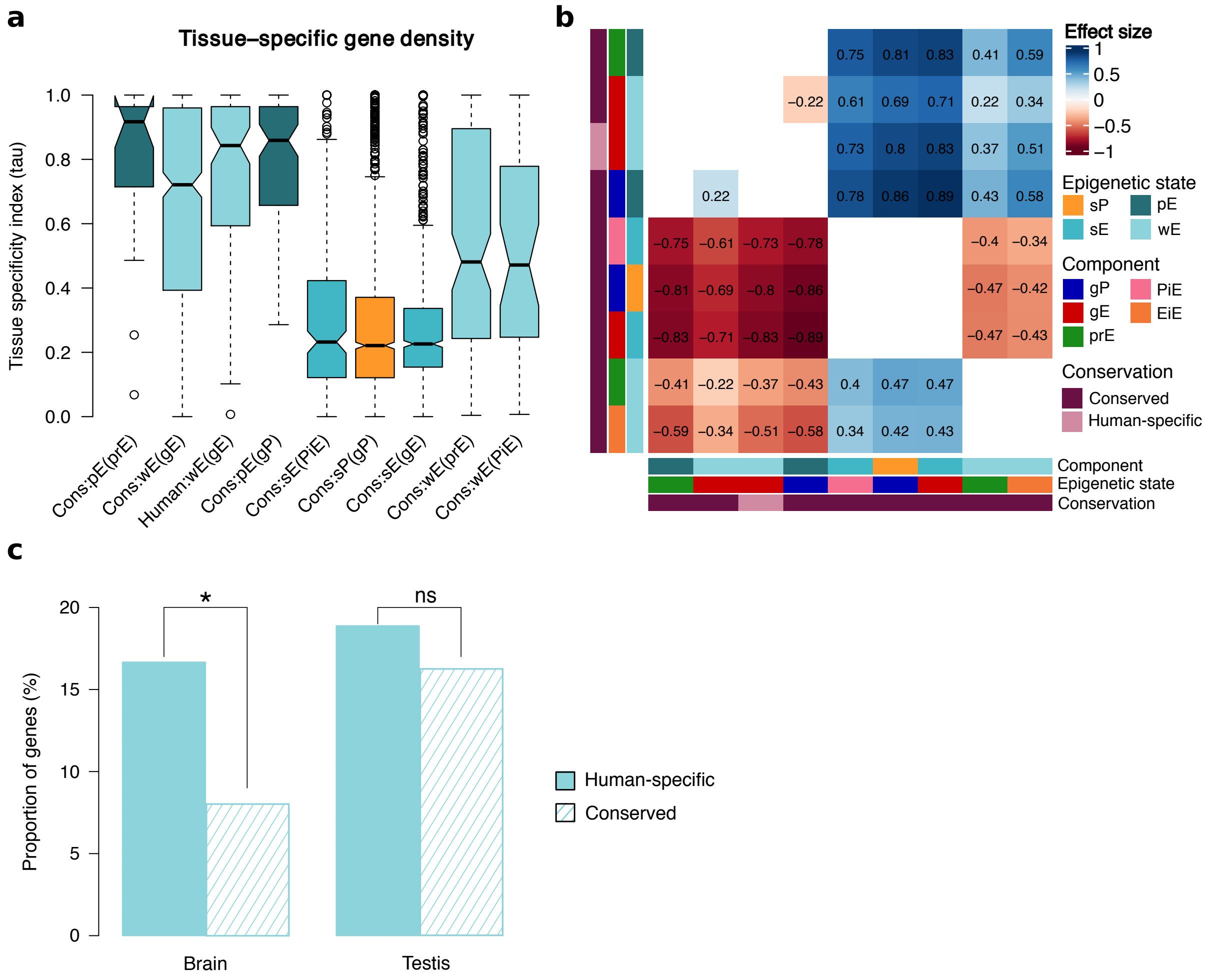


**Supplementary Figure 61. Tissue specificity analyses show different patterns across regulatory components and epigenetic states. a,** Distributions of the tissue-specific indices (tau) for genes associated with particular component-state combinations. **b,** Effect sizes corresponding to significant pairwise comparison from **(a)** (Dwass-Steel-Critchlow-Fligner test, $P < 0.05$) **c**, Comparison of the proportion of brain-specific (tau(Brain) > 0.8) and testis-specific genes (tau(testis) > 0.8) associated with either human-specific or conserved intragenic enhancers with weak enhancer states. Fisher's exact test: P-values below and above 0.05 are indicated by an asterisk (*) or ns, respectively.

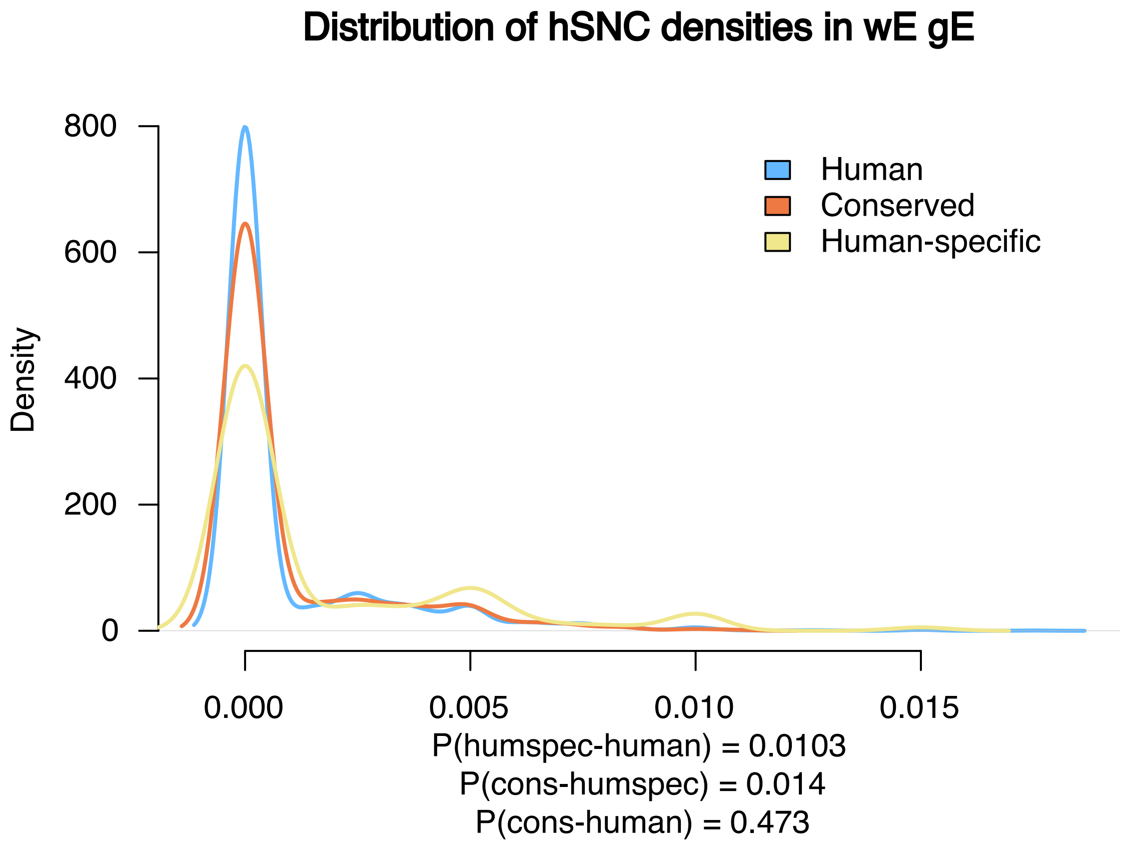


**Supplementary Figure 62**. **Human-specific intragenic enhancers with weak enhancer states are associated with a higher number of hSNCs.** Distribution of single nucleotide changes fixed in human and distinct from other primates (hSNC) in human genes associated with intragenic enhancers (yellow), in the subset of genes associated with fully conserved weak intragenic enhancers (orange) and in the subset of genes with human-specific weak intragenic enhancers (blue). P-values in the plot correspond to the indicated Mann-Whitney U tests.

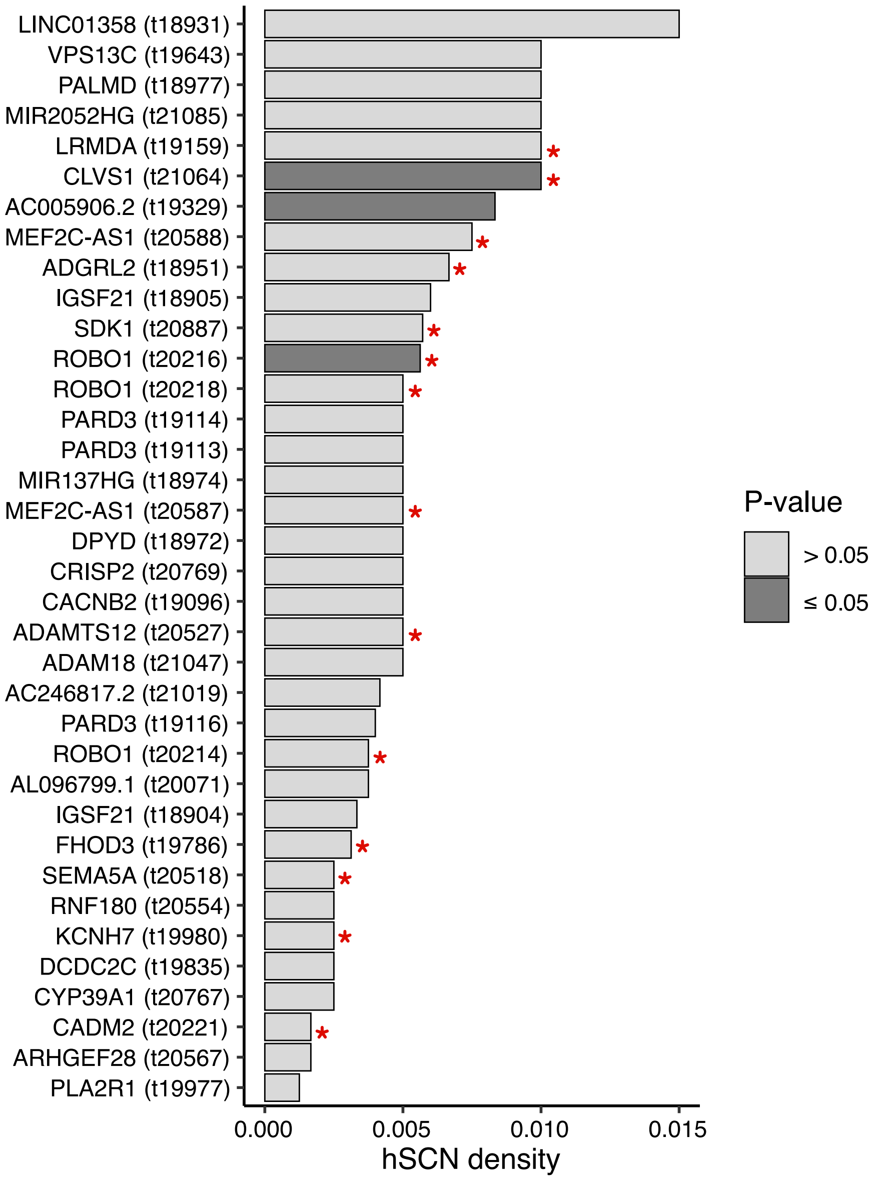


**Supplementary Figure 63. Human-specific intragenic enhancers with weak states enriched in human single nucleotide changes.** Barplots show the density of hSNCs in the 36 human-specific weak intragenic enhancers in 30 genes (enhancer identifier between brackets). Corresponding P-values were calculated through randomization analysis (Methods, 10,000 simulations). Red stars denote genes with signals of positive selection. Several human-specific intragenic weak enhancers can be contained in the same gene so that the same gene name can appear multiple times.

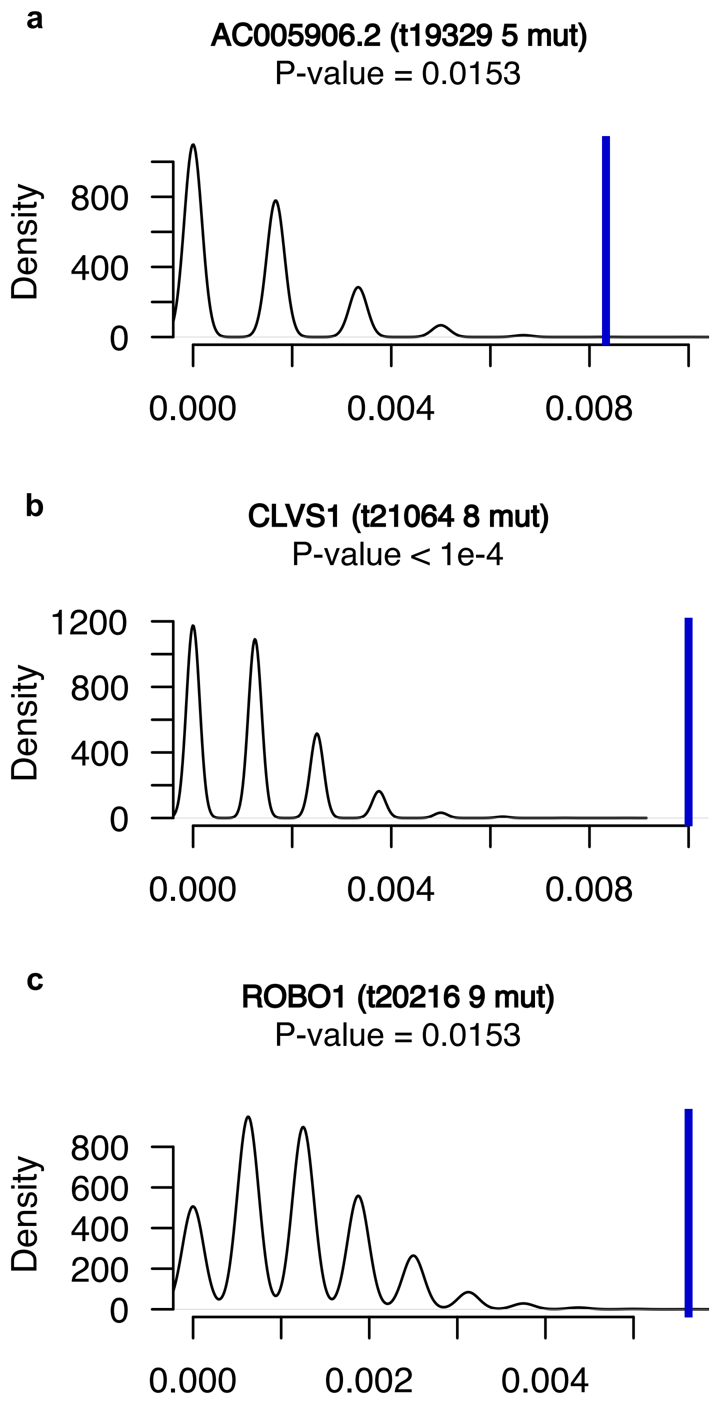


**Supplementary Figure 64. Three genes associated with human-specific intragenic enhancers with weak enhancer states accumulate more hSNCs than expected.** Distribution of simulated hSNCs densities (10,000 simulations) for the three statistically significant human-specific weak intragenic enhancers (or hits) in **a,** *AC005906.2* transcript and **b,** *CLVS1* and **c,** *ROBO1* protein-coding genes. The blue vertical line represents the value of the observed mutation density for each particular enhancer.

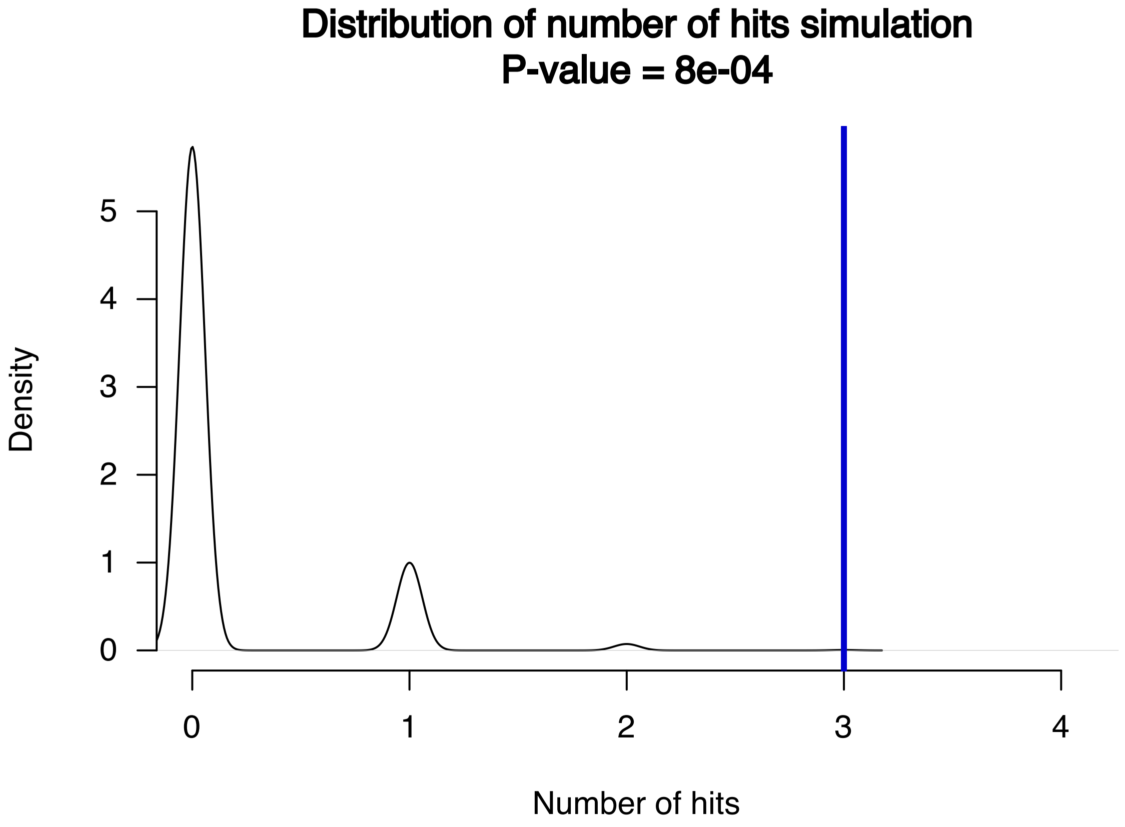


**Supplementary Figure 65. Significance of the observed number of hits.** Distribution of the simulated number of hits (number of human-specific weak intragenic enhancers with a density of hSNC higher than expected) obtained after 10,000 simulations. The blue vertical line represents the observed number of hits in our data.

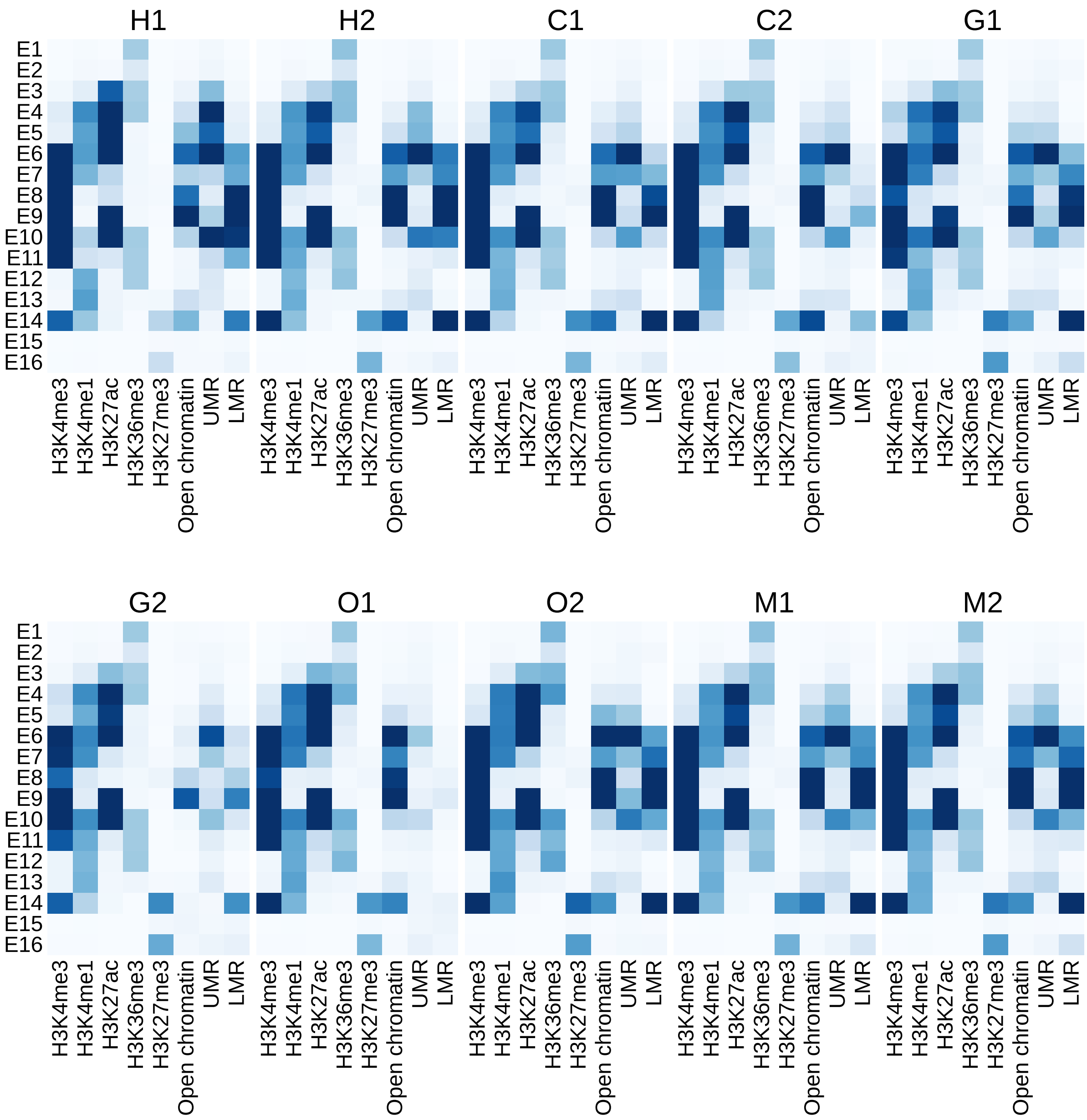


**Supplementary Figure 66. Histone and open chromatin peaks are enriched in the expected chromatin states.** See Supplementary Fig. 2.

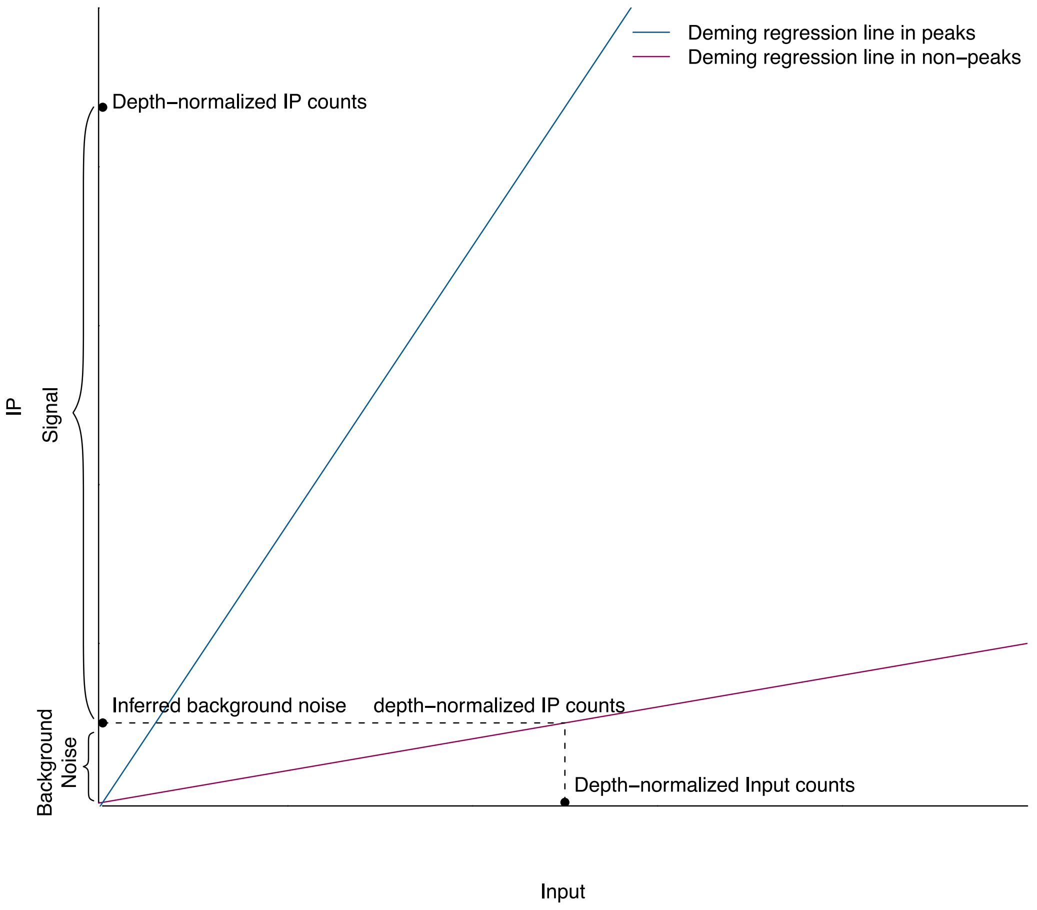


**Supplementary Figure 67. Schematic representation of how background noise signal is removed from the immunoprecipitated (IP) signal.** Regression parameters were estimated in a robust set of sample peaks and non-peaks and used later on to adjust the IP signal recovered at regulatory elements and orthologous regulatory regions (Supplementary Methods).

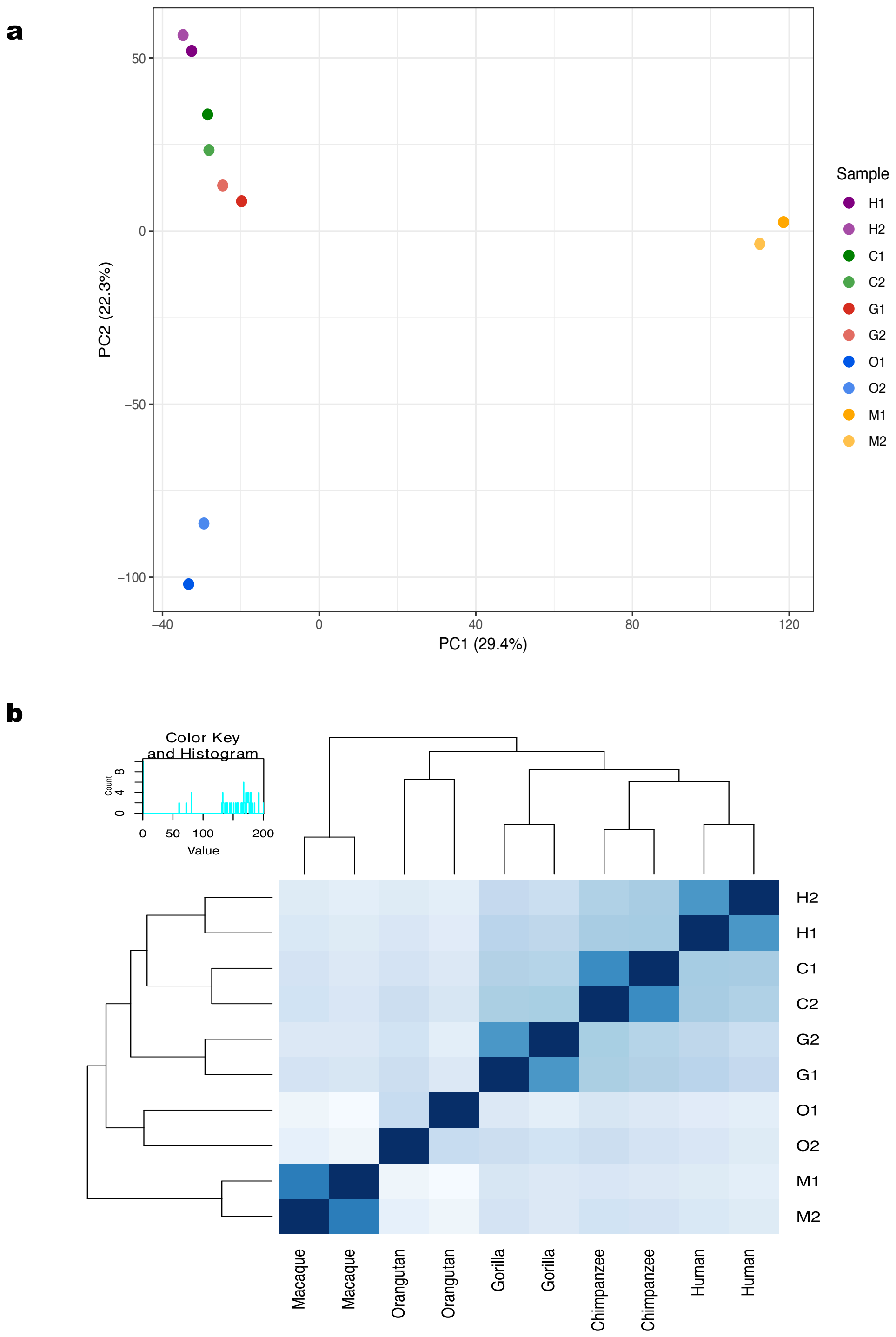


**Supplementary Figure 68. Gene expression patterns recapitulate the known phylogenetic relationships between species. a,** PCA and **b,** heatmap of the sample-to-sample Euclidean distances based on the expression levels of 1-to-1 orthologous protein-coding genes.

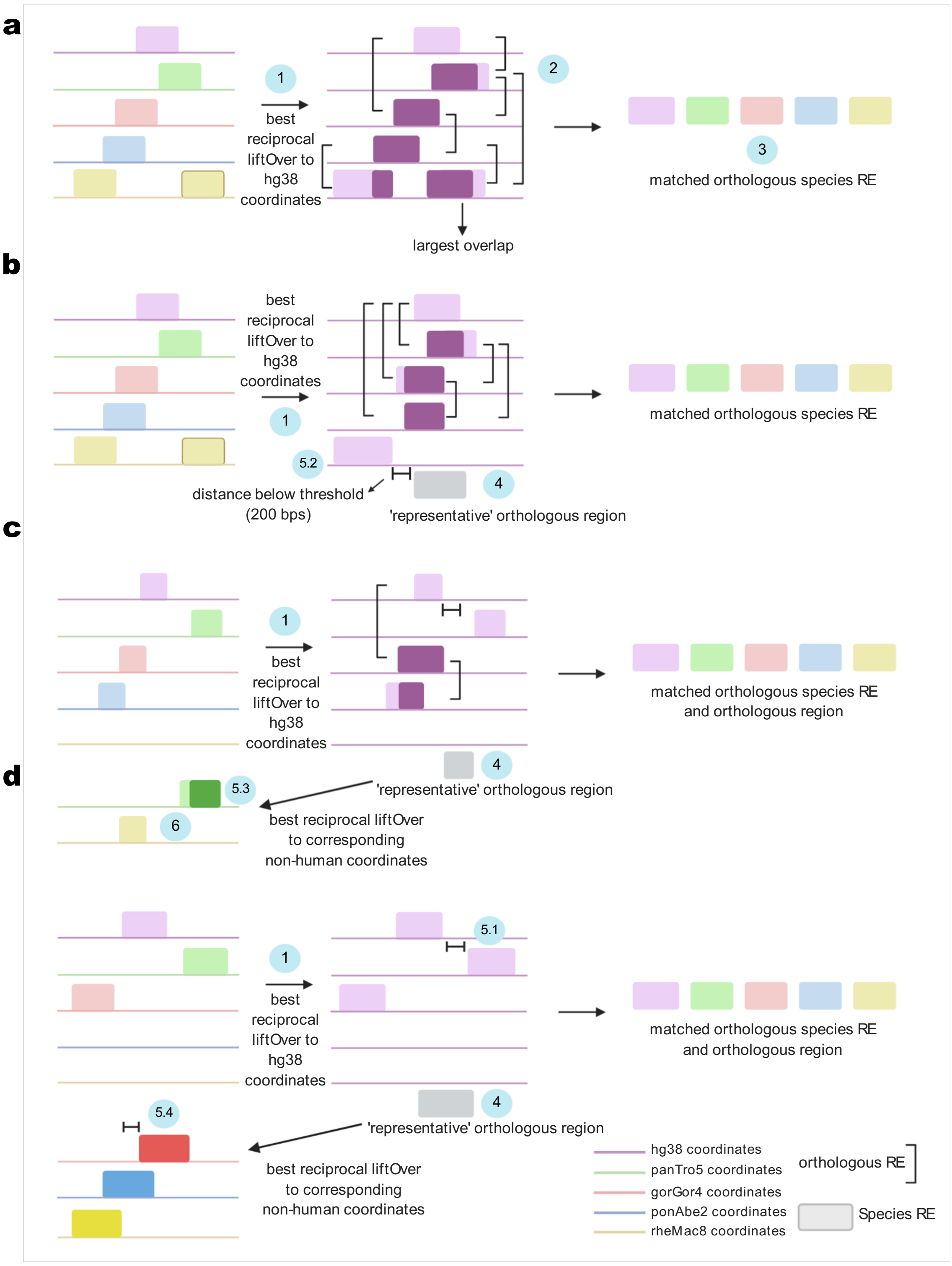


**Supplementary Figure 69. Schematic representation of how orthologous relationships among species regulatory elements were established.** Several different scenarios are illustrated. Boxes represent regulatory elements (RE). Colors represent the different reference genome coordinates (purple: human; green: chimpanzee; red: gorilla; blue: orangutan; yellow: macaque). Dark colors indicate overlapping bp. Outlined boxes with color gradient denote orthologous regions. Numbers in circles refer to the steps described in the text. **a,** Pairwise overlaps are found for all species. **b**, Although the macaque RE does not overlap with the REs of the other species, the representative orthologous region is in close proximity to the macaque RE which is then recovered as the corresponding orthologous. **c,** Neither the chimpanzee nor macaque REs overlap with REs from the remaining species. When the representative orthologous region is mapped to the chimpanzee reference genome, a RE is found in close proximity and recovered as orthologous RE. When the representative orthologous region is mapped to the macaque reference genome, no RE is found but the coordinates can be mapped and this genomic region is recovered as the corresponding orthologous region in macaque. In downstream quantitative analysis, reads will be counted and normalized in defined orthologous regions despite absence of RE. This approach allows the recovery of overlooked RE. **d,** Gorilla orthologous RE is recovered through proximity to the reference orthologous RE in gorilla coordinates. Orangutan and macaque are assigned their corresponding orthologous regions.

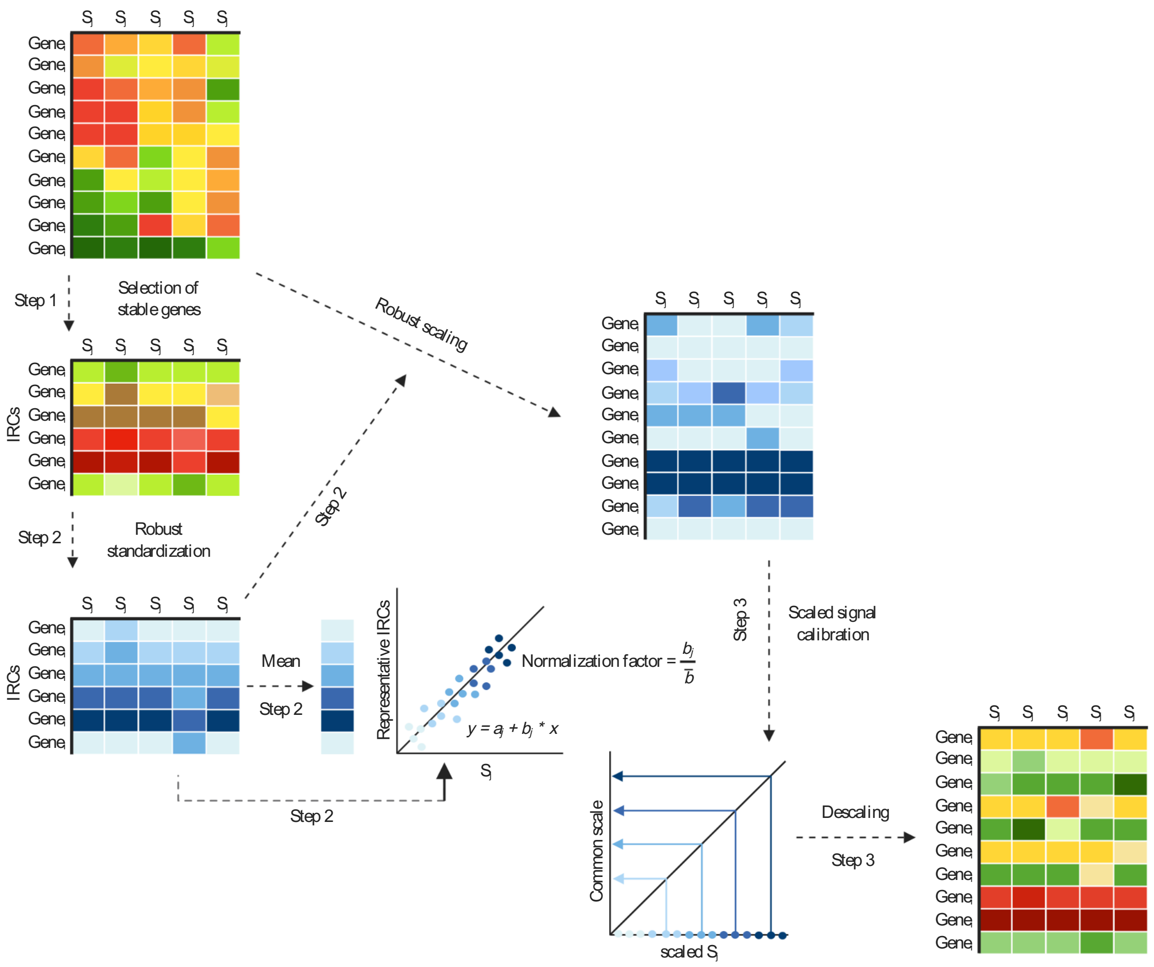


**Supplementary Figure 70. Schematic representation of the approach employed to calibrate the expression and epigenetic signals between species.** Step 1. Color gradient represents different degrees of expression/enrichments (green: low expression/enrichment; yellow: medium expression /enrichment; red: high expression/enrichment). IRCs are composed of units (genes, regulatory elements, epigenetic signals associated with a particular type of gene component) with similar enrichments across samples. Step 2. After robust standardization, values are normally distributed and more similar across samples (blue gradient; light blue: low enrichment; blue: medium enrichment; dark blue: high enrichment). Normalization factors are estimated considering the linear relationships between sample-specific and representative robust standardized values at IRCs. Step 3. Values at genes/elements/components are scaled to the common space where normalization factors are computed, and after the calibration values are brought back to the original scale. S stands for sample and j ∈ [1,10]. N is the number of observations and I ∈ [1,N].

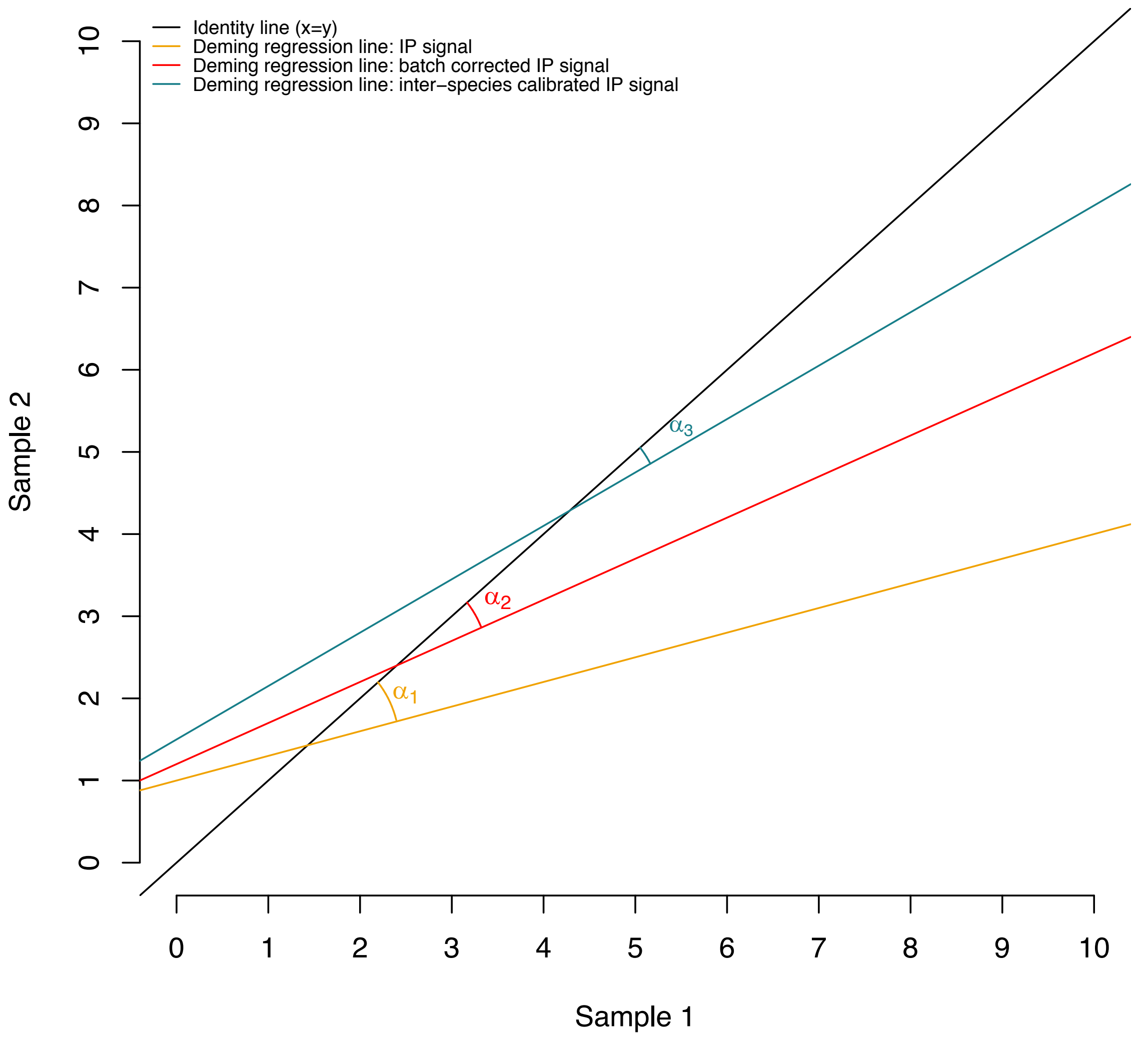


**Supplementary Figure 71. Schematic representation of how the performance of our calibration method was evaluated.** A progressive reduction in the angles to the identity line is expected upon successful normalization of the signa: $\alpha_1 < \alpha_2 < \alpha_3$.

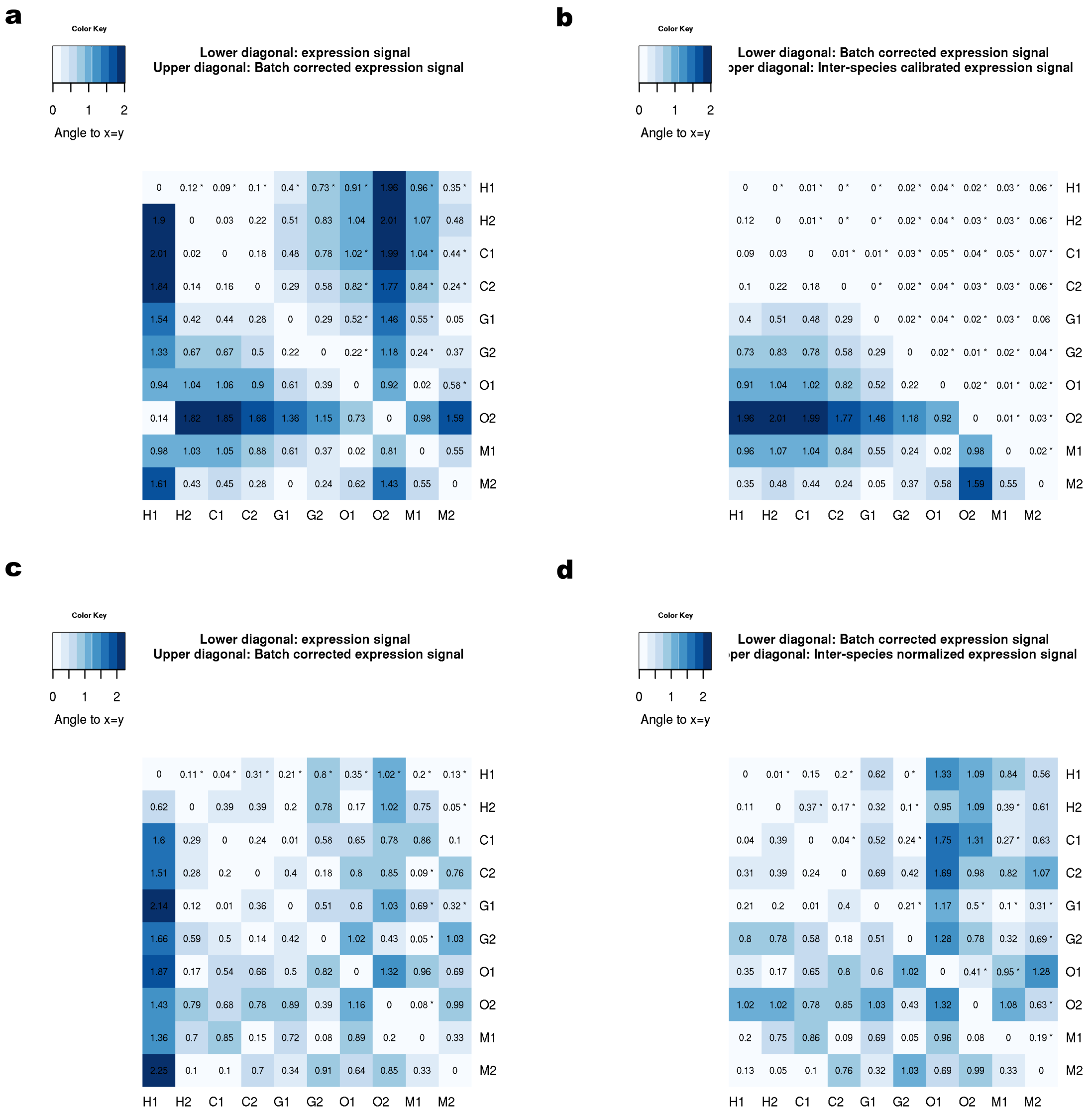

**Supplementary Figure 72. Our signal calibration method effectively reduces noise variability.** Cell values correspond to the angle to identity line (Supplementary Methods, Supplementary Fig. 79) **a,** Batch correction effect on IRCs, **b,** Inter-species calibration effect on IRCs **c,** Batch correction effect on 1-to-1 orthologous protein-coding genes excluding IRCs and **d,** Inter-species calibration effect on IRCs 1-to-1 orthologous protein-coding genes excluding IRCa. **a** and **c,** Lower diagonal cell values correspond to the original angles prior batch correction ($\alpha_1$) whereas upper diagonal correspond to the angle after batch correction ($\alpha_2$). **c** and **d,** Lower diagonal cell values correspond to the angles after batch correction ($\alpha_2$) whereas upper diagonal cell values correspond to the angles after inter-species signal calibration ($\alpha_3$).

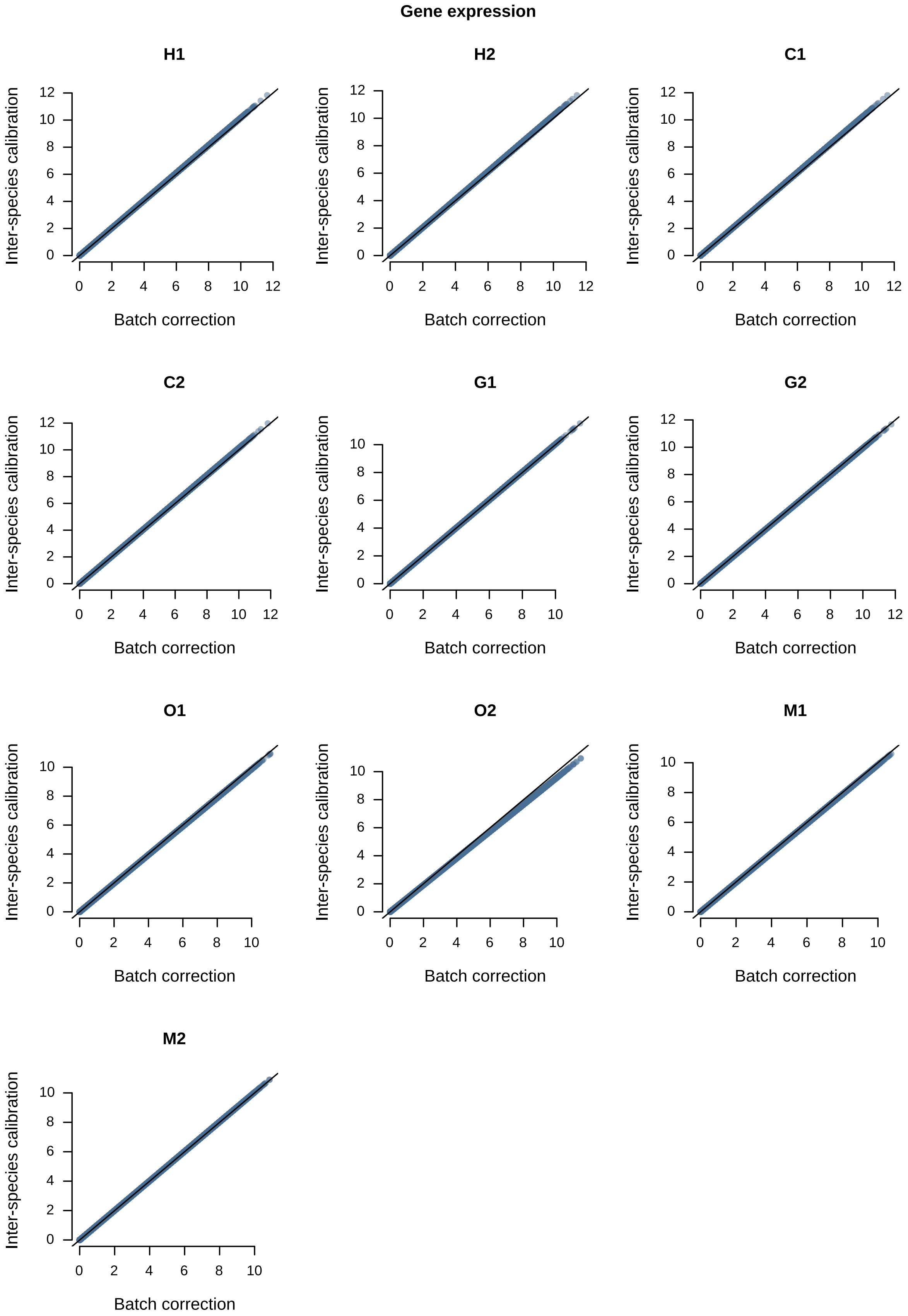


**Supplementary Figure 73. Our signal calibration method effectively normalizes the signals at the tails of the distribution.** Our signal calibration method outperforms quantile normalization (Supplementary Figure 75) since it is able to handle properly the larger inter-species differences at extreme values. For each sample included in this study we plot the batch-corrected signal (x-axis) versus the calibrated signal (y-axis).

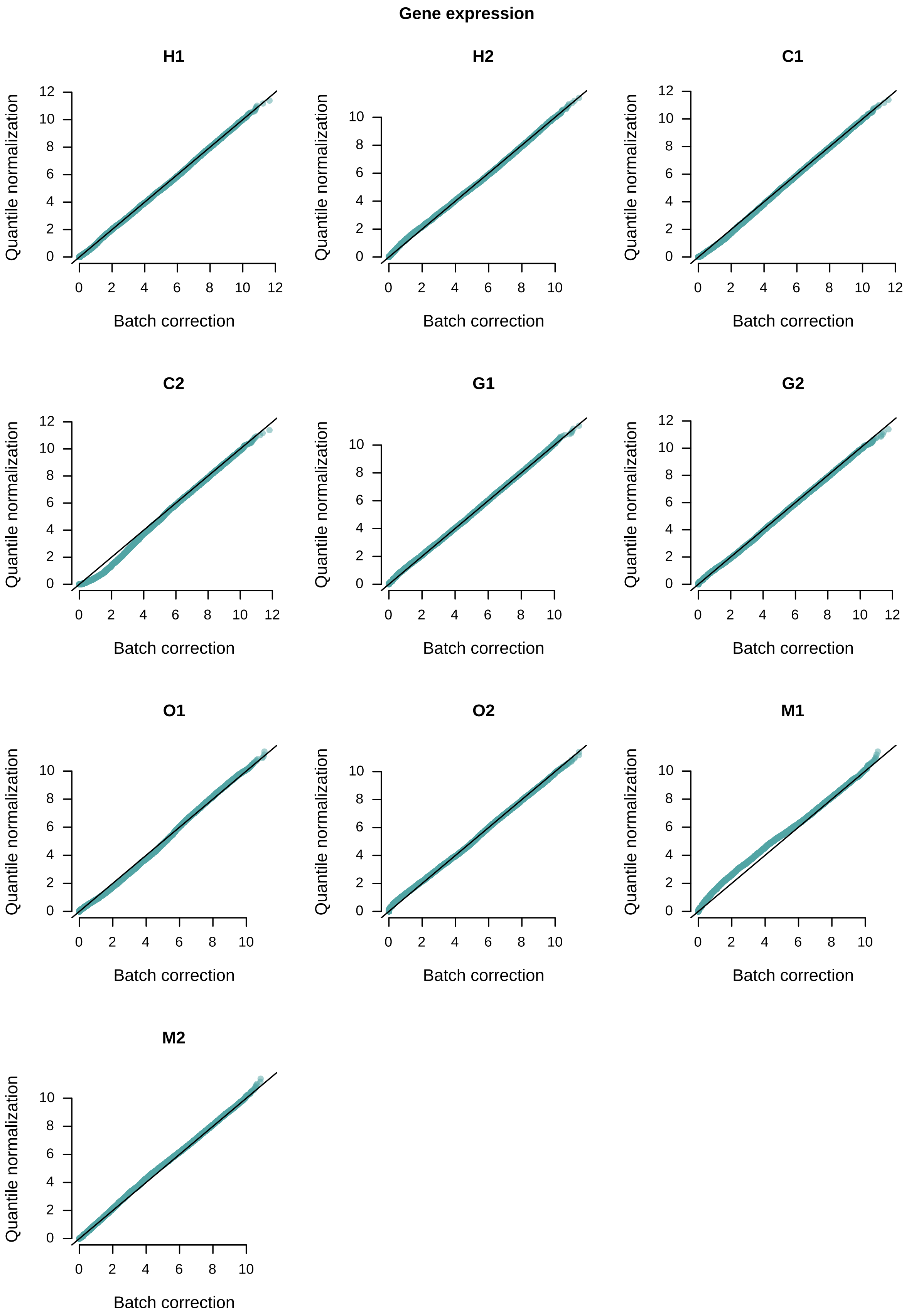

**Supplementary Figure 74. Quantile normalization fails to properly normalize the signals at the tails of the distribution.** For each sample included in this study we plot the batch-corrected signal (x-axis) versus the quantile-normalized signal (y-axis). Note the deviation at the tails.

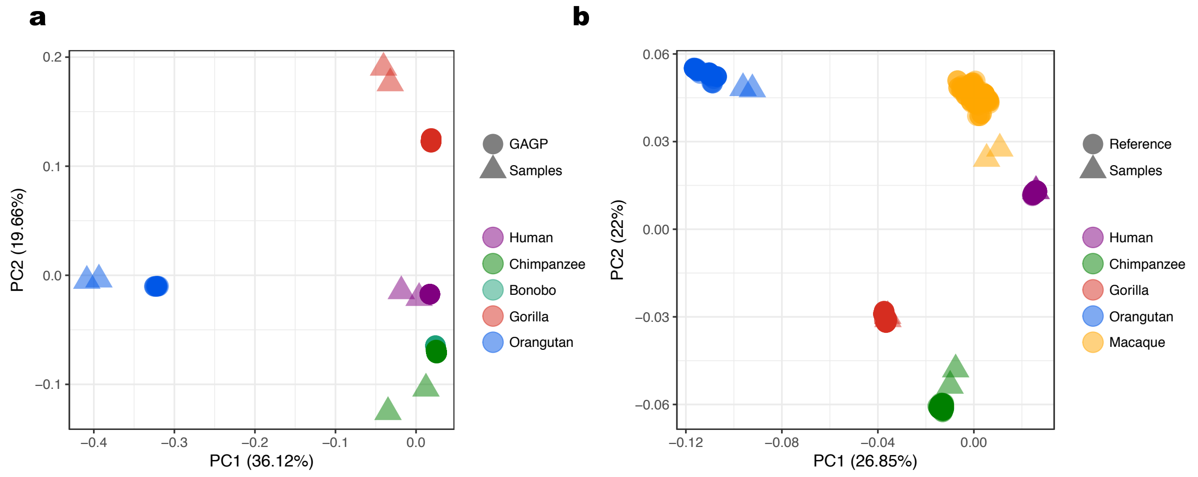


**Supplementary Figure 75. Single nucleotide polymorphisms (SNPs) patterns recapitulate the known phylogenetic relationships between species.** PCA based on **a,** autosomal (chr21) or **b,** mitochondrial SNPs. Triangles represent samples characterized in this study. Circles, samples from different studies (Supplementary Methods).

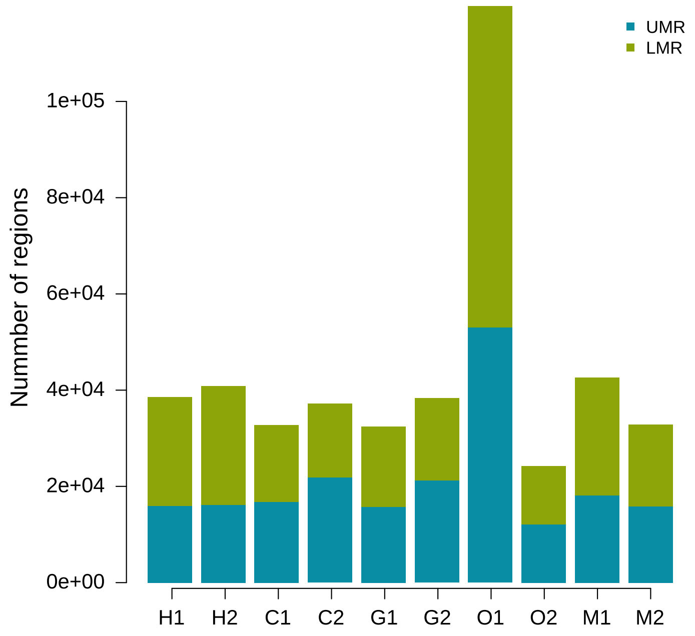


**Supplementary Figure 76. Orangutan sample O1 has a disproportionate number of low methylated regions.** Number of UMRs (unmethylated regions) and LMRs (low methylated regions) annotated per sample (Supplementary Methods).

**Supplementary Table 1.** Annotated regulatory elements for human, chimpanzee, gorilla, orangutan and macaque. Excel files (one excel sheet per species), include the genomic coordinates of the regulatory elements, the assigned epigenetic state in each replicate as well as at the species level, the type of regulatory components and the associated genes.

**Supplementary Table 2.** Number of regulatory elements with promoter and enhancers states annotated in each species.

| | Human | Chimpanzee | Gorilla | Orangutan | Macaque |
|---|---|---|---|---|---|
| **sP** | 7,444 | 7,677 | 8,708 | 8,400 | 7,278 |
| **pP** | 1,103 | 1,156 | 787 | 790 | 930 |
| **wP** | 662 | 2,415 | 2,637 | 2,115 | 698 |
| **aP** | 885 | 941 | 1,153 | 1,414 | 839 |
| **sE** | 21,267 | 29,567 | 25,179 | 20,246 | 29,193 |
| **pE** | 3,078 | 3,518 | 1,719 | 2,313 | 4,892 |
| **wE** | 41,549 | 31,676 | 34,744 | 24,660 | 41,083 |
| **aE** | 18,833 | 9,883 | 12,530 | 15,612 | 11,786 |

**Supplementary Table 3.** Number of regulatory elements classified as each type of regulatory component and number of orphan regulatory elements per species.

| | Human | Chimpanzee | Gorilla | Orangutan | Macaque |
|---|---|---|---|---|---|
| **gP** | 14,191 | 10,932 | 10,171 | 9,399 | 11,365 |
| **gE** | 32,662 | 29,654 | 26,772 | 18,869 | 35,519 |
| **prE** | 6,409 | 5,289 | 5,866 | 3,693 | 6,767 |
| **PiE** | 6,190 | 6,635 | 5,188 | 4,125 | 4,548 |
| **EiE** | 1,437 | 1,969 | 1,733 | 1,033 | 1,208 |
| **Orphan** | 17,662 | 25,204 | 28,747 | 29,021 | 28,427 |

**Supplementary Table 4.** Gene expression levels (TPM) for human, chimpanzee, gorilla, orangutan and macaque. Excel files include the Ensembl gene ID, the gene biotype and the expression level in each replicate.

**Supplementary Table 5.** Number of expressed genes per sample.

| Sample | Expressed genes (TPM>0.5) |
|---|---|
| **H1** | 18,011 |
| **H2** | 19,662 |
| **C1** | 14,525 |
| **C2** | 14,557 |
| **G1** | 14,263 |
| **G2** | 13,308 |

| | |
|---|---|
| **O1** | 14,063 |
| **O2** | 13,308 |
| **M1** | 13,852 |
| **M2** | 14,177 |

**Supplementary Table 6.** Spearman's rho correlation values between TPM expression levels in biological replicates in autosomal protein-coding genes.

| Species | Spearman's rho |
|---|---|
| **Human** | 0.96 |
| **Chimpanzee** | 0.97 |
| **Gorilla** | 0.96 |
| **Orangutan** | 0.92 |
| **Macaque** | 0.97 |

**Supplementary Table 7.** Species pairwise Spearman's rank correlation $\rho$ values in 1-to-1 orthologous protein-coding genes.

| | Human | Chimpanzee | Gorilla | Orangutan | Macaque |
|---|---|---|---|---|---|
| **Human** | 1 | 0.92 | 0.9 | 0.87 | 0.86 |
| **Chimpanzee** | 0.92 | 1 | 0.92 | 0.88 | 0.87 |
| **Gorilla** | 0.9 | 0.92 | 1 | 0.89 | 0.87 |
| **Orangutan** | 0.87 | 0.88 | 0.89 | 1 | 0.84 |
| **Macaque** | 0.86 | 0.87 | 0.87 | 0.84 | 1 |

**Supplementary Table 8.** Number of orthologous regulatory regions with each type of regulatory state.

| | Human | Chimpanzee | Gorilla | Orangutan | Macaque |
|---|---|---|---|---|---|
| **sP** | 2,079 | 2,061 | 2,201 | 2,133 | 1,996 |
| **pP** | 173 | 159 | 103 | 95 | 182 |
| **wP** | 72 | 209 | 324 | 155 | 73 |
| **aP** | 187 | 152 | 208 | 228 | 159 |
| **sE** | 4,905 | 7,193 | 6,172 | 4,919 | 6,229 |
| **pE** | 729 | 742 | 391 | 561 | 1,054 |
| **wE** | 9,879 | 10,040 | 11,685 | 9,360 | 10,166 |
| **aE** | 5,369 | 3,338 | 3,549 | 5,204 | 3,571 |
| **Non-RE** | 5,310 | 4,809 | 4,070 | 6,048 | 5,273 |

**Supplementary Table 9.** Species regulatory state at each orthologous regulatory region, including the type of component assigned to the orthologous regulatory region and the associated gene(s).

**Supplementary Table 10.** Genomic coordinates of orthologous regulatory regions.

**Supplementary Table 11.** Evolutionary conservation of promoters and enhancer states with different activity levels per species in orthologous regulatory regions. Evolutionary conservation is defined as the average number of species in which a particular epigenetic state is conserved.

| | Human | Chimpanzee | Gorilla | Orangutan | Macaque |
|---|---|---|---|---|---|
| **sP** | 4.82 | 4.86 | 4.74 | 4.81 | 4.86 |
| **pP** | 2.9 | 2.86 | 2.51 | 2.84 | 2.26 |
| **wP** | 3.07 | 1.8 | 1.7 | 2.1 | 2 |
| **sE** | 4.31 | 4.1 | 4.09 | 4.3 | 4.01 |
| **pE** | 4.06 | 4.04 | 4.68 | 4.26 | 3.55 |
| **wE** | 3.68 | 3.88 | 3.66 | 3.8 | 3.51 |

**Supplementary Table 12.** Sparse Partial Correlation Networks for the eigencomponents. Excel files include the partial correlation values and the corresponding P-values for the Sparse Partial Correlation Analyses performed for Gene expression and the eigencomponents. Partial correlations and P-values for analyses performed using all 1-to-1 orthologous protein coding genes associated with at least one regulatory element in all species (genes with defined gene regulatory architectures) and with the subset of genes with full regulatory architectures (associated with at least one regulatory element in every type of regulatory component).

**Supplementary Table 13.** Sparse Partial Correlation Networks for the residuals of the eigencomponents for all histone marks. Excel files include the partial correlation values and the corresponding P-values the Sparse Partial Correlation Analyses performed for Gene expression and the residuals of the eigencomponents for H3K4me1, H3K4me3, H3K27ac, H3K27me3 and H3K36me3 together. Partial correlations and P-values for analyses performed using all the genes with regulatory architectures and using genes with a full regulatory architecture are provided.

**Supplementary Table 14.** Sparse Partial Correlation Networks for all histone marks. Excel files include the partial correlation values and the corresponding P-values for the Sparse Partial Correlation Analyses performed for gene expression, H3K4me1, H3K4me3, H3K27ac,

H3K27me3 and H3K36me3 together. Partial correlations and P-values for analyses performed using all 1-to-1 orthologous protein coding genes associated with at least one regulatory element in all species (genes with defined gene regulatory architectures) and with the subset of genes with full regulatory architectures (associated with at least one regulatory element in every type of regulatory component).

**Supplementary Table 15.** Sparse Partial Correlation Networks for histone marks. Excel files, one per histone mark, include the partial correlation values and the corresponding P-values for each of the Sparse Partial Correlation Analysis performed for gene expression and H3K4me1, H3K4me3, H3K27ac, H3K27me3 and H3K36me3. Partial correlations and P-values are included for analyses performed using all 1-to-1 orthologous protein coding genes associated with at least one regulatory element in all species (genes with defined gene regulatory architectures) and with the subset of genes with full regulatory architectures (associated with at least one regulatory element in every type of regulatory component).

**Supplementary Table 16.** Gene expression variability explained by a generalized linear model of gene expression based on H3K27ac, H3K27me3 and H3K36me3 signals at genic promoters with promoter or enhancer states and intragenic enhancers with enhancer states and their interactions (15 variables).

| Histone:Regulatory_component:Epigenetic_State | % explained variance |
|---|---|
| **H3K27ac_P_epiP** | 27.16 |
| **H3K27ac_gE_epiE** | 6.76 |
| **H3K27ac_P_epiE** | 0.48 |
| **H3K27me3_P_epiP** | 1.97 |
| **H3K27me3_gE_epiE** | 10.35 |
| **H3K27me3_P_epiE** | 4.75 |
| **H3K36me3_P_epiP** | 1.44 |
| **H3K36me3_gE_epiE** | 5.1 |
| **H3K36me3_P_epiE** | 0.01 |
| **H3K27ac_P_epiP:H3K27ac_gE_epiE** | 6.52 |
| **H3K27ac_gE_epiE:H3K27ac_P_epiE** | 0.18 |
| **H3K27me3_P_epiP:H3K27me3_gE_epiE** | 0.57 |
| **H3K27me3_gE_epiE:H3K27me3_P_epiE** | 0.44 |
| **H3K36me3_P_epiP:H3K36me3_gE_epiE** | 1.02 |
| **H3K36me3_gE_epiE:H3K36me3_P_epiE** | 0.04 |

**Supplementary Table 17.** Gene expression variability explained by a generalized linear model of gene expression based on H3K4me3, H3K4me1, H3K27ac, H3K27me3 and H3K36me3 signals at all types of regulatory components, including both promoter and enhancer states for each component as well as all possible interaction terms (1,225 variables). Excel file.

**Supplementary Table 18.** GTEx tissue group correspondence.

| Tissue | Tissue Group |
|---|---|
| **Adipose - Visceral (Omentum)** | AdiposeTissue |
| **Adipose - Subcutaneous** | AdiposeTissue |
| **Adrenal Gland** | AdrenalGland |
| **Whole Blood** | Blood |
| **Artery - Coronary** | BloodVessel |
| **Artery - Aorta** | BloodVessel |
| **Artery - Tibial** | BloodVessel |
| **Brain - Substantia nigra** | Brain |
| **Brain - Spinal cord (cervical c-1)** | Brain |
| **Brain - Amygdala** | Brain |
| **Brain - Anterior cingulate cortex (BA24)** | Brain |
| **Brain - Hippocampus** | Brain |
| **Brain - Hypothalamus** | Brain |
| **Brain - Putamen (basal ganglia)** | Brain |
| **Brain - Cerebellar Hemisphere** | Brain |
| **Brain - Frontal Cortex (BA9)** | Brain |
| **Brain - Caudate (basal ganglia)** | Brain |
| **Brain - Nucleus accumbens (basal ganglia)** | Brain |
| **Brain - Cortex** | Brain |
| **Brain - Cerebellum** | Brain |
| **Breast - Mammary Tissue** | Breast |
| **Colon - Sigmoid** | Colon |
| **Colon - Transverse** | Colon |
| **Esophagus - Gastroesophageal Junction** | Esophagus |
| **Esophagus - Muscularis** | Esophagus |
| **Esophagus - Mucosa** | Esophagus |
| **Heart - Atrial Appendage** | Heart |
| **Heart - Left Ventricle** | Heart |
| **Kidney - Cortex** | Kidney |
| **Liver** | Liver |
| **Lung** | Lung |
| **Muscle - Skeletal** | Muscle |
| **Nerve - Tibial** | Nerve |
| **Ovary** | Ovary |
| **Pancreas** | Pancreas |
| **Pituitary** | Pituitary |
| **Prostate** | Prostate |

| | |
|---|---|
| **Minor Salivary Gland** | SalivaryGland |
| **Skin - Not Sun Exposed (Suprapubic)** | Skin |
| **Skin - Sun Exposed (Lower leg)** | Skin |
| **Small Intestine - Terminal Ileum** | SmallIntestine |
| **Spleen** | Spleen |
| **Stomach** | Stomach |
| **Testis** | Testis |
| **Thyroid** | Thyroid |
| **Uterus** | Uterus |
| **Vagina** | Vagina |
| **LCLs** | LCLs |
| **Fibroblasts** | Fibroblasts |

**Supplementary Table 19.** GO term clustering with group labels used for representation.

**Supplementary Table 20.** Functional enrichment of genes associated with genic promoters (gP) with conserved strong promoter states (sP) compared to genes with associated genic promoters.

**Supplementary Table 21.** Functional enrichment of genes associated with intragenic enhancers (gE) with conserved strong enhancer states (sE) compared to genes with associated intragenic enhancers.

**Supplementary Table 22.** Functional enrichment of genes associated with proximal enhancers (prE) with conserved poised enhancer states (pE) compared to genes with associated proximal enhancers.

**Supplementary Table 23.** Functional enrichment of genes associated with genic promoters (gP) with conserved poised enhancer states (pE) compared to genes with associated genic promoters.

**Supplementary Table 24.** Functional enrichment of genes associated with intragenic enhancers (gE) with conserved weak enhancer states (wE) compared to genes with associated intragenic enhancers.

**Supplementary Table 25.** Functional enrichment of genes associated with human-specific intragenic enhancers (gE) with weak enhancer states (wE) compared to genes associated with intragenic enhancers.

**Supplementary Table 26.** Effect sizes of significant pairwise comparisons (Wilcoxon-Nemenyi-McDonald-Thompson test, $P < 0.05$) of GTEx tissue median expression values within datasets of genes with conserved and human-specific regulatory elements.

**Supplementary Table 27.** Genomic coordinates of human-specific gains of intragenic enhancers with weak states and further characterization.

**Additional File 1.** Genomic coordinates of open chromatin regions detected in each species. Excel files, one per species.

**Additional File 2.** Genotyping statistics.

| | Mean coverage | X to mean coverage | # SNPs | # Indels | # STRs | Chr21 | Mitochondrial |
|---|---|---|---|---|---|---|---|
| H1 | 38.08 | 1 | 3,575,007 | 891,779 | 304,942 | Eurasian | - |
| H2 | 33.93 | 0.45 | 4,340,954 | 1,063,458 | 489,526 | African | - |
| C1 | 19.94 | 0.47 | 8,412,644 | 1,782,217 | 715,246 | *Pan troglodytes troglodytes* | *Pan troglodytes troglodytes* |
| C2 | 14.27 | 0.46 | 3,185,759 | 884,459 | 481,686 | *Pan troglodytes verus* | *Pan troglodytes verus* |
| G1 | 18.58 | 0.94 | 7,790,413 | 1,422,292 | 551,236 | *Gorilla gorilla gorilla* | *Gorilla gorilla gorilla* |
| G2 | 17.39 | 0.53 | 7,640,735 | 1,391,387 | 532,458 | *Gorilla gorilla gorilla* | *Gorilla gorilla gorilla* |
| O1 | 21.89 | 1.1 | 11,221,590 | 2,825,670 | 624,672 | *Pongo pygmaeus* | *Pongo pygmaeus* |
| O2 | 15.01 | 0.6 | 11,545,091 | 2,813,677 | 623,89 | *Pongo pygmaeus* | *Pongo pygmaeus* |
| M1 | 13.97 | 0.5 | 10,668,361 | 2,495,486 | 691,368 | Closer to Chinese | - |
| M2 | 14.83 | 01.09 | 11,113,027 | 2,624,693 | 764,158 | Closer to Chinese | - |

**Additional File 3.** Mitochondrial reference sequences.

| Species | Citation | Link |
|---|---|---|
| Human | Ignman and Gyllensten, 2006 | https://www.ncbi.nlm.nih.gov/pmc/articles/PMC1347373/ |
| Chimpanzee | Lobon et al., 2016 | https://academic.oup.com/gbe/article/8/6/2020/2574100 |
| Bonobo | Lobon et al., 2016 | https://academic.oup.com/gbe/article/8/6/2020/2574100 |
| Gorilla | Xue et al., 2015 | http://science.sciencemag.org/content/348/6231/242 |
| Orangutan | Nater et al., 2017 | https://www.cell.com/current-biology/fulltext/S0960-9822(17)31245-9 |
| Macaque | Matsudaira et al., 2017 | https://academic.oup.com/jhered/article/109/4/360/4653541 |
| Macaque | Liedigk et al., 2015 | https://bmcgenomics.biomedcentral.com/articles/10.1186/s12864-015-1437-0 |
| Macaque | Chen et al., 2016 | https://www.tandfonline.com/doi/abs/10.3109/19401736.2015.1025265 |

**Additional File 4.** Genomic coordinates of unmethylated (UMR) and low methylated regions (LMR). Excel files, one per species.

**Additional File 5.** VCF files with the SNPs, indels and STRs genotyped in each sample. Visit http://biologiaevolutiva.org/tmarques/data/
